# Systematic Profiling of Essential Fungal Transcriptional Regulators Uncovers Ezt1 as a Central Pathobiological and Morphogenic Regulator in *Cryptococcus neoformans*

**DOI:** 10.64898/2026.04.30.722081

**Authors:** Seung-Heon Lee, Sheng Sun, Yu-Byeong Jang, Seong-Ryong Yu, Yeseul Choi, Jin-Tae Choi, Eui-Seong Kim, Jong-Seung Lee, Leah E. Cowen, Joseph Heitman, Kyung-Tae Lee, Yong-Sun Bahn

**Author notes:** **Correspondence and requests for materials should be addressed to JH, KTL, and YSB**.

## Abstract

*Cryptococcus neoformans* is a global fungal pathogen that causes fatal cryptococcosis, and the limitations of current antifungals underscore the urgent need for new therapeutics. Here we systematically investigate essential transcriptional regulators in *C. neoformans* as potential antifungal targets, developing experimental pipelines that assess growth requirement, essentiality and function through conditional gene expression, constitutive overexpression, and meiotic spore analysis. We identify one quasi-essential (growth-required but non-essential) transcription factor, Fhl1, and 13 essential transcriptional regulators, three of which are transcription factors (Ezt1, Ezt2 and Cbf1) highly divergent from counterparts in other eukaryotes. Notably, Ezt1 modulates the expression of more than 1,200 genes, controlling growth, antifungal drug and stress responses, sexual development and virulence. Our findings define the essential transcriptional regulator landscape of *C. neoformans* and provide a framework for prioritising divergent essential regulators, particularly Ezt1, for antifungal target discovery.

## Introduction

*Cryptococcus neoformans*, a basidiomycetous fungus within the pathogenic *Cryptococcus* species complex, is responsible for fatal cryptococcosis in humans^1^. Found in diverse natural environments such as soils and avian excreta, *C. neoformans* primarily affects immunocompromised individuals^2^. Infection commences with the inhalation of spores, which can subsequently disseminate to various organs. Remarkably, the fungus can evade or proliferate within alveolar macrophages^3^ and exhibit neurotropism, allowing it to breach the blood-brain barrier (BBB) and cause lethal meningoencephalitis^4,5^. *C. neoformans* accounts for 19% of AIDS-related deaths, with an estimated annual death toll of 112,000^6^.

Cryptococcosis treatment is hindered by the limited availability of antifungal drugs, further complicated by the limited permeability of the BBB. Furthermore, the prolonged use of antifungal medications, coupled with undesirable side effects and limited efficacy, has led to the emergence of drug-resistant strains, underscoring the urgent need for novel anticryptococcal therapies with unique mechanisms of action. Although targeting essential fungal proteins has proven effective in antifungal drug development, it presents challenges due to potential toxicity risks: many of these proteins are conserved between fungi and humans. For example, the essential TOR kinase Tor1 can be inhibited by rapamycin, leading to significant antifungal activity^7–9^. However, its associated cellular toxicity in humans^10,11^ prevents the use of its inhibitors as a viable antifungal strategy. Thus, developing safer antifungal drugs may require targeting evolutionarily less conserved, species-specific essential proteins.

Essential transcription factors (TFs) have been proposed as promising drug targets due to their evolutionary divergence from other essential signalling and metabolic proteins^12–14^. However, developing TF-targeting drugs poses challenges, as they must disrupt protein-DNA or protein-protein interactions. Despite these difficulties, there have been successful efforts in cancer treatment and other fields^15–17^. Several essential TFs have been identified in fungi, such as heat shock factor 1 (Hsf1), which regulates thermotolerance and heat shock protein expression in various fungal species, including *C. neoformans*, *Saccharomyces cerevisiae*, and *Candida albicans*^18,19^. In humans, HSF1 functions as a master regulator of heat shock protein expression under thermal stress and plays a role in various aspects of cancer^20^. Other essential TFs have been identified in *S. cerevisiae* and *C. albicans*^21–23^.

Our previous systematic analysis of 178 TFs in *C. neoformans* identified 23 likely essential transcriptional regulators, as these genes could not be deleted despite repeated attempts^12^. Among these, Hsf1 was confirmed to be essential by our prior study^24^. In contrast, five TFs, namely Mig1, Cys3, Cir1, Ecm2201, and Ccd6, were previously identified as quasi-essential (defined here as dispensable for viability but required for robust vegetative growth or colony formation^25^) or non-essential and partially characterised^26–30^. Therefore, this study systematically examined the remaining 17 putative essential transcriptional regulators using conditional gene expression, constitutive overexpression, and spore analysis of heterozygous mutants in engineered diploid strains. Our approach confirmed the essentiality of 13 transcriptional regulators and elucidated their pathobiological roles both in vitro and in vivo. Notably, Ezt1 emerged as a key transcriptional regulator modulating the expression of more than 1,200 genes, many of which are involved in the pathobiology and reproduction of *C. neoformans*. This research functionally characterised an unprecedented number of essential transcriptional regulators in the global fungal meningitis pathogen, providing a foundation for future studies of antifungal vulnerabilities in *C. neoformans*.

## Results

### Strategy for the identification and functional analysis of essential transcription regulators

In our study, we focused on 17 previously uncharacterised essential putative transcriptional regulators (Supplementary Data 1). For those already annotated in FungiDB or assigned by homology to *S. cerevisiae*, names and rationales are listed in Supplementary Data 1. For the TFs we newly named, we used domain-based or functional classification: we named *HLH7* (<u>h</u>elix-loop-<u>h</u>elix type <u>7</u>) and *FZC52* (fungal <u>Z</u>n_2_-<u>C</u>ys_6_ type <u>52</u>) based on their TF domain family classification, as no clear yeast orthologues could be identified. Lastly, *EAT1* (<u>e</u>ssential <u>A</u>PSES <u>T</u>F 1), *EZT1* and *EZT2* (<u>e</u>ssential <u>z</u>inc-finger <u>T</u>Fs 1 and 2) were named due to their classification as essential TFs, as elucidated later in this paper.

We employed two complementary strategies to assess the essentiality of the transcriptional regulators (Fig. 1). First, we generated conditional gene expression strains for each gene by replacing its native promoters with the copper-repressible *CTR4* (copper transporter gene 4) promoter^31^. Transcriptional regulators exhibiting impaired growth in repressive conditions were classified as ‘growth-required’, while those without growth defects were subjected to further gene deletion attempts. While the *CTR4* promoter replacement approach only indicates whether a target gene is required for growth, it does not confirm the essentiality of the genes. To resolve this, we constructed heterozygous mutants for spore analysis of all essential transcriptional regulator candidates, following a reported method based on engineered diploid platform strains CnLC6683 (KN99 *MAT*α/*MAT***a**)^32^ and AI187 (*ADE2*/*ade2 URA5*/*ura5 MAT*α/*MAT***a**)^33^. The next steps involved whole-genome sequencing (WGS) to avoid misinterpretations arising from unintended genetic anomalies, such as aneuploidy, chromosomal rearrangements and segmental duplication or deletions.

**FIG 1.**
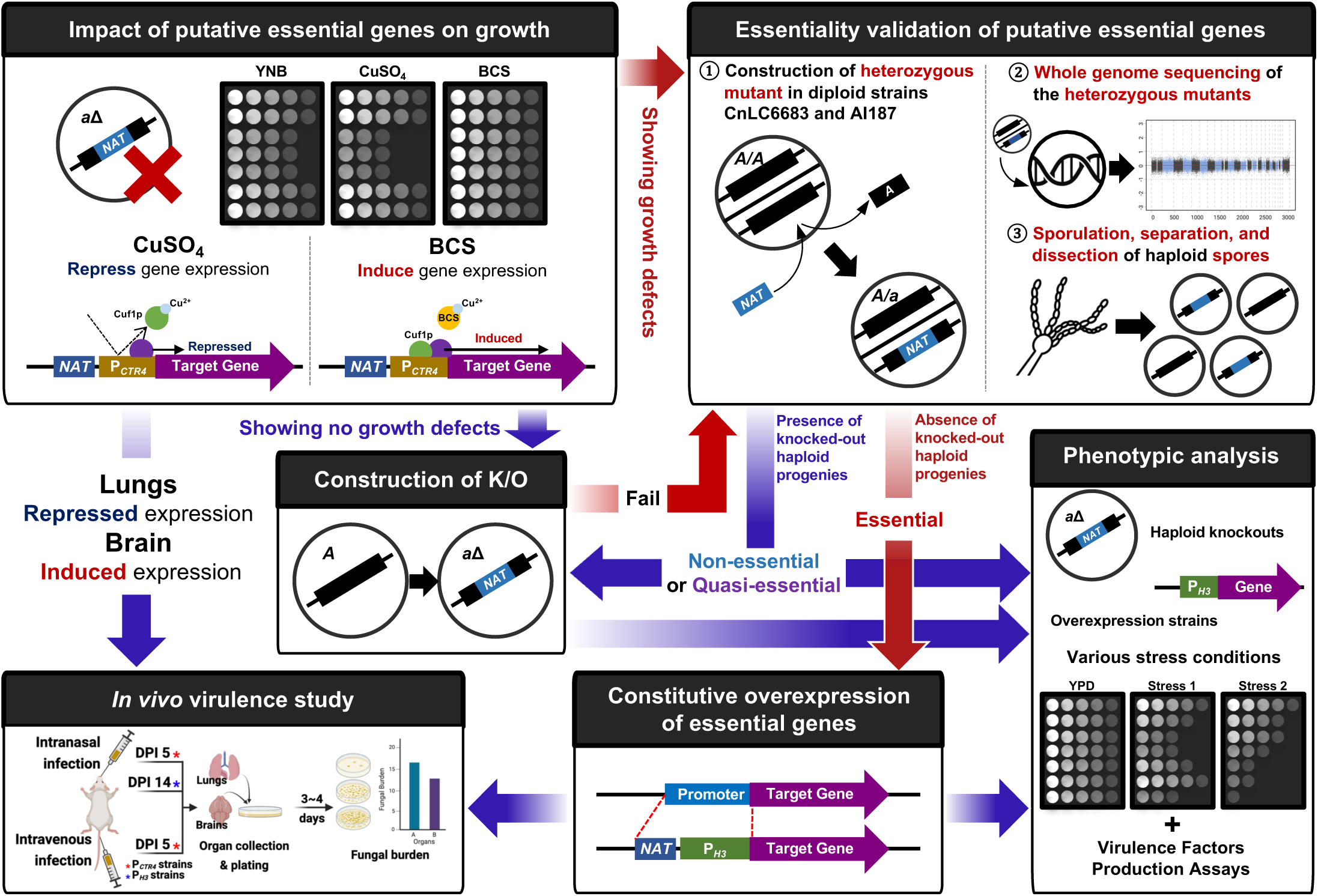
Schematic representation of the strategy used for assessing cryptococcal essential transcriptional regulators. Essential transcriptional regulators were identified using both *CTR4* promoter replacement and meiotic spore analysis of heterozygous mutants. Their functional roles were further examined through in vitro phenotypic analysis of constitutive overexpression strains and deletion mutants, as well as in vivo virulence assays using *CTR4* promoter-driven conditional expression and *H3* promoter-driven constitutive overexpression strains in murine intranasal and intravenous infection models. The images illustrating the in vivo virulence study and whole-genome sequencing were created with BioRender.

For WGS-validated heterozygous mutants, we then used two complementary spore isolation methods—spore dissection and spore enrichment—together with genotypic analysis to identify haploid progeny. Target genes were classified as essential genes if no viable spores carrying the disrupted allele were recovered (Fig. 1). To minimise the effect of background mutations, we primarily analysed heterozygous mutants in CnLC6683, a marker-free strain derived from fusion of KN99α and KN99**a**^32^, by spore dissection directly from basidiospore chains, followed by genotyping. To analyse larger numbers of spores than would be feasible by dissection, and genes for which genetically validated CnLC6683-based heterozygous mutants could not be obtained, we additionally performed spore enrichment method for AI187-derived heterozygous mutants using Percoll^®^ density-gradient centrifugation^34^. Haploids were isolated using auxotrophic markers and then genotyped by PCR using primers internal to the deleted gene, as described in Methods. Together, these two approaches enabled a more precise and comprehensive assessment of transcriptional regulator essentiality.

Because viable haploid null mutants cannot be recovered for essential genes, we developed constitutive overexpression strains utilising the histone *H3* promoter to analyse the function of these putative essential transcriptional regulators. Subsequently, we assessed their phenotypic traits under 28 distinct in vitro growth conditions. For in vivo phenotypic analysis, we employed fungal burden assays in murine intranasal and intravenous infection models, utilising wild-type, constitutive overexpression, and *CTR4* promoter replacement strains (Fig. 1), as previously documented^35^.

### Uncovering essential transcriptional regulators in *C. neoformans*

Among the 17 putative essential transcriptional regulators, analysis of the constructed *CTR4* promoter replacement strains (Supplementary Fig. 1) showed that six genes (*CEF1*, *EAT1*, *SNU66*, *CDC39*, *ZAP101* and *EZT2*) are strongly required for growth, whereas eight genes (*CBF1*, *RSC8*, *ESA1*, *EZT1*, *SGT1*, *PZF1*, *FHL1* and *TOP3*) are weakly required for growth (Fig. 2a and 2b). The remaining three TFs (*ASR2*, *HLH7*, and *FZC52*) exhibited no apparent growth defect in the presence of CuSO_4_ (Fig. 2c and 2d).

**FIG 2.**
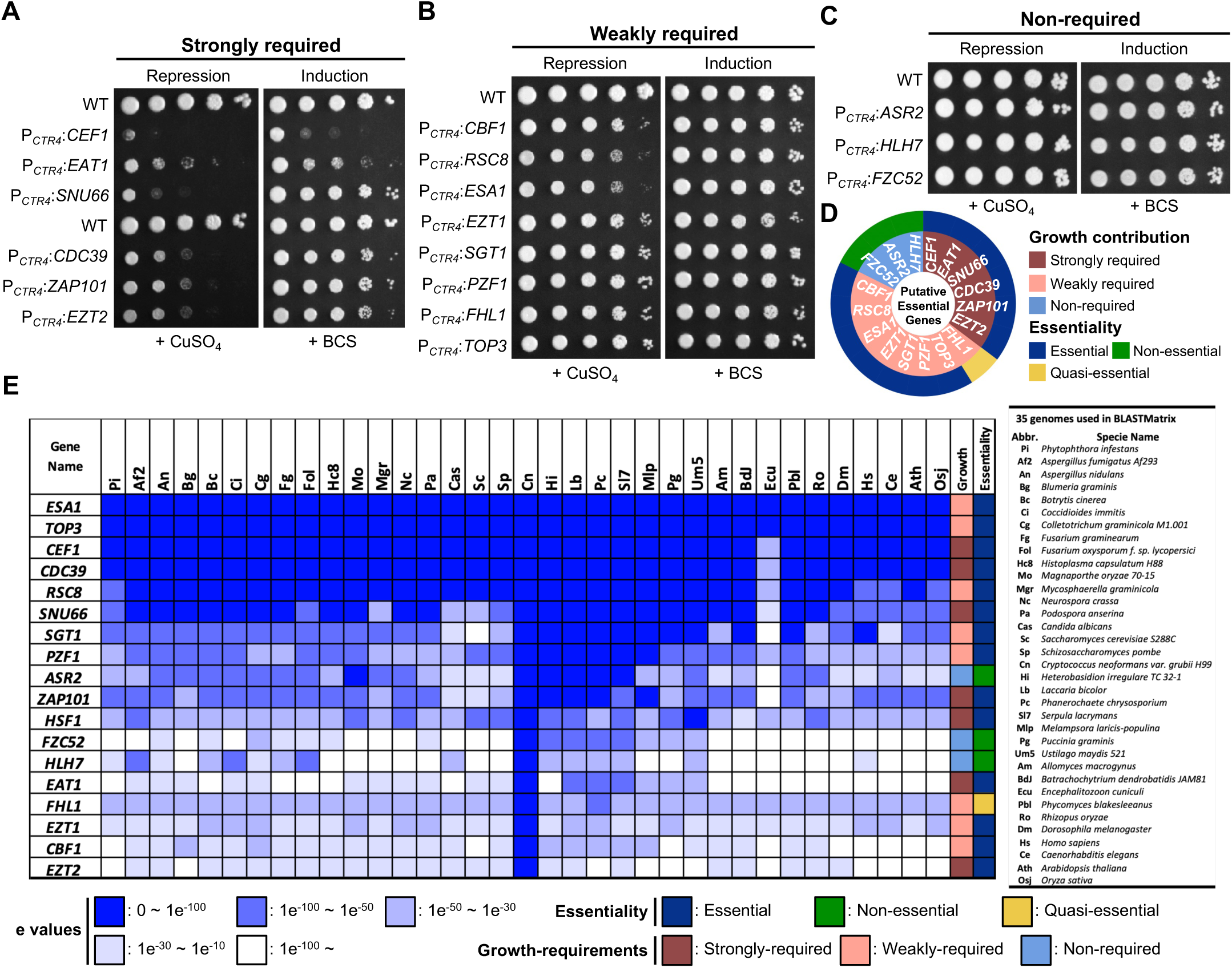
Essentiality assessment of 17 putative essential regulators in *C. neoformans*. (A-C) The wild-type (WT; H99L) and *CTR4* promoter replacement strains were cultured overnight at 30 in liquid YPD medium, serially diluted (1 to 10^4^), and spotted onto YPD plates containing 25 μM CuSO_4_ for repression or 200 μM BCS for induction of the putative essential gene. The plates were incubated at 30 for 3 days and photographed. *CTR4* promoter replacement strains of *CEF1*, *EAT1*, *SNU66*, *CDC39*, *ZAP101*, and *EZT2* exhibited notable growth defects when CuSO_4_ was added, which were partially rescued when BCS was added to the medium, indicating that these genes are strongly required for the growth of *C. neoformans*. (B) Using the same experimental method as in (A), eight transcriptional regulators (*CBF1*, *RSC8*, *ESA1*, *EZT1*, *SGT1*, *PZF1*, *FHL1*, and *TOP3*) were classified as weakly required, and three TFs (*ASR2*, *HLH7*, and *FZC52*) (C) showed no detectable growth requirement under the tested repression conditions. (D) Spore analysis of heterozygous mutants confirmed that 13 regulators are essential for *C. neoformans* viability, with *FHL1* classified as quasi-essential because it is required for normal growth but dispensable for viability. (E) A BLAST matrix of the putative essential transcriptional regulators was generated using the Comparative Fungal Genomics Platform (http://cfgp.riceblast.snu.ac.kr). Abbreviations (Abbr.) are indicated next to the matrix.

We next generated heterozygous mutants in engineered diploid platform strains (CnLC6683 and AI187) for all 17 transcriptional regulators for subsequent spore analysis (Supplementary Figs. 2, 3, and 4). For most targets, we obtained at least one heterozygous isolate from each diploid background whose WGS profile confirmed loss of a single allele, with no additional truncations or copy-number alterations in the genomic region surrounding the target gene. However, for two genes (*CBF1* and *TOP3*), WGS-validated intact isolates were obtained only in the AI187 background, whereas for two others (*CEF1* and *FZC52*) such isolates were obtained only in the CnLC6683 background. From 56 heterozygous mutants, we isolated and analysed 1,543 spores: 720 spores from CnLC6683-based spore dissection and 823 spores from AI187-based spore dissection or enrichment (examples in Supplementary Fig. 5). These analyses indicated that 13 transcriptional regulators are essential for viability, with the exception of *HLH7*, *ASR2*, *FHL1* and *FZC52*. Haploid *NAT-*positive progeny carrying only the deletion alleles of the genes were recovered from selfing of *HLH7/hlh7*, *ASR2*/*asr2*, *FHL1*/*fhl1,* and *FZC52*/*fzc52* mutants, confirming that these genes are not essential. For the other 13 candidate essential transcriptional regulators, while a small number of *NAT*-positive meiotic progeny were obtained from each of them, all of them appeared to be aneuploid, containing both deletion and the wild-type alleles of the target gene. Thus, these 13 putative essential transcriptional regulators are indeed essential in *C. neoformans*. It should be noted that whereas *FHL1* is required for normal growth, it is dispensable for viability, consistent with classification as a quasi-essential gene (Fig. 2d and Supplementary Data 1).

Homology searches using BLAST (Basic Local Alignment Search Tool), together with published essentiality annotations, indicated that except for the seven proteins (Cbf1, Zap101, Eat1, Ezt1, Ezt2, Hlh7, and Fzc52), the remaining transcriptional regulators have orthologues reported to be essential in at least one of the following model fungi: *S. cerevisiae*, *Candida albicans*, *Nakaseomyces glabratus* and *Schizosaccharomyces pombe*^36–41^ (Supplementary Data 1). Notably, the six transcriptional regulators with the highest sequence homology (Esa1, Top3, Cef1, Cdc39, Rsc8, and Snu66) were essential and comparatively conserved across diverse eukaryotes (Fig. 2e). According to their functional annotations, these evolutionarily conserved essential regulators are primarily components of fundamental cellular machinery. Specifically, Cef1 (a pre-mRNA splicing factor) and Snu66 (a U4/U6.U5 tri-snRNP-associated protein 1) are critical components of the spliceosome complex, rendering them indispensable for proper RNA processing^42^. Additionally, Cdc39 (a CCR4-NOT transcription complex subunit), Rsc8 (an RSC chromatin remodelling complex subunit), and Esa1 (a histone acetyltransferase) serve as key regulators of transcriptional control and chromatin remodelling^43–45^. Furthermore, Top3 functions as a DNA topoisomerase III, essential for maintaining DNA topology^37^, and Sgt1 is involved in kinetochore assembly and faithful chromosome segregation^46^. By contrast, the three most divergent TFs—Cbf1, Ezt1 and Ezt2—remained essential despite substantial sequence divergence (Fig. 2e). Based on these functional annotations, the identified transcriptional regulators were classified into four broad functional categories: putative TFs (Eat1, Ezt2, Zap101, Pzf1, Ezt1, Fhl1, Cbf1, Hlh7, Asr2, and Fzc52), RNA-processing-associated factors (Cef1 and Snu66), a transcriptional co-regulator Cdc39, and chromosome/chromatin-associated regulators (Rsc8, Esa1, Top3, and Sgt1) (summarised in Supplementary Data 1).

### In vitro and in vivo phenotypic profiling of essential cryptococcal transcriptional regulators

To functionally characterise the essential transcriptional regulators, we constructed constitutive overexpression strains driven by the histone H3 promoter (Supplementary Fig. 6) in the H99L *MAT*α (a hypervirulent sub-lineage of the type strain H99^47^). As *FZC52* was determined to be non-essential by meiotic spore analysis and H99L *MAT*α-background *fzc52*Δ mutants were available, *FZC52* was analysed using the knockout strains rather than overexpression strains. For the other three non-essential TFs, phenotypes were assessed using haploid knockout progeny obtained by spore dissection from *HLH7/hlh7*, *ASR2/asr2* and *FHL1/fhl1* heterozygous mutants in the CnLC6683 background (Supplementary Fig. 7). The *ASR2* and *HLH7* overexpression strains, by contrast, had been generated and characterised at an earlier stage, when both genes were still predicted to be essential, and were retained in the phenotypic profiling. Using the available overexpression and knockout strains, we performed a comprehensive in vitro phenotypic analysis to elucidate regulator function in *C. neoformans* (Fig. 3a).

**FIG 3.**
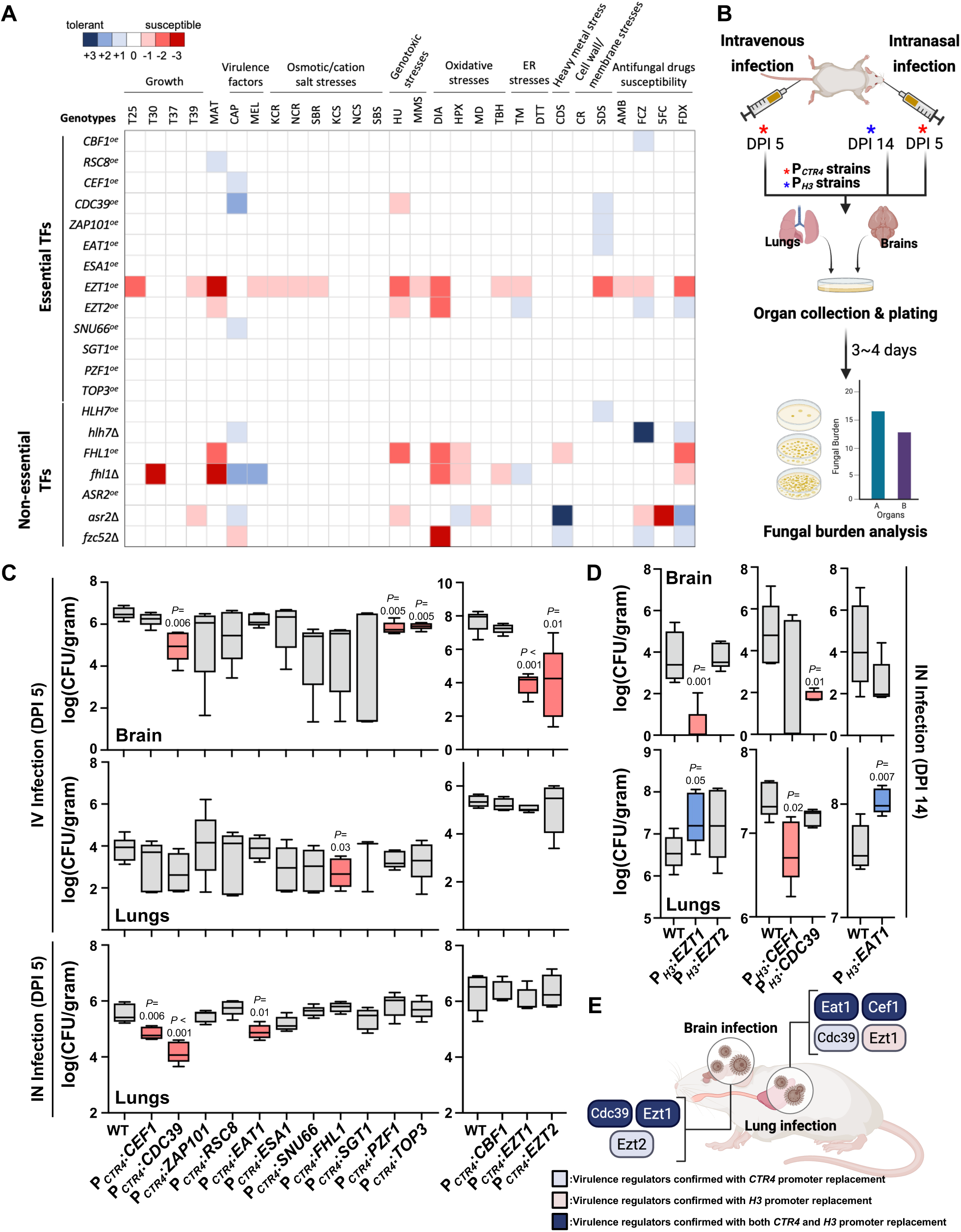
Phenotypic and pathogenicity profiling of essential transcriptional regulators in *C. neoformans*. (A) In vitro phenotypic traits of essential gene overexpression strains and non-essential gene knockout/overexpression strains (with *FZC52* assessed using knockout strains only) were assessed under various growth environments, stress conditions, antifungal treatments, and virulence-factor-inducing conditions. In the heat map, colours represent ordinal phenotypic scores relative to the wild-type strain, ranging from -3 to +3; red denotes negative scores whereas blue denotes positive scores, with colour intensity reflecting differences of approximately one, two, or three or more 10-fold dilutions; for capsule, melanin, and mating phenotypes, scores reflect the corresponding magnitude of phenotypic change, as defined in Methods. For growth, stress, and antifungal assays, red indicates reduced growth or increased susceptibility, whereas blue indicates enhanced growth or increased tolerance. For virulence-factor and developmental assays, red indicates reduced capsule production, melanisation, or mating, whereas blue indicates the corresponding increase. Abbreviations: T25 (25), T30 (30), T37 (37), T39 (39), MAT (mating), CAP (capsule production), MEL (melanin production), KCR (YPD + 1.5 M KCl), NCR (YPD + 1.5 M NaCl), SBR (YPD + 2 M sorbitol), KCS (YP + 1 M KCl), NCS (YP + 1 M NaCl), SBS (YP + 2 M sorbitol), HU (hydroxyurea), MMS (methyl methanesulphonate), DIA (diamide), HPX (hydrogen peroxide), MD (menadione), TBH (*tert*-butyl hydroperoxide), TM (tunicamycin), DTT (dithiothreitol), CDS (cadmium sulphate), CR (Congo red), SDS (sodium dodecyl sulphate), AMB (amphotericin B), FCZ (fluconazole), 5FC (5-flucytosine), FDX (fludioxonil). (B) Schematic representation of fungal burden assays in murine intranasal (IN) and intravenous (IV) infection models. *CTR4* promoter replacement strains were used to assess fungal burden following both IV and IN infections. The *H3* promoter overexpression strains were tested following IN infection to evaluate the impact of transcriptional regulation on fungal burden. This image was created with BioRender. (C) Fungal burden analysis of *CTR4* promoter replacement strains in murine lungs and brains after IV and IN infections, compared with the parental wild-type strain (WT; H99L). Box plots indicate fungal CFU levels for each strain. Genes influencing brain and lung burdens are highlighted in red. Statistical significance of difference was determined by two-tailed Welch’s *t*-tests using Prism 11.0 and *P* values are indicated above or below the box plot. (D) Fungal burden in lungs and brain of overexpression strains for *EZT1*, *EZT2*, *CEF1*, *CDC39*, and *EAT1* following IN infection. (E) Overview diagram illustrating the essential regulators affecting brain and lung infections by *C. neoformans*. Essential regulators such as *EZT1* and *CDC39* were associated with both brain and lung infection, while *EAT1* and *CEF1* played roles in lung infection. This image was created with BioRender.

We tested 28 conditions spanning temperature sensitivity, mating efficiency, virulence factor production (capsule and melanin), stress responses and antifungal drug susceptibility (Fig. 3a and Supplementary Fig. 8). While most overexpression strains showed minor or no phenotypic changes, *EZT1oe* and *EZT2oe* strains exhibited more pronounced phenotypes (Fig. 3a). Notably, *EZT1* overexpression markedly impaired growth at 25 and 39 and increased susceptibility to osmotic, genotoxic and oxidative stresses (diamide and organic peroxide), tunicamycin, SDS and antifungal agents (amphotericin B, fluconazole and fludioxonil). By contrast, *EZT2* overexpression increased susceptibility to hydroxyurea and diamide, while enhancing resistance to tunicamycin and antifungal agents (fluconazole and fludioxonil).

*FHL1*, a quasi-essential gene, conferred dynamic phenotypic effects in both the overexpression and knockout contexts. The *FHL1oe* strain showed increased susceptibility to hydroxyurea and heavy metal CdSO_4_. In contrast, the *fhl1*Δ mutant exhibited enhanced melanin and capsule production, increased susceptibility to organic peroxide, and heightened resistance to tunicamycin (Fig. 3a). Notably, both *FHL1oe* and *fhl1*Δ strains displayed impaired growth under oxidative stress (diamide and H_2_O_2_) and reduced mating efficiency. These results underscore the importance of tightly balanced *FHL1* expression for adaptation to diverse stresses and for sexual development.

To elucidate the roles of quasi-essential and essential transcriptional regulators in the pathogenicity of *C. neoformans*, we performed fungal burden assays in murine intranasal (IN) and intravenous (IV) infection models using *CTR4* promoter replacement strains (Fig. 3b), with the latter bypassing the pulmonary stage to model systemic infection. This approach was based on previous work showing that *CTR4* promoter-driven transcription is repressed during lung infection but induced during brain infection^35^. Five days after IV infection, fungal burdens in the brain and lungs were quantified. In the brain, five *CTR4* promoter replacement strains (*CDC39*, *PZF1*, *TOP3*, *EZT1* and *EZT2*) showed significant reductions in fungal burden compared with the wild-type, with *CDC39*, *EZT1* and *EZT2* displaying the most pronounced effects (Fig. 3c). In the lungs following IV infection, only the *FHL1 CTR4* promoter-replacement strain showed a significant reduction relative to the wild-type (Fig. 3c). In addition, IN infection revealed that three targets (*CEF1*, *CDC39*, and *EAT1*) showed reduced lung fungal burdens measured at day 5 compared to the wild-type strain (Fig. 3c), indicating that these essential transcriptional regulators influence the early stage of pulmonary infection.

We next reassessed virulence using *H3* promoter-driven constitutive overexpression strains (Fig. 3b), focusing on the essential regulators *CEF1*, *CDC39*, *EAT1*, *EZT1* and *EZT2*. The *CEF1oe* strain exhibited a reduced lung fungal burden compared with the wild-type strain, whereas the *EAT1oe* strain showed an increased lung fungal burden (Fig. 3d). In addition, *CDC39* overexpression significantly reduced brain fungal burden, and *EZT1* overexpression reduced brain fungal burden but increased lung colonisation (Fig. 3d). These findings indicate that a subset of essential transcriptional regulators exerts organ-specific control of cryptococcal virulence. Specifically, Eat1 acts as a positive regulator of pulmonary infection, whereas the balanced regulation of *CEF1* expression is required for effective lung infection. Moreover, the reduced brain fungal burden observed for *EZT1oe* and *CDC39oe* strains suggests that Ezt1 and Cdc39 contribute to the regulation of brain infection.

Collectively, our in vitro and in vivo phenotypic analyses suggest that the essential transcriptional regulators Cdc39 and Ezt1 are key factors of both brain and lung infection, whereas Cef1 and Eat1 primarily modulate pulmonary infection in *C. neoformans* (Fig. 3e).

#### Transcriptional networks governed by essential TFs Ezt1, Cbf1, and Ezt2 in *C. neoformans*

We next sought to delineate the transcriptional networks governed by evolutionarily divergent essential TFs Cbf1, Ezt1 and Ezt2 (Fig. 2e), two of which (Ezt1 and Ezt2) are linked to virulence. We performed RNA sequencing (RNA-seq) on the *EZT1oe*, *CBF1oe*, and *EZT2oe* strains and compared their transcriptomes with that of the wild-type strain under basal growth conditions. Notably, *EZT1* overexpression elicited extensive transcriptional remodelling, altering the expression of 1,264 genes—substantially more than observed upon overexpression of *EZT2* or *CBF1* (Fig. 4a, Supplementary Data 3 and Supplementary Fig. 9a). Principal component analysis (PCA) and the interquartile range (IQR) plot further showed that *EZT1* overexpression induced broader and higher-magnitude transcriptomic changes than overexpression of either *CBF1* or *EZT2* (Supplementary Fig. 9b and c), underscoring the broad regulatory scope of Ezt1 in *C. neoformans*.

**FIG 4.**
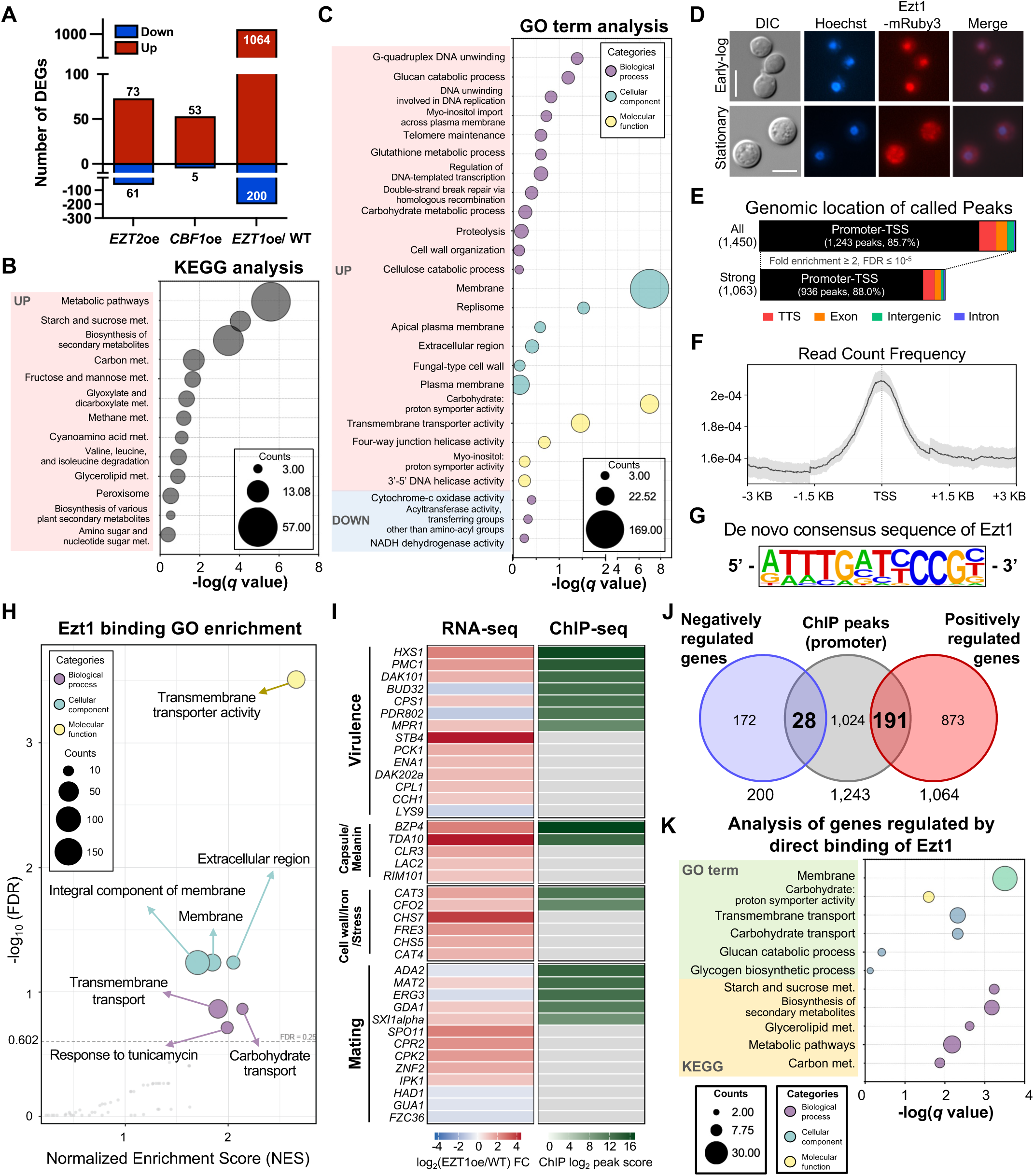
Transcriptional network regulated by *EZT1*, *EZT2*, and *CBF1* overexpression. (A) RNA sequencing (RNA-seq) analysis showing the number of differentially expressed genes (DEGs) in *EZT1oe*, *CBF1oe*, and *EZT2oe* strains compared to the wild-type strain (WT). (B) Kyoto Encyclopedia of Genes and Genomes (KEGG) pathway analysis showing the functional categories of genes regulated by *EZT1*. No KEGG pathways were significantly enriched among the downregulated genes. (C) Gene ontology (GO) analysis revealing that *EZT1* overexpression broadly upregulates genes involved in biological processes, cellular components, and molecular functions. (D) Cellular localisation of Ezt1 during exponential (6 h) and stationary (30 h) growth. Fluorescence microscopy of Ezt1-mRuby3 cells grown in YPD for 6 h and 30 h showed predominantly nuclear localisation of Ezt1 at 6 h and redistribution outside the nucleus at 30 h (scale bar: 5 μm). The complete growth-phase localisation analysis is shown in Supplementary Fig. 10C. (E) Genomic distribution of Ezt1-binding peaks annotated by HOMER. Of the 1,450 reproducible peaks identified, 1,243 (∼85.7%) were located in promoter–transcription start site (TSS) regions; 936 high-confidence promoter peaks (fold enrichment > 2, q < 10) were used for downstream motif analysis. (F) Metagene plot showing that Ezt1 occupancy is sharply centred on the TSS. The shaded grey region indicates the 95% confidence interval estimated by bootstrap resampling. This plot was generated using ChIPseeker in R. (G) De novo consensus binding motif identified at Ezt1-bound sites by HOMER. (H) Volcano plot of pre-ranked gene set enrichment analysis (FGSEA) of GO terms across genes ranked by Ezt1 promoter-proximal binding intensity. Each point represents a GO term; point size indicates gene set size, and fill colour indicates ontology category. The dashed horizontal line denotes FDR = 0.25. This plot was generated using fgsea in R. (I) Heatmap showing Ezt1 ChIP-seq peak scores (right, green) and RNA-seq log2 fold changes in *EZT1oe* versus WT (left, red–blue) for manuscript-referenced virulence-, capsule/melanin-, cell wall/iron/stress-, and mating-associated genes. Grey tiles indicate the absence of a promoter-proximal Ezt1 peak. Within each category, genes with Ezt1 binding are ordered by peak score (top), followed by genes without binding ordered by log2 fold change. This heatmap was generated using R. (J) Integration of RNA-seq and ChIP-seq data identified 219 genes candidate genes as direct Ezt1 regulatory targets, including 191 induced and 28 repressed genes. (K) GO analysis and KEGG pathway analysis of the candidate direct Ezt1 target genes.

KEGG pathway analysis of the differentially expressed genes (DEGs) indicated that Ezt1 regulates diverse metabolic programmes, including starch and sucrose metabolism, biosynthesis of secondary metabolites, carbon metabolism, amino acid degradation, glycerolipid metabolism, peroxisome and amino sugar/nucleotide sugar metabolism (Fig. 4b). No KEGG pathways were significantly enriched among the downregulated genes. In contrast, Cbf1 and Ezt2 affected a more restricted set of pathways: with Cbf1 affecting metabolic pathways, secondary metabolite biosynthesis, and amino acid degradation, while Ezt2 preferentially targeted pathways such as tyrosine metabolism (Supplementary Fig. 9d). Gene ontology (GO) term analysis using DAVID further showed that Ezt1 broadly upregulates genes involved in multiple biological processes, cellular components, and molecular functions (Fig. 4c). While Cbf1 and Ezt2 similarly regulated specific processes, their overall regulatory scope was markedly narrower than that of Ezt1 (Supplementary Fig. 9d).

Notably, Ezt1 regulated a wide range of virulence-associated genes in *C. neoformans*. Among the upregulated genes, fourteen (*STB4, HXS1*, *PMC1*, *ZNF2*, *PCK1*, *ENA1*, *DAK101*, *DAK202a*, *CPL1*, *CPS1*, *IPK1*, *RIM101*, *MPR1* and *CCH1*) have previously been implicated in virulence^13,48–61^, along with genes linked to capsule biosynthesis (*TDA10*, *BZP4*, *CLR3* and *RIM101*)^12,62–64^. *EZT1* overexpression also induced coordinated transcriptional changes in other virulence-related modules, including melanin biosynthesis and regulation (*LAC2* and *BZP4*)^65,66^, cell wall remodelling (*CHS7* and *CHS5*)^67^, iron acquisition (*FRE3*, *CFO2* and *STR1*)^68,69^, and oxidative stress defence (*CAT3* and *CAT4*)^70^. Furthermore, *EZT1* overexpression altered the expression of several sexual-development-associated genes (*SXI1*α, *CPR2*, *ZNF2*, *SPO11* and *CPK2*)^71–75^. Conversely, *EZT1* overexpression downregulated several pathogenicity-associated genes, including *PDR802*, *LYS9*, and *BUD32*^76–78^, as well as mitochondrially encoded electron transport genes such as cytochrome c oxidase III and NADH dehydrogenase subunits (Supplementary Data 3). Collectively, these findings highlight Ezt1 as a central, lineage-specific TF in *C. neoformans* that coordinates key virulence factors and related biological functions.

### Genome-wide identification of Ezt1 binding sites and regulated genes

To further investigate Ezt1 as a transcriptional regulator, we examined its subcellular localisation by tagging the C-terminus of Ezt1 with the mRuby3 fluorescent protein (Supplementary Fig. 10a). The Ezt1-mRuby3 strain in the H99L *MAT*α background phenocopied the wild-type strain under all conditions tested, indicating that C-terminal tagging with mRuby3 did not impair Ezt1 function (Supplementary Fig. 10b). Fluorescence microscopy showed that Ezt1-mRuby3 was predominantly nuclear during exponential growth, while time-course analysis revealed nuclear localisation during mid-lag and early-to-mid exponential growth, followed by relocalisation to the cytoplasm from late exponential phase through stationary phase (Fig. 4d and Supplementary Fig. 10c). These observations suggest a role for Ezt1 in regulating growth stage-dependent transcriptional programmes.

To validate Ezt1 as a nuclear DNA-binding protein, we generated an Ezt1-4×FLAG strain by tagging the C-terminus of Ezt1 with 4×FLAG epitopes (Supplementary Fig. 10a) in the H99L *MAT*α background. This strain phenocopied the wild-type strain, confirming that the Ezt1-4×FLAG protein is functional (Supplementary Fig. 10b and 10d). Using this strain, we performed chromatin immunoprecipitation sequencing (ChIP-seq) and identified 1,450 reproducible binding peaks (FDR < 0.05) by three-way intersection of three biological replicates. Of these, 1,243 (85.7%) were annotated to promoter-transcription start site (TSS) regions by HOMER, corresponding to 1,243 target genes, and 936 peaks with fold enrichment >2 and FDR < 10^-5^ were defined as high-confidence promoter binding sites (Fig. 4e and Supplementary Data 4). Metagene analysis showed that Ezt1 occupancy was sharply centred on the TSS (Fig. 4f). De novo motif analysis of these peaks revealed a conserved 12-bp motif (5’–ATTTGATYCCGY–3’, Y = C/T; *P* = 1 × 10^−1^^11^; Fig. 4g and Supplementary Data 4). Collectively, these results demonstrate widespread Ezt1 occupancy at promoter regions across the *C. neoformans* genome.

Pre-ranked gene set enrichment analysis (FGSEA), which assesses enrichment across a ranked gene list, revealed that strong Ezt1 occupancy was preferentially directed toward genes encoding transmembrane transporter activity, extracellular region components, and integral membrane components, together with transmembrane transport and carbohydrate transport processes (Fig. 4h). To examine how Ezt1 binding intersects with the virulence-, capsule/melanin-, cell wall/iron/stress-, and mating-associated genes identified in our RNA-seq analysis above, we constructed a gene-level integration map combining RNA-seq log_2_ Fold changes with Ezt1 ChIP-seq peak scores (Fig. 4i; Supplementary Data 3 and 4). Prominent Ezt1 occupancy was observed at promoters of key virulence factors (*HXS1*, *PMC1*, *DAK101*, *BUD32*, *CPS1*, *PDR802*, and *MPR1*)^13,48,52,53,56,61,77^, capsule/melanin regulators (*BZP4* and *TDA10*)^13,63^, and a subset of mating-associated genes (*ADA2*, *MAT2*, *ERG3*, *GDA1*, and *SXI1*α)^14,71,74,79,80^. Although *ADA2* did not meet the predefined RNA-seq threshold, it showed a modest but statistically significant decrease in expression (log_2_FC = -0.55). Other mating-associated genes with altered expression in *EZT1oe* (*SPO11*, *CPK2*, *ZNF2*, and *HAD1*)^14,72–74^ did not show detectable Ezt1 binding within their promoters, suggesting that their transcriptional changes may be mediated indirectly, downstream of Ezt1.

Building on this gene-by-gene view, we integrated the RNA-seq and ChIP-seq datasets to define the Ezt1 regulon at a genome-wide scale. We identified 219 candidate direct targets, comprising 191 induced and 28 repressed genes (Fig. 4j and Supplementary Data 4). Functional enrichment of these genes recapitulated the membrane-transport signature seen in the binding-intensity analysis (Fig. 4h): the induced set was enriched for membrane components and carbohydrate-related transport and metabolism (Fig. 4k).

To validate genes selected from the integrated ChIP-seq and RNA-seq analyses, we performed qRT-PCR on independently generated biological replicates of the *EZT1oe* strain. The 22-transcript panel comprised 15 genes that were both differentially expressed and bound by Ezt1 at their promoter regions (*HXS1*, *PMC1*, *DAK101*, *BUD32*, *CPS1*, *PDR802*, *MPR1*, *BZP4*, *TDA10*, *CAT3*, *CFO2*, *MAT2*, *ERG3*, *GDA1*, and *SXI1*α), four differentially expressed genes without detectable promoter binding (*CHS7*, *CHS5*, *CPK2*, and *ZNF2*), *EZT1* as an overexpression control, and *CPK1* and *DMC1*, which met neither criterion. qRT-PCR confirmed induction of *EZT1*, whereas *CPK1* and *DMC1* remained unchanged. Among the promoter-bound genes, *HXS1*, *DAK101*, *BZP4*, *TDA10*, *GDA1*, and *MPR1* increased significantly and *BUD32* and *ERG3* decreased (*P* < 0.05; Supplementary Fig. 11a), consistent with the RNA-seq data; *PMC1*, *CPS1*, *CFO2*, *PDR802*, and *SXI1*α changed concordantly but not significantly, and *CAT3* and *MAT2* showed no clear changes. All four genes lacking detectable promoter-binding were significantly induced, consistent with indirect regulation downstream of Ezt1. Across the 22 transcripts, qRT-PCR and RNA-seq log2 fold-change results correlated strongly (Pearson’s r = 0.907, 95% confidence interval = 0.785 to 0.961, *P<*0.0001; Supplementary Fig. 11b).

To complement this with a reduction-of-function approach, we examined the same panel following partial *EZT1* repression, comparing P*_CTR4_*:*EZT1* with the wild-type in the presence of CuSO_4_. *EZT1* expression was significantly reduced, confirming Cu-mediated repression (Supplementary Fig. 11a). *DAK101*, *TDA10*, *ERG3*, *SXI1*α, and *CAT3* decreased significantly, whereas *BZP4*, *CHS7*, *CHS5*, *CPS1*, and *MPR1* increased. Four genes upregulated in *EZT1oe* (*DAK101*, *TDA10*, *CAT3*, and *SXI1*α) changed in the opposite direction, but *CPS1*, *MPR1*, *BZP4*, *CHS7*, and *CHS5* increased under both conditions and *ERG3* decreased under both. Accordingly, transcriptional changes upon partial repression did not correlate with the *EZT1oe* profile (Pearson’s r = −0.0593, *P =* 0.7933; Supplementary Fig. 11c), indicating gene-selective, context-dependent responses rather than a global inversion.

Collectively, integrating ChIP-seq and RNA-seq with independent qRT-PCR validation supports the promoter-bound, Ezt1-responsive genes as candidate direct targets, whose transcriptional output is not governed by a simple reciprocal relationship with *EZT1* abundance. Together with the functional and phenotypic analyses, these findings support a model in which Ezt1 coordinates metabolic and virulence-associated programmes, contributing to both pathogenicity and cellular homeostasis in *C. neoformans*.

### Ezt1 is a negative regulator of sexual reproduction and morphogenic transitions

ChIP-seq and transcriptome analyses together suggest that Ezt1 is linked to the regulation of mating-associated genes, supporting a role for this factor in sexual differentiation in this pathogenic fungus. Consistent with this, overexpression of *EZT1* drastically reduced mating even in unilateral settings (Fig. 3a). To further support the transcriptional evidence linking Ezt1 to mating-associated pathways, we additionally constructed an *EZT1* overexpression strain in the *MAT***a** YL99 strain background^81^, driven by the *H3* promoter (Supplementary Fig. 12a). The *MAT***a** YL99 P*_H3_*:*EZT1* (*EZT1oe* **a**) strain phenocopied the *MAT*α H99L P*_H3_*:*EZT1* (*EZT1oe* α) strain, indicating that Ezt1 performs conserved functions in both mating types (Supplementary Fig. 12b). *EZT1* overexpression nearly abolished mating in both unilateral and bilateral mating set-ups (Fig. 5a), consistent with Ezt1 functioning as a negative regulator of mating. We quantified *EZT1* expression levels at multiple time points during mating by qRT-PCR. *EZT1* expression was significantly reduced up to 16 h post-mating but subsequently increased after 48 h (Fig. 5b). Notably, at 48 h and 72 h post-mating, Ezt1 became enriched at the cell periphery, consistent with a potential shift in its regulatory activity during mating progression (Fig. 5c).

**FIG 5.**
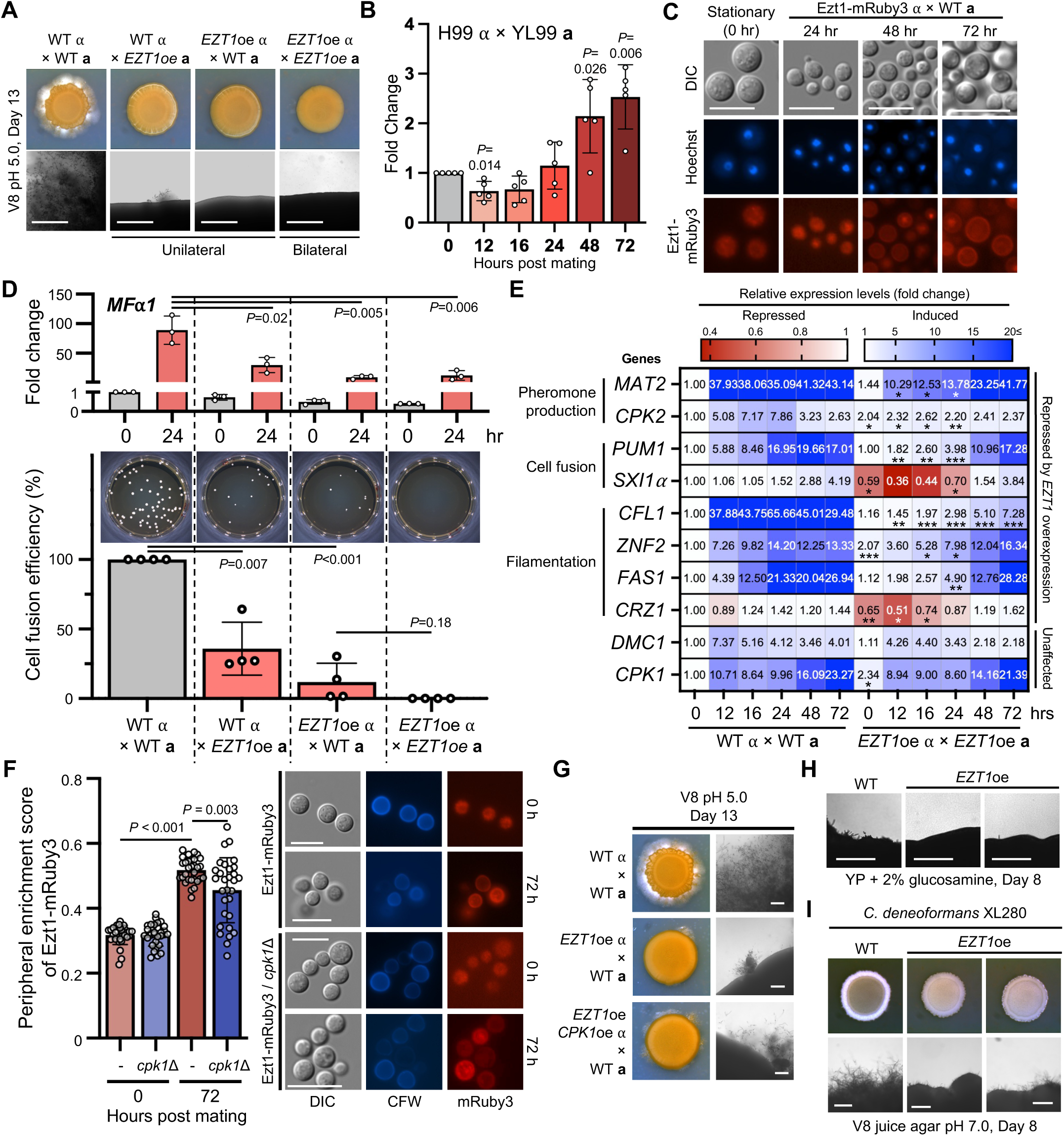
Ezt1 functions as a negative regulator of sexual reproduction and morphogenic transitions in *C. neoformans*. (A) Overexpression of *EZT1* in both *MAT*α H99L and *MAT***a** YL99 strains significantly inhibited mating in both unilateral and bilateral crosses (scale bar: 500 μm). (B) qRT-PCR analysis showing dynamic *EZT1* expression during mating. *EZT1* expression decreased up to 16 h post-mating but increased after 48 h. Data are represented as mean ± SD (standard deviation). Statistical significance was analysed at each time point relative to the starting point using a two-tailed one-sample *t*-test compared to 0 h time point (=1) with Prism 11.0 and *P* values were stated above (*n*=5). (C) Fluorescence microscopy of the Ezt1-mRuby3 strain showing Ezt1 translocation to the cell periphery after mating, with nuclei visualised by Hoechst staining (scale bar: 10 μm). (D) *EZT1* overexpression repressed the expression of the mating pheromone gene *MF*α*1*, reducing cell fusion efficiency. Data are represented as mean ± SD. Statistical significance for mating pheromone production was analysed using a two-tailed unpaired *t*-test with all of the samples compared to the WT cross (*n*=3). For cell fusion assay, the two unilateral crosses were compared to the *EZT1oe* × WT **a** cross using two-tailed unpaired *t-* test. All of the statistical tests were performed using Prism 11.0 (*n*=4). (E) *EZT1* overexpression delayed or reduced the expression of key mating-related genes, while genes such as *DMC1* and *CPK1* remained unaffected. In bilateral crosses, the temporal expression levels of each gene were statistically analysed by comparing them to the corresponding time points in wild-type mating using two-tailed unpaired *t*-test, and two-tailed one-sample *t*-test for 0 h time point with Prism 11.0 (*n*=5; * *P* < 0.05; ** *P* < 0.01; *** *P* < 0.001, *P* values for each gene and time points were listed at Supplementary Data 7). (F) Quantification of Ezt1-mRuby3 peripheral enrichment at 0 and 72 hr after mating induction. Peripheral enrichment was calculated as *P*/(*P+I*), where *P* and *I* represent the mean normalised mRuby3 fluorescence intensities in the peripheral and interior regions, respectively. Deletion of *CPK1* in the Ezt1-mRuby3 strain reduced Ezt1 peripheral enrichment during mating, indicating that Cpk1 modulates Ezt1 localisation. Each point represents the peripheral enrichment score of an individual cell (30 cells per condition). Statistical significance was determined using two-tailed Welch’s *t*-test with Prism 11.0 with *P* values stated together. (G) Co-overexpression of *CPK1* in the *EZT1oe* strain partially rescued the severe mating defect caused by *EZT1* overexpression, suggesting that Cpk1 regulates Ezt1 function. (H) *EZT1* overexpression reduced self-filamentation on glucosamine media (scale bar: 200 μm). (I) In *C. deneoformans* XL280, *EZT1* overexpression abolished unisexual reproduction (scale bar: 200 μm).

Initiation of sexual reproduction typically involves induction of mating pheromones followed by cell fusion. We found that *EZT1* overexpression significantly repressed *MF*α*1* expression, correlating with reduced cell-fusion efficiency (Fig. 5d). Moreover, in bilateral mating (*EZT1oe* α × *EZT1oe* **a**), *EZT1* overexpression significantly reduced expression of *SXI1*α and *CRZ1*, both of which are required for hyphal formation^71,82^. Induction of other key mating-associated genes involved in pheromone production, cell fusion and filamentation—including *CFL1*, *MAT2*, *ZNF2, CPK2*, *PUM1* and *FAS1*^74,83^—was delayed or reduced during the early phase (∼24 h) of sexual reproduction, whereas *DMC1* and *CPK1* expression was unchanged (Fig. 5e). Collectively, these data indicate that Ezt1 acts as a transcriptional negative regulator during the early stages of sexual differentiation in *C. neoformans*.

We next investigated the upstream regulator controlling Ezt1 during mating. Given the overlap in function, we hypothesised that Cpk1, a mitogen-activated protein kinase (MAPK) that governs pheromone production and cell fusion, regulates Ezt1. To test this, we deleted *CPK1* in the *EZT1-mRuby3* strain (Supplementary Fig. 13a) and monitored Ezt1 localisation during mating. Notably, quantitative analysis using calcofluor white (CFW) staining to define the cell boundary showed that the deletion of *CPK1* reduced the peripheral enrichment of Ezt1 at 72 h post-mating, indicating that Cpk1 promotes the late-stage peripheral localisation of Ezt1 (Fig. 5f and Supplementary Fig. 14). A previous study in the closely related species *Cryptococcus deneoformans* reported that *CPK1* disruption did not alter *EZT1* transcript levels during sexual reproduction^84^, suggesting that Cpk1-dependent regulation of Ezt1 need not involve changes in *EZT1* transcriptional abundance. Given that Ezt1 acts as a negative regulator during the early stage of mating, we therefore postulated that Cpk1 modulates Ezt1 post-translationally rather than by altering its expression. To further test this model, we generated a double overexpression strain by introducing a *CPK1* overexpression construct into the *EZT1oe* strain (Supplementary Fig. 13b). Under mating conditions, the severe mating defect caused by *EZT1* overexpression was partially rescued by *CPK1* overexpression, consistent with a functional interaction between Cpk1 and Ezt1 during mating (Fig. 5g).

In addition to its role in bisexual reproduction, we assessed the impact of Ezt1 on self-filamentation and unisexual differentiation. *EZT1* overexpression reduced self-filamentation on glucosamine medium, which induces this developmental programme^82^ (Fig. 5h). Moreover, in *C. deneoformans* strain XL280, which undergoes robust unisexual differentiation, *EZT1* overexpression abolished this process (Fig. 5i and Supplementary Fig. 15). Collectively, these findings indicate that Ezt1 functions as a central negative regulator of sexual reproduction and morphogenic transitions across the pathogenic *Cryptococcus* species complex.

### Functional complementation of *C. neoformans* Ezt1 by a divergent *Cryptococcus* orthologue

Because Ezt1 is among the most sequence-divergent essential cryptococcal TFs, we assessed its conservation across fungi using BLAST. Ezt1 sequence conservation was strongest within *Cryptococcus* species. Notably, Ezt1 is highly conserved among the pathogenic *Cryptococcus* species complex—including *C. neoformans*, *C. deneoformans*, *C. decagattii*, *C. deuterogattii*, *C. gattii*, *C. tetragattii* and *C. bacillisporus*—with e-values close to zero. Ezt1 orthologues were also detected in non-pathogenic *Cryptococcus* species, with significant sequence similarity (Fig. 6a). Notably, in *Cryptococcus depauperatus*, conservation was largely restricted to the central region containing the Zn2-Cys6 DNA-binding domain, resulting in a markedly shorter alignment than in other *Cryptococcus* species (Fig. 6a). In *Kwoniella* species, which are closely related to *Cryptococcus*, overall similarity to *C. neoformans* Ezt1 was substantially reduced, particularly outside the DNA-binding domain (Fig. 6a).

**FIG 6.**
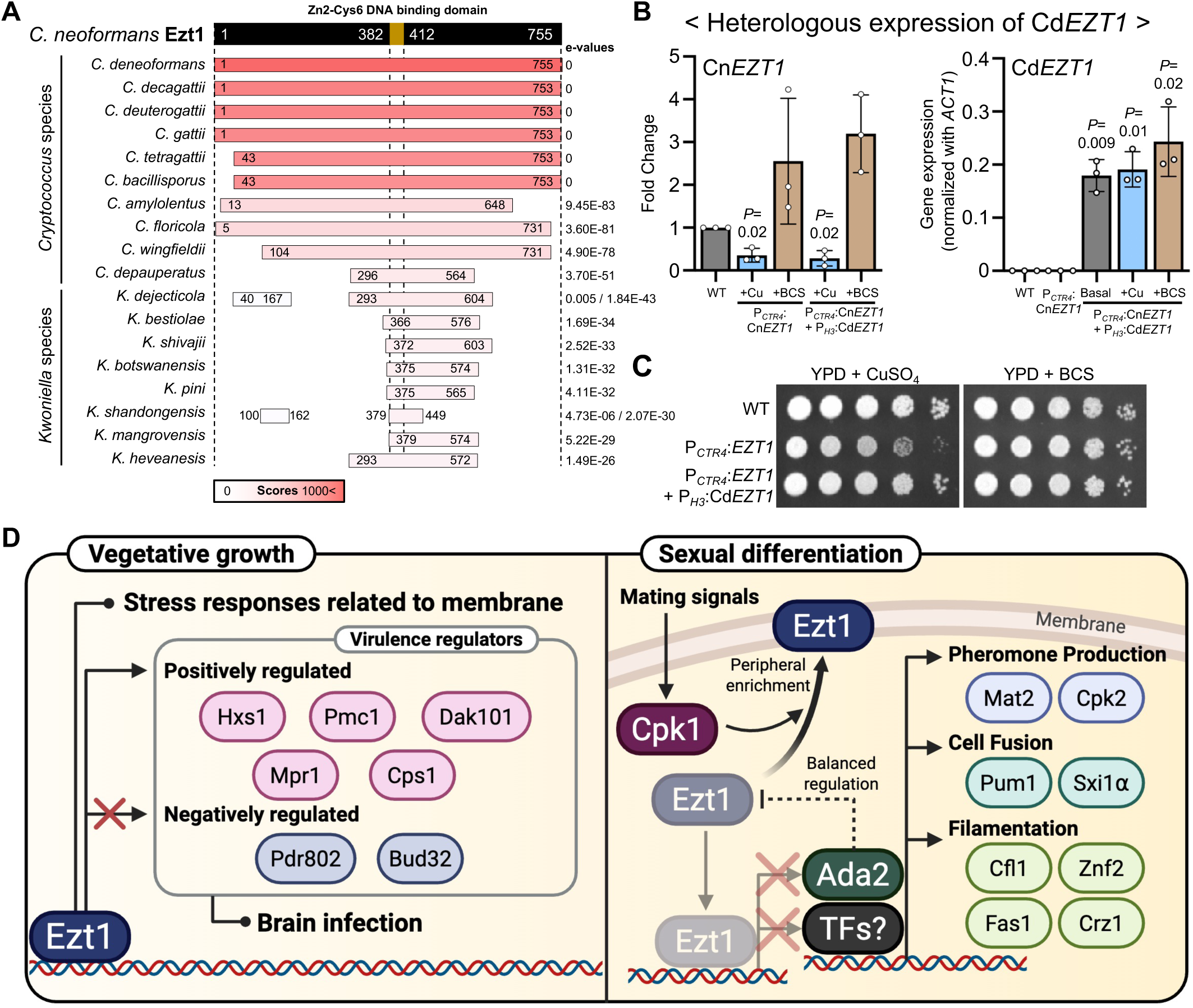
Functional complementation of *C. neoformans* Ezt1 by a divergent *Cryptococcus* orthologue. (A) BLASTP analysis showing sequence conservation of Ezt1 across *Cryptococcus* species. In *C. depauperatus*, only the Zn2-Cys6 DNA-binding domain in the middle region is conserved, with a significantly shorter alignment length compared to other species. (B) Functional complementation assay. Overexpression of *C. depauperatus EZT1* (*CdEZT1*) was tested in the *C. neoformans* conditional *EZT1* expression strain (P*_CTR4_*:*EZT1*) under copper-repressed conditions. qRT-PCR confirmed *CdEZT1* expression and conditional regulation of Cn*EZT1*. Data are represented as mean ± SD. Statistical significances of *CnEZT1* repression and *CdEZT1* overexpression were determined by comparison with wild-type expression levels using two-tailed one-sample *t*-test with Prism 11.0 against the respective reference values (*CnEZT1* = 1; *CdEZT1* = 0), and *P* values are stated above (*n*=3). (C) Growth assay showing that *CdEZT1* overexpression successfully restores wild-type growth of the P*_CTR4_*:*EZT1* strain under Cn*EZT1* repressed conditions, demonstrating that the *C. depauperatus* orthologue can compensate for reduced CnEzt1 expression under the conditions tested. (D) Proposed model of Ezt1 regulation in growth, stress responses, thermotolerance, virulence, and morphogenic development. Under normal growth conditions, Ezt1 controls membrane stress responses, positively regulating virulence factors (Hxs1, Pmc1, Cps1, Mpr1, and Dak101) while negatively regulating others (Pdr802 and Bud32), contributing to brain infection. During mating, Cpk1-dependent redistribution of Ezt1 towards the cell periphery is proposed to contribute to balanced regulation of pheromone production (Mat2 and Cpk2), cell fusion (Pum1 and Sxi1α), and filamentation (Cfl1, Znf2, Fas1, and Crz1). The interaction between Ezt1, Ada2, and other transcription factors remains to be fully elucidated. This image was created with BioRender.

We next tested whether the divergent Ezt1 orthologue from *C. depauperatus* (CdEzt1) could functionally complement *C. neoformans* Ezt1 (CnEzt1). To this end, we overexpressed *CdEZT1* in a *CTR4-*regulated conditional *EZT1* expression strain (P*_CTR4_*:*EZT1*) by integrating a P*_H3_*:*CdEZT1* allele into a previously characterised safe-haven locus^85^ (Supplementary Fig. 16). We confirmed the successful regulation of *CnEZT1* under *CTR4* conditions and constitutive *CdEZT1* overexpression (Fig. 6b). Strikingly, *CdEZT1* overexpression restored wild-type growth of the P*_CTR4_*:*EZT1* strain under *EZT1*-repressing conditions (Fig. 6c), demonstrating that CdEzt1 can compensate for reduced CnEzt1 expression in supporting growth under the conditions tested and suggesting conservation of a core growth-related Ezt1 function among *Cryptococcus* orthologues.

## Discussion

In this study, we identified and functionally characterised 13 essential transcriptional regulators (Esa1, Top3, Cef1, Cdc39, Rsc8, Snu66, Sgt1, Pzf1, Zap101, Eat1, Cbf1, Ezt1 and Ezt2) in *C. neoformans*, a leading cause of fungal meningitis worldwide. These factors were uncovered by prioritising 17 putative essential transcriptional regulators that had not been disrupted in previous genome-wide functional analyses of cryptococcal TFs^12^. Our findings are notable because, prior to this study, Hsf1 was the only essential TF to have been functionally characterised in *C. neoformans*^24^. We also established an experimental pipeline to define essentiality and interrogate in vitro and in vivo functions of essential cryptococcal proteins. To determine and validate essentiality, we used two complementary approaches: a conditional gene expression system and spore analysis of heterozygous mutants generated in engineered diploid strains. To investigate function, we constructed constitutive overexpression strains and performed a murine fungal burden assay using both conditional expression and overexpression strains. We propose that this integrated strategy provides a robust platform for the identification and functional characterisation of essential cryptococcal proteins.

Conditional repression from the *CTR4* promoter indicated that fourteen transcriptional regulators are required for growth. Subsequent spore analyses showed that thirteen of these are essential for viability in *C. neoformans*, whereas loss of Fhl1 resulted in impaired but viable growth, consistent with a quasi-essential role. By contrast, the remaining three TFs showed no growth defects upon *CTR4* promoter-mediated repression, and spore analyses confirmed that they are non-essential. Together, these results indicate that growth phenotypes obtained with the *CTR4* promoter-based conditional expression system are generally concordant with gene essentiality assessments in *C. neoformans*, although this conclusion is based on a limited number of genes. Despite this internal consistency, comparison with a previous transposon sequencing (Tn-seq) study revealed several discrepancies in essentiality assignments^86^. That study identified *ZAP101*, *EZT1* and *TOP3* as non-essential genes^86^. Moreover, *FHL1* was reported as essential in the Tn-seq study, yet our findings indicate that it is quasi-essential. These differences highlight how experimental conditions and genetic approaches can influence essentiality assignments. For example, genes required for spore germination may be scored as essential in our platforms even if they are dispensable at other life-cycle stages. Further work will be required to determine whether these genes are universally required throughout the cryptococcal life cycle, underscoring the value of multifaceted approaches for accurate essentiality assessment.

Our in vivo virulence assays, using murine infection models, with both *CTR4* promoter*-* based conditional expression and H3 promoter-driven constitutive overexpression strains, identified several essential transcriptional regulators associated with *C. neoformans* pathogenicity: *EZT1*, *EZT2* and *CDC39* for brain infection, and *EZT1*, *EAT1*, *CEF1* and *CDC39* for lung infection. The inhibitory effects of *EZT1* and *CDC39* overexpression on brain infection were consistently observed across both approaches. Moreover, conditional repression of *CDC39* reduced fungal burdens in the lungs, whereas *EZT1* overexpression increased lung colonisation, indicating that these essential regulators exert multifaceted roles during the infection cycle. Similarly, the positive role of *EAT1* in pulmonary colonisation was supported by both approaches. For *CEF1*, both repression and overexpression reduced lung fungal burden, indicating that tightly balanced *CEF1* expression is required for efficient pulmonary colonisation; this is consistent with our finding that both *CEF1* repression and overexpression markedly impaired in vitro growth. More generally, however, fungal burden in the P*_CTR4_* strains is not expected to track simply with in vitro growth, given the niche-dependent activity of the promoter noted above; reduced fungal burden in these strains should therefore not be interpreted solely as a consequence of impaired basal growth. Further studies will be required to define the molecular mechanisms by which these essential genes regulate virulence. Our results also showed that *FHL1* repression reduced lung colonisation, suggesting that quasi-essential genes may still exert substantial tissue-specific effects. Extending classical virulence assays to the newly identified quasi- or non-essential TFs Asr2, Fhl1, Fzc52, and Hlh7 should therefore provide additional insights into how these dispensable regulators contribute to cryptococcal pathogenicity. Collectively, delineating the roles of both essential and non-essential transcriptional regulators may inform the development of targeted antifungal strategies that exploit vulnerabilities specific to brain or lung infection.

A key finding of this study is the identification of Ezt1, an essential TF with strong sequence divergence outside *Cryptococcus* species. Although recent Tn-seq data suggested that Ezt1 may be non-essential in *C. neoformans*, the same dataset notably showed the absence of transposon insertions within the central DNA-binding domain and the nuclear localisation signal of Ezt1^86^. Together with our results, this pattern supports the conclusion that Ezt1 is an essential TF in this pathogenic fungus. Ezt1 is a fungal-specific Zn2-Cys6 TF conserved largely within the *Cryptococcus* species complex, yet absent or highly divergent in closely related taxa. Although evolutionarily divergent proteins are often assumed to be dispensable, our data demonstrate that Ezt1 is essential and regulates a broad range of pathobiological processes in *C. neoformans*. *EZT1* overexpression altered the expression of 1,264 genes, ∼60% of which were previously unannotated in *C. neoformans* and other fungi (Supplementary Data 3), underscoring the unique regulatory scope of Ezt1. Consistent with this, ChIP-seq identified Ezt1 binding at 1,243 genes under basal vegetative growth conditions, reinforcing its role as a broad-spectrum regulator of the *C. neoformans* transcriptome, while qRT-PCR independently supported overall reproducibility of the *EZT1oe*-associated transcriptional changes. Notably*, EZT1* overexpression markedly reduced brain infection by *C. neoformans*. This effect may be partially attributable to the significant downregulation of *PDR802*, a TF required for crossing the BBB^77^, and *BUD32*, a KEOPS complex component implicated in tRNA modification and pathogenicity^76^ (summarised in Fig. 6d). Partial *EZT1* repression, by contrast, elicited both reciprocal and non-reciprocal responses. Because *CTR4-*mediated repression requires altered copper conditions and prolonged depletion of an essential regulator may trigger secondary effects, copper-independent, acute and titratable systems will be needed to resolve the dosage-dependent Ezt1 regulon.

Our integrated ChIP-seq and RNA-seq analyses identified candidate direct Ezt1 targets under basal vegetative growth conditions, including genes involved in diverse metabolic pathways and in membrane structure and function. Consistent with this, in vitro phenotypic profiling showed that altered Ezt1 abundance affects tolerance to temperature extremes, osmotic stress and membrane stress—phenotypes closely linked to membrane fluidity, integrity and composition^87–89^. Notably, *EZT1* overexpression increased sensitivity at 25 and 39, underscoring a key role for Ezt1 in thermal adaptation. Interestingly, recent ChIP-seq data from Gao et al. showed that Hsf1 directly binds to the *EZT1* promoter under heat shock conditions (37 and 40)^90^. Consistent with this, our own data from a separate study also support an Hsf1-*EZT1* regulatory link (unpublished data), suggesting that Hsf1 may act upstream of *EZT1* to promote membrane stability and cell survival at elevated temperatures. Additionally, the antifungal agents to which *EZT1* overexpression increased susceptibility (amphotericin B, fluconazole and fludioxonil) are known to target processes associated with membrane stability^91–93^. Together, these observations support the conclusion that balanced Ezt1 activity contributes to membrane homeostasis and stress adaptation in *C. neoformans*.

Our findings indicate that Ezt1 functions as a central morphogenetic regulator in *C. neoformans*. *EZT1* overexpression strongly inhibited mating, cell fusion, self-filamentation and unisexual reproduction, and qRT-PCR confirmed altered expression of mating genes at early stages of sexual differentiation. Integrated transcriptomic and ChIP-seq analyses showed that Ezt1 binds a subset of mating-associated genes under basal vegetative conditions. Because these binding profiles were obtained under non-mating conditions, and because *EZT1* overexpression also alters stress responses, the transcriptional changes seen during sexual development likely reflect both direct regulation and indirect effects on adaptation to mating-induced conditions.

Deletion of *CPK1* impaired Ezt1 enrichment at the cell periphery during mating, and *CPK1* overexpression partially suppressed the *EZT1* overexpression mating defect. These observations indicate a functional relationship between Cpk1 and Ezt1 but do not establish direct regulation of Ezt1 by Cpk1. Because the two proteins do not appear to regulate each other’s transcript levels during mating^84^, the mechanism may involve post-transcriptional or post-translational control of Ezt1, or indirect effects of Cpk1 on mating-associated processes. We therefore regard the Cpk1-Ezt1 relationship as a working model. This model is compatible with the proposal of Jang et al. that an uncharacterised TF downstream of Cpk1 regulates Mat2^74^. Ezt1 bound the promoter of *ADA2*, whose deletion impairs mating^12^, and modestly repressed its expression under basal conditions; conversely, Ada2-dependent chromatin regulation at the *EZT1* promoter represses *EZT1*^79^, suggesting a reciprocal interplay that may help ensure appropriate mating progression. Defining these connections and their influence on Ezt1 localisation and activity will require interaction mapping and post-translational analysis. Collectively, these findings raise the possibility that Ezt1 integrates metabolic, virulence-associated and morphogenetic programmes, which may partly underlie its essential role in this pathogen (Fig. 6d).

How Ezt1, a lineage-divergent protein, evolved into an essential TF remains unclear. Its Zn2-Cys6 DNA-binding domain is highly conserved across *Cryptococcus* species and, unusually, resides in the central rather than the N-terminal region typical of fungal TFs^94^; the flanking N- and C-terminal regions are highly divergent and may contribute to *Cryptococcus*-specific transcriptional networks. Consistent with the primacy of the central domain, the *C. depauperatus* orthologue, which retains conservation only in this region, restored growth funder partial *CnEZT1* repression. ChIP-seq revealed widespread Ezt1 occupancy across the genome, supporting a model in which Ezt1 influences multiple transcriptional programmes underlying essential metabolic pathways and core biological processes. These data do not, however, establish how Ezt1 recognises target promoters, how binding is translated into activation or repression, or how the conserved central domain and the divergent terminal regions each contribute. Domain dissection, DNA-binding-defective mutants and interaction mapping will therefore be important for defining its structure-function relationships and evaluating its therapeutic potential.

Despite the evolutionary divergence that makes TFs attractive antifungal drug targets, they have been overlooked because their activity is difficult to inhibit. Nonetheless, TF-targeting strategies have been pursued in cancer and other diseases^15–17^, and in *C. albicans* the azole-responsive sterol regulator Upc2 has been used to screen for small molecules that block its expression or DNA binding^95^. We therefore propose Ezt1, Eat1 and Ezt2 as prioritised candidates for anti-cryptococcal drug discovery, given their structural divergence, essentiality and contributions to virulence, although their tractability as targets remains to be established.

The conserved essential transcriptional regulators identified here—Esa1, Top3, Cef1, Cdc39, Rsc8, and Snu66—govern fundamental machinery required for growth and survival and may likewise warrant investigation. Host toxicity is an obvious concern, but subtle structural divergences or fungus-specific interactions with binding partners could permit selective inhibition; alternatively, targeting their virulence-critical downstream genes may offer a complementary strategy. Establishing tractability will require regulatable conditional expression systems, such as a tetracycline-repressible knockdown system^96^, which would provide precise temporal control during infection and avoid the expression fluctuations of the *CTR4* promoter in the host. Our work thus provides a foundation for therapeutic approaches against this globally important pathogen by defining essential fungal transcriptional regulators and their associated vulnerabilities.

Several limitations should be noted. Functional characterisation relied on constitutive overexpression from the *H3* promoter and conditional repression from the *CTR4* promoter, neither of which recapitulates native expression dynamics; our conclusions therefore concern altered regulator dosage rather than complete loss of function, and residual Ezt1 activity under partial repression may contribute to the gene-selective responses observed. In vivo analyses were limited to fungal burden at a fixed endpoint, without survival, dissemination kinetics or direct measurement of gene expression in cells recovered from infected tissues. Consequently, this study does not establish Ezt1, or any other essential regulator identified here, as a validated antifungal target: we have not shown that acute depletion or pharmacological inhibition produces the growth arrest or clearance required for therapeutic development. Finally, in-depth analysis was confined to Ezt1, leaving both its biochemical mechanism and the functions of the remaining twelve essential regulators to be defined.

## Methods

### Ethics statement

Animal experiments were performed in accordance with the ethical guidelines of the Institutional Animal Care and Use Committee of Jeonbuk National University (NON2023-132). The committee approved all vertebrate studies, and all experiments adhered to the experimental ethics guidelines.

### Construction of *C. neoformans CTR4* promoter replacement strains

*CTR4* promoter replacement cassettes were generated by double-joint PCR (DJ-PCR), as previously described^97^, using nourseothricin-resistant marker (nourseothricin N-acetyltransferase; *NAT*). The primer pair B354/B355 was used to amplify the *NAT*-*CTR4* promoter with pNAT-*CTR4* as a template. Approximately 800 bp of the 3′ region of the promoters and 800 bp of the 5′ region of the exons of the target genes were amplified using the primer pairs *CTR4*_L1/*CTR4_*L2 and *CTR4_*R1/*CTR4*_R2, respectively, with H99L genomic DNA (gDNA) as the template (Supplementary Data 5). After amplification of the 3′ promoter regions, 5′ exon regions, and *NAT*-*CTR4* in the first round of PCR, primer pairs *CTR4*_L1/SM2 and *CTR4*_R2/SM1 were used to amplify 5′ and 3′ regions of the *NAT*-*CTR4*-split cassettes in the second round of PCR. All primers and generated *CTR4* promoter replacement strains were listed in Supplementary Data 5 and 6. The targeted insertion cassettes were then introduced into the *C. neoformans* H99L using biolistic transformation, as described previously^98^. The H99L strain was cultured in liquid yeast extract-peptone-dextrose (YPD) medium (1% yeast extract, 2% peptone, and 2% dextrose) for 16 h at 30 in a shaking incubator, spun down, resuspended in 5 ml distilled water, plated onto YPD agar plates containing 1 M sorbitol, and further incubated at 30 for 3 h. Purified targeted insertion cassettes were prepared with 600 μg of 0.6 μm gold microcarrier beads (Bio-Rad, CA, USA) using calcium chloride and spermidine and delivered into H99L using a particle delivery system (PDS-100, Bio-Rad). Plates were incubated at 30 for 4 h to restore the disrupted cell membrane, scraped, and spread on YPD agar plates containing 100 μg/ml nourseothricin. Stable transformants were selected and further screened for correct insertion by diagnostic PCR. Finally, Southern blot analysis was performed to validate the correct genotypes of the promoter-replacement strains using gene-specific probes.

### Construction of *C. neoformans* heterozygous mutants

Single copies of the target genes were deleted in the diploid *C. neoformans* AI187 and CnLC6683 through homologous recombination using gene disruption cassettes containing signature-tagged *NAT.* The gene disruption cassettes were generated by DJ-PCR with the primer pairs listed in the Supplementary Data 5. PCR amplified the 5′- and 3′-flanking regions of the target genes with the primer pairs L1/L2 and R1/R2, using gDNA as a template, while the M13Fe/M13Re pair was used to amplify signature-tagged *NAT*. Split-gene deletion cassettes were then generated using the primer pairs L1/SM2 and R2/SM1 and introduced to the parental diploids via biolistic transformation, as described above. Stable transformants were selected, screened with diagnostic PCR, and confirmed through Southern blot analysis with gene-specific probes. All of the generated heterozygous mutants were listed in Supplementary Data 6.

### Whole-genome sequencing of the constructed heterozygous mutants

Genomic DNA for whole-genome sequencing (WGS) was extracted using either a MasterPure Yeast DNA Purification Kit (LGC Biosearch Technologies, MPY80200)^32^, or a CTAB-based extraction method^99^, as previously described. Illumina WGS was performed on a NovaSeq 6000 platform either at the Duke Sequencing and Genomic Technologies Core Facility (https://genome.duke.edu) or by Macrogen Inc. (Seoul, Republic of Korea). Paired-end whole-genome sequencing reads were aligned to the *Cryptococcus neoformans* reference genome assembly (GCA_000149245.3) using BWA-MEM v0.7.19 with the default parameters. The resulting alignments were coordinate-sorted and indexed using SAMtools v1.6. Genome-wide read-coverage tracks were generated from the sorted BAM files using the ‘bamCoverag’ function in deepTools v3.5.1 with a bin size of 100 bp. The sorted BAM and BigWig files were visualized using the Integrative Genomics Viewer v2.16.1. Chromosome-wide read-coverage profiles were visually inspected to assess coverage uniformity and to identify gross chromosome-level coverage abnormalities indicative of aneuploidy or large-scale copy-number alterations.

### Construction of *C. neoformans* constitutive overexpression strains

The essential transcriptional regulators were subjected to overexpression by replacing their native promoters with the histone *H3* promoter using targeted insertion, similar to the *CTR4* promoter replacement method. The primer pairs *CTR4*_L1/*CTR4*_L2 and *H3*_R1/*CTR4*_R2 were used to amplify the 3′ region of the promoter and the 5′ region of the exons, respectively, with H99L gDNA as a template (primers listed in Supplementary Data 5). The primer pair B4017/B4018 was used to amplify the *NAT*-*H3* or *NEO* (G418/neomycin resistance marker)*-H3* promoter, using pNAT-H3 or pNEO-H3 as the template. The second round of PCR was conducted using primer pairs *CTR4*_L1/SM2 and *CTR4*_R2/SM1 to construct the *NAT*-*H3*-split cassettes. These generated cassettes were introduced into the H99L strain via biolistic transformation. Stable transformants were selected on YPD plates containing nourseothricin, screened with diagnostic PCR, and confirmed through Southern blot analysis with gene-specific probes. Two independent overexpression strains had their genotypes and overexpression levels confirmed by Southern blot analysis and quantitative reverse transcription PCR (qRT-PCR), respectively. All the overexpression strains used in this study are listed in Supplementary Data 6.

### Expression analysis by quantitative RT-PCR

To monitor the gene expression levels in conditional expression and constitutive overexpression strains, qRT-PCR was performed. For *CTR4* promoter replacement strains, cells were cultured in liquid YPD medium at 30 for 16 h in a shaking incubator. The cultures were then diluted to an OD_600_ of 0.2 with fresh YPD, YPD containing 40 μM CuSO_4_, and YPD containing 200 μM of bathocuproinedisulfonic acid (BCS; a copper chelator). The mixtures were cultured at 30 in a shaking incubator for 8 h, collected by centrifugation, and subjected to overnight lyophilisation. For constitutive overexpression strains, cells were cultured in liquid YPD medium at 30 for 16 h in a shaking incubator, diluted to an OD_600_ of 0.2 with fresh YPD, and grown further at 30 in a shaking incubator until reaching an OD_600_ of ∼0.8. Cell cultures were collected by centrifugation and lyophilised overnight. For mating-associated gene expression, *MAT*α and *MAT***a** strains were cultured in liquid YPD medium at 30 ℃, washed three times with phosphate-buffered saline (PBS), and combined at equal concentrations (10^8^ cells/mL). A 500 μl aliquot of the cell mixture was plated onto V8 agar medium (5% V8 juice, 4% agar, pH 5.0) and incubated at room temperature in the dark for designated time points. Cells were then scraped and lyophilised overnight. Total RNA was extracted using the Easy-Blue RNA extraction kit (iNtRON Biotechnology, Korea), and cDNA synthesis was performed using Maxima H Minus RTase (Thermo Scientific, USA) with 5 μg of RNA. qRT-PCR was conducted with gene-specific primers (Supplementary Data 5) on the CFX96 Touch Real-Time PCR Detection System (Bio-Rad), as previously described^74^. Relative gene expression fold change was analysed using the 2^-ΔΔCT^ method^100^, and data were illustrated using Prism 11.0.

For independent validation of *EZT1* and its candidate target levels in the constitutive *EZT1* overexpression strain and the copper-repressed P*_CTR4_*:*EZT1* strain, cells were cultured and harvested under the corresponding conditions described above. Frozen cell pellets were disrupted by bead-beating four times for 45 sec with 1 min chilling on ice between cycles. Total RNA was extracted using the RNeasy Kit (Qiagen) according to the manufacturer’s instructions and treated with RNase-free DNase to remove contaminating genomic DNA. cDNA was synthesised using the iScript cDNA Synthesis Kit (Bio-Rad). qRT-PCR was performed using Fast SYBR Green Master Mix (Applied Biosystems) on a CFX384 Real-Time PCR System (Bio-Rad). Relative transcript abundance was calculated using the 2^-ΔΔCT^ method^100^ with *ACT1* as the reference gene.

### Sporulation, spore separation, and spore analysis of heterozygous mutants

For spore purification and random spore analysis of heterozygous mutants constructed in the AI187 background, we followed a previously reported protocol with slight modifications^34,101^. Heterozygous mutants were induced to sporulate on V8 juice agar pH 5.0. Cells were cultured at 30 for 16 h in a shaking incubator, collected, washed twice with PBS, adjusted to a density of 10^7^ cells/ml, and spotted (3 μl) on V8 juice agar plates. The plates were incubated in the dark for 4 to 6 weeks to induce unisexual reproduction and sporulation. Once spores were visible under the microscope, the mixed cell mass, including filaments, basidia, spores, and yeast cells, was suspended in 75% Percoll^®^ (Sigma Aldrich, MO, USA) made isotonic with PBS. The suspensions were then centrifuged at 3,000 rpm for 20 min at 4 to generate a gradient. Spores formed a band near the bottom of the gradient, while all other cell types remained at the top. The spore band was collected by puncturing the tube wall with a 20 ml tuberculin syringe, followed by centrifugation and two washes with PBS. Spores were diluted 1/100 and spread onto yeast nitrogen base (YNB) medium with amino acids and ammonium sulphate, supplemented with 1 mg/ml uracil, 2 mg/ml adenine, and 1 mg/ml 5-fluoroorotic acid (5-FOA; for excluding diploid *ura5*/*URA5* yeast cells). Plates were incubated at 30 for 7 days, and the resulting colonies were subjected to mating type locus PCR using mating type-specific primer pairs (Supplementary Data 5). Spores with confirmed mating types were used for further analysis. To test genetic markers segregation during meiosis (*ade2*/*ADE2*, *ura5*/*URA5*, *MAT***a**/*MAT*α, and *NAT*), spores were cultured at 30 in liquid YPD for 16 h and spotted onto the following plates: YPD, YPD + 100 μg/ml nourseothricin, YNB, YNB + 2 mg/ml adenine, and YNB + 1 mg/ml uracil plates. Knockout spores were identified by their growth on YPD + nourseothricin and confirmed by internal PCR using gene-specific primers.

For direct spore dissection analysis, heterozygous mutants constructed in the AI187 or CnLC6683 strain backgrounds were sporulated on MS medium (pH 5.8). Spores were mechanically detached through microscopic manipulation using a fibre optic needle spore dissection system, as previously described^101,102^. After germination, progeny demonstrating growth on YPD + nourseothricin were analysed by internal PCR to confirm gene knockouts as described above.

### Growth and chemical susceptibility test

*C. neoformans* cells were grown at 30 in liquid YPD for 16 h, serially diluted (1 to 10^4^), and then spotted on YPD plates containing various chemicals to induce stress, as previously reported^12^. The types of stresses tested and chemicals used to induce them were as follows: antifungal drug susceptibility [Fludioxonil (FDX), flucytosine (5-FC), fluconazole (FCZ), and amphotericin B (AMB)], toxic heavy metal stress [cadmium sulphate (CdSO_4_, abbreviated to CDS)], cell wall stress [Congo Red (CR) and sodium dodecyl sulphate (SDS)], oxidative stress [hydrogen peroxide (HPX), tert-butyl hydroperoxide (TBH), menadione (MD), and diamide (DIA)], genotoxic stress [hydroxyurea (HU) and methyl methanesulphonate (MMS)], and ER stress [tunicamycin (TM) and dithiothreitol (DTT)]. Osmotic pressures were induced with 1.5 M NaCl, 1.5 M KCl, and 2 M sorbitol in YPD medium, as well as 1 M NaCl, 1 M KCl, and 2 M sorbitol in YPD without dextrose (YP) medium. The spotted plates were incubated at 30 ℃ and photographed daily for 1 to 5 days using a Bio-Rad Gel-doc imaging system. Growth phenotypes were scored using the same semi-quantitative scoring method established in our previous studies^12–14^. Phenotypes were scored relative to the corresponding wild-type strain on an ordinal scale ranging from -3 to +3, based on the difference in the highest 10-fold dilution exhibiting visible growth under each condition. Scores of 0, ±1, ±2, and ±3 represented no difference, approximately one, two, or three (or more) dilution differences, respectively. Positive scores indicated enhanced growth or resistance, whereas negative scores indicated reduced growth or increased sensitivity.

### Capsule and melanin induction assays

For capsule and melanin induction assays, *C. neoformans* strains were grown overnight in liquid YPD medium at 30 and washed twice with PBS. For melanin production, 3 μl of each cell suspension was spotted onto Niger seed (35 g/L) agar supplemented with the indicated concentrations of glucose and incubated at 37 for the indicated amounts of time. Melanisation was documented using a BX41 microscope (Olympus, Japan) equipped with a SPOT digital camera (Diagnostic Instruments, USA). For capsule induction, 3 μl of each cell suspension was spotted onto Dulbecco’s modified Eagle (DME) agar and incubated at 37 for 2 days. Cells were collected and mixed with Remel™ BactiDrop India Ink to counterstain the polysaccharide capsule. Images were acquired by a differential interference contrast microscope ECLIPSE Ni (Nikon, Japan) equipped with DS-QI2 camera, and capsule thickness was quantified by measuring both cell body and total cell diameters using Nikon NIS software. Capsule thickness was calculated as (total diameter – cell body diameter)/2.

Capsule and melanin phenotypes were scored using the same semi-quantitative scoring system established in our previous studies^12–14^. For capsule, categorical scores were assigned only when the difference in capsule thickness relative to the wild-type strain was statistically significant. Melanin production was assessed qualitatively by visual comparison of pigmentation with the wild-type strain. Phenotypes were evaluated relative to the wild-type strain on an ordinal scale ranging from -3 to +3, with positive values indicating increased capsule production or melanisation and negative values indicating reduced capsule production or melanisation. A score of 0 indicated no detectable difference from the wild-type. The resulting scores were used for heatmap generation.

### Murine infectivity assays

Animal experiments in this study were conducted at the Core Facility Center for Zoonosis Research (Jeonbuk National University, South Korea). Inbred 6-week-old female BALB/cAnNCrlOri mice, confirmed as SPF/VAF, were purchased from ORIENT BIO INC. (Korea) and acclimated to the breeding environment for one week before the experiment. Infections with *CTR4* promoter-driven conditional expression strains were performed with reference to a previous study^35^. Each strain was inoculated into fresh liquid YPD medium and cultured overnight at 30 with shaking. An equal number of 5×10^4^ cells in 40 μl of PBS were administered via lateral tail vein injection (IV) or intranasal inhalation (IN). For IN, mice were anaesthetised with isoflurane vapour (Hana Pharm. Co., Ltd., South Korea) at a flow rate of 80 cc/min. Mice were humanely sacrificed 5 days post-infection, and fungal burden was measured in both the brain and lungs (IV model) or in the lungs (IN model).

For the murine infectivity assay using *H3* promoter-driven constitutive overexpression strains, an equal number of 5×10^4^ cells in 40 μl of PBS was administered via IN. Mice were sacrificed 14 days post-infection, and fungal burden was measured in both the brain and lungs. To quantify fungal burden, tissue samples were homogenised in 1 ml of PBS and plated on YPD medium supplemented with 100 μg/ml chloramphenicol. The plates were incubated at 35 ℃ for two days, after which the fungal burden was quantified and expressed as the logarithmic CFU per gram of tissue. Statistical analysis was performed using Welch’s *t*-test on log-transformed fungal burden (CFU/gram) to compare each tested strain with the wild-type control using GraphPad Prism 11.0.

### Transcriptome analysis of the *EZT1, EZT2,* and *CBF1* overexpression strains

The wild-type (H99L), P*_H3_*:*CBF1*, P*_H3_*:*EZT1*, and P*_H3_*:*EZT2* strains were grown in YPD liquid medium at 30 overnight. The cultures were then transferred to fresh YPD liquid medium and incubated until the OD_600_ reached 0.8. Three biologically independent cultures were prepared for each strain, and total RNA was purified. A cDNA library was constructed using 1 µg of total RNA for each sample with the Illumina TruSeq RNA library kit v2 (Illumina) and sequenced using the Illumina platform. The reads were processed to remove adaptor and low-quality sequences using Cutadapt v2.4 with Python 3.7.4 with adapter sequence^103^. The reference genome sequence of *C. neoformans* H99 and annotation data were obtained from the NCBI FTP server. The reads were aligned to the genome sequence using Hisat2 v2.2.1 with the Hisat and Bowtie2 algorithm, as previously reported^104^. Hisat2 was performed with the “-p 30” and “—dta -1” options, while other parameters were set to default. Aligned reads were converted and sorted using Samtools v0.1.19 with default parameters except for the “-Sb -@ 8” option for converting and the “-@ 20 –m 2000000000” option for sorting^105^. Transcript quantification was performed using the featureCounts command within the Subread package v2.0.8, installed via Bioconda^106^. The parameters used for featureCounts were “-T 4-M—largestOverlap -g gene_id”. Differential gene expression analysis was performed using DESeq2 v1.24. The volcano plot was illustrated using R v4.1.0, with the cutoff set to more than two-fold changes with *P* < 0.05.

### Chromosomal tagging of Ezt1 with mRuby3 and 4×FLAG

To express Ezt1 fusion proteins tagged with either 4×FLAG or mRuby3 tag, two separate constructs were generated using the following procedure. For the Ezt1-4×FLAG construct, the 3′ exon region was amplified using primer pair B19021/B10922, and the terminator region was amplified with B10923/B10924. For the Ezt1-mRuby3 construct, the 3′ exon region was amplified using B19021/B20540, and the terminator region was amplified with B20541/B19024. The *NAT* marker containing *4×FLAG*-*HOG1t*-*NAT* allele was PCR-amplified with primer pair B5536/B1966, whereas the *mRuby3-HOG1t-NEO* allele was amplified using B16758/M13Re.

The amplified 3′ exon, terminator, epitope, and selection marker regions were fused via DJ-PCR; for the 3′ exon, primer pair B10921/B1445 was used for 4×FLAG, and B10921/B1997 for mRuby3. For the terminator, B10924/B1444 were used for 4×FLAG, and B10924/B1886 for mRuby3. Finally, the resulting targeted insertion cassettes were introduced to the H99L strain via biolistic transformation. Correct genotypes were confirmed by performing 3′ junction PCR using primers JOHE12579/B18848, followed by additional validation through Southern blot analysis with an *EZT1*-specific probe amplified using primers B19068/B19069.

### Chromatin immunoprecipitation-sequencing (ChIP-seq) of the Ezt1-4×FLAG strain

The *C. neoformans* Ezt1-4×FLAG strain was cultured in liquid YPD at 30 overnight. Chromatin extraction was performed as previously reported^62^. The culture was transferred to fresh YPD liquid medium and incubated until the OD_600_ reached 0.8. In vivo crosslinking was achieved by adding 37% formaldehyde to a final concentration of 1%, and the mixture was incubated at 30 for 5 min with agitation. Crosslinking was quenched by adding 2.5 M glycine (to a final concentration of 300 mM), and the mixture was incubated at 30 for an additional 5 min with agitation. Cells were collected, washed twice with ice-cold PBS, and once with ice-cold FA lysis buffer (50 mM HEPES pH 7.5, 140 mM NaCl, 1 mM EDTA, 1% Triton X-100, 0.1% sodium deoxycholate, 0.1% SDS, and protease inhibitors), and resuspended in 0.8 ml of ice-cold FA lysis buffer. Cells were homogenised using a bead beater (FastPrep-24 5G, MP Biomedicals) for 5 intervals of 2 min each, with 1 min of cooling on ice between intervals. The lysate was centrifuged for 2 min at 4,000 rpm, and the supernatant was subjected to sonication until the DNA fragment size reached 150-500 bp. A 20 μl aliquot of each chromatin sample was reserved prior to immunoprecipitation as a matched input control, rather than a separate mock immunoprecipitation control, and DNA was purified using a DNA purification kit (Cosmogenetech, South Korea).

ChIP was performed as previously described^62^. Protein A Dynabeads^®^ (60 μl per sample) were washed twice with ice-cold PBS containing 5 mg/ml bovine serum albumin (BSA). The beads were resuspended in 60 μl of PBS with 5 mg/ml of BSA, and anti-FLAG antibody (Sigma-Aldrich, USA) was added. Each sample was incubated at 4 overnight with gentle rotation. The antibody-conjugated beads were washed twice with ice-cold PBS containing 5 mg/ml BSA and then aliquoted into four tubes. The extracted chromatin was added to the washed beads, and each sample was incubated at 4 for 3 h with gentle rotation. After incubation, the beads were washed three times with FA lysis buffer, three times with FA lysis buffer containing 0.5 M NaCl, and once with TE buffer (10 mM Tris-HCl and 10 mM EDTA, pH 8.0), using a magnetic separator (General Electric, USA). DNA was eluted from the beads with 35 μl of elution buffer (50 mM Tris-HCl, pH 8.0, 10 mM EDTA, and 1% SDS) at 65 for 1 min. The eluted samples were treated with 15 μl of 10% SDS, 1 μl of RNase H, and 3 μl of RNase cocktail (A&T, Ambion, USA) at 65 overnight. The samples were then treated with 7 μl of proteinase K at 42 for 90 min to reverse the crosslinking. Reverse-crosslinked DNA was purified using a PCR purification kit (Cosmogenetech, South Korea). Following evaluation of DNA quality by absorbance measurement using a NanoDrop spectrophotometer (Thermo Scientific, USA), sequencing libraries were constructed using the TruSeq ChIP-seq library preparation kit (Illumina, USA). Libraries were sequenced on an Illumina NovaSeq 6000.

### ChIP-seq data processing and binding-site analysis

Raw reads were preprocessed to remove adaptor sequences and filter out low-quality reads using Cutadapt (v2.4; Python 3.7.4). Cleaned reads were aligned to the *C. neoformans* H99 reference genome using HISAT2 (v2.2.1) with default parameters. Aligned reads were converted to BAM format and sorted using SAMtools (v0.1.19). Peak calling from each ChIP-seq replicate was performed using MACS2 (v2.2.7.1), with the corresponding matched input chromatin sample used as the control for enrichment analysis, consistent with established fungal ChIP-seq approaches^62,107,108^. Significant peaks were identified based on a threshold of ≥ 2-fold enrichment and an adjusted *P* value < 0.05. Reproducible Ezt1-binding sites detected in all three biological replicates were extracted using BEDTools (v2.31.1), and the resulting consensus peaks were annotated to the *C. neoformans* H99 genome using HOMER (v5.1). High-confidence promoter-binding sites, used for motif analysis and downstream enrichment tests, were defined as peaks located in promoter–TSS regions with fold enrichment > 2 and q < 10^-5^. De novo and known motif analyses on the high-confidence peak set were performed using HOMER. Binding profiles were visualised using the Integrative Genomics Viewer (v2.16.1), and metagene analysis around TSSs (±3 kb) was performed using the ChIPseeker package in R, weighted by peak intensity to emphasise strongly bound sites, with 95% confidence intervals estimated by bootstrap resampling.

To assess the functional categories preferentially associated with strong Ezt1 binding without applying arbitrary peak-score cut-offs, pre-ranked gene set enrichment analysis was performed using the fgsea package in R. HOMER-annotated peaks within ±2 kb of the nearest TSS were collapsed to the gene level by retaining the maximum peak score per gene, log_2_-transformed, and ranked in decreasing order. GO annotations for C. neoformans H99 were obtained from FungiDB (release 68); some GO terms (e.g., GO:0016021, GO:0055114) have since been deprecated and merged into broader categories in recent GO releases, and their historical names are retained here for clarity. fgsea was run separately for Biological Process, Molecular Function, and Cellular Component ontologies with gene sets filtered to contain 10– 500 genes. Multiple-testing correction was performed using the Benjamini–Hochberg procedure applied separately within each ontology, and GO terms with FDR < 0.25 were considered significantly enriched.

### Mating, filamentation, and cell fusion assays

To evaluate bisexual mating efficiency in both unilateral and bilateral settings, wild-type and genetically engineered strains constructed in *MAT*α H99L and *MAT***a** YL99 strain backgrounds were cultured in YPD medium at 30 for 16 h, washed twice with PBS, and combined at equal cell densities (10 cells/ml). The mixtures were plated onto V8 juice agar (pH 5.0) and incubated in the dark at 25 for 7 to 14 days. For glucosamine-induced filamentation, H99L and *EZT1oe MAT*α strains were cultured in YPD medium at 30 for 16 h, washed twice with PBS, and adjusted to a density of 10 cells/ml. A 3 μl aliquot of resuspended cells was spotted onto YP + 2% glucosamine plates and incubated at 30 for 2 days before transferring to 25 to observe self-filamentation. Filamentous growth was monitored and documented via photography. To induce unisexual reproduction in *C. deneoformans* XL280 and its derived *EZT1oe* strains, cultures were prepared as described above, and 3 μl of cell suspension was plated onto V8 juice agar (pH 7.0). Plates were incubated in the dark at 25 for 5 to 10 days, and filamentous growth was monitored and photographed using a differential interference contrast microscope ECLIPSE Ni (Nikon, Japan) equipped with DS-QI2 camera and a BX51 microscope (Olympus, Japan) with a SPOT digital camera (Diagnostic Instruments, Inc. USA).

For assessing cell fusion efficiency, we followed an established protocol^109^. Equal volumes of *MAT*α and *MAT***a** control or tested strains, carrying *NAT* or *NEO* markers, respectively, were cultured in liquid YPD medium at 30 for 16 h and combined at a cell density of 10^7^ cells/ml. A 5 μl aliquot of the mixed cell suspension was plated onto V8 juice agar (pH 5.0) and incubated in the dark at room temperature for 24 h. Cells were collected by scraping and resuspended in 1 ml of distilled water, and a 20 μl aliquot was plated onto a YPD medium supplemented with nourseothricin (100 μg/ml) and G418 (50 μg/ml). Colony counts were recorded after 4 days of incubation in the dark at room temperature.

### Fluorescent microscopy for visualisation of Ezt1-mRuby3

For growth-stage-dependent localisation, overnight culture of Ezt1-mRuby3 strain was diluted to an OD_600_ of 0.2 in fresh YPD medium and incubated at 30 with shaking. Aliquots were collected at 2, 6, 12, 24, and 30 h after inoculation and processed for fluorescence microscopy as described below. For analysis of localisation during mating, overnight cultures of the Ezt1-mRuby3 *MAT* strains and the wild-type *MAT***a** YL99 strain were separately diluted to fresh YPD medium at an OD_600_ of 0.2 and cultured at 30 for 48 h. The cells were then washed twice with PBS, adjusted to equal concentrations of 10^7^ cells/ml, mixed at a 1:1 ratio, and spotted (3 μl) onto V8 juice agar (pH 5.0) to initiate mating. The mating cultures were incubated in the dark at 25, and samples were harvested at the indicated time points and processed for fluorescence microscopy as described below. Cells collected from the growth-stage and mating-stage experiments were fixed with 4% paraformaldehyde containing 3.4% sucrose for 15 min at room temperature. After fixation, the cells were pelleted, washed with 0.1 M potassium phosphate buffer (pH 7.5) containing 1.2 M sorbitol, and stored in potassium phosphate buffer until further analysis. For nuclear staining, the cells were incubated with an equal volume of 20 μg/ml Hoechst 33342 (Thermo Fisher Scientific) in the dark for 30 min. Samples were examined and imaged using a differential interference contrast and fluorescence microscopy with a Nikon Eclipse microscope (Nikon, Tokyo, Japan) equipped with a DS-Qi2 digital camera.

### Quantification of Ezt1-mRuby3 peripheral enrichment

Ezt1-mRuby3 *MAT* × WT *MAT***a** and Ezt1-mRuby3 *cpk1*Δ *MAT* × WT *MAT***a** crosses were analyzed at 0 and 72 hr after mating induction on V8 juice agar plates. After fixation, cells were treated with equal volume of 25 μg/ml CFW (calcofluor white) and incubated for 30 min in the dark. Cells were washed thrice with PBS, and mounted on glass slides and imaged. For each cell, a line was drawn through the cell center and across two opposing CFW-stained cell walls using Fiji/imageJ. Cell-specific background intensity was estimated from the cell-free regions flanking the cell and subtracted separately from the mRuby3 and CFW fluorescence profiles. The background-corrected CFW profile was smoothed using a three-point moving average for boundary detection. The highest CFW peak on each side of the cell center was defined as the corresponding cell-wall boundary. The region between the two boundaries was normalized from 0 to 100% of the cell width, and the fluorescence profiles were linearly interpolated to 101 equally spaced position at 1% intervals. Negative background-corrected mRuby3 values were set to zero, and each interpolated mRuby3 signal profile was normalised to its maximum value. Peripheral intensity (*P*) was defined as the mean normalised mRuby3 signal intensity within the 0-15% and 85-100% regions of the cell width. Interior intensity (*I*) was defined as the mean normalised mRuby3 signal intensity across the 41 positions within the 30-70% region of the cell width. Peripheral enrichment was calculated as *P*/(*P+I*). A score of 0.5 indicates equal mean fluorescence intensity in the peripheral and interior regions, whereas values above and below 0.5 indicate peripheral- and interior-biased distributions, respectively.

### Heterologous expression of *CdEZT1* in *C. neoformans*

To construct a *C. neoformans* P*_CTR4_:EZT1* strain heterologously overexpressing *Cryptococcus depauperatus EZT1* (*CdEZT1*), the overexpression cassette was integrated into a safe haven region as previously described^85^. The intergenic region between CNAG_00777 and CNAG_00778 was divided into two fragments and amplified by PCR using primer pairs B21128/B21129 and B21130/B21131, respectively. Overlap PCR was then performed to introduce AscI, BaeI, and PacI restriction enzyme sites into the middle of the intergenic region. The JEC21 *H3* promoter (P*_H3_*), amplified with primer pair B21132/B21133, was fused to this intergenic region using overlap PCR. The fused product was assembled into the pHYG vector^110^ using Gibson assembly (New England Biolabs, USA), creating the pHYG-Safe Haven (SH)-P*_H3_*plasmid, and its sequence was verified by sequencing. Using this vector, the *EZT1* orthologous gene (L203_03018) from *C. depauperatus* CBS7841 was amplified by PCR using primer pair B21896/B21897 and inserted into the vector via Gibson assembly (New England Biolabs, USA). The resulting construct, pHYG-SH-P*_H3_*-Cd*EZT1* plasmid, was linearised using PacI restriction digestion, purified, and introduced into P*_CTR4_*:*EZT1* (YSB9280). Stable transformants were selected on YPD plates containing hygromycin and confirmed with diagnostic PCR. Overexpression of *CdEZT1* was confirmed by qRT-PCR.

## Supporting information

Description of Addtional Supplementary Informations

## Acknowledgements

This work was supported by National Research Foundation of Korea (NRF) funded by the Korean government (MSIT) (2018R1C1B6009031 and 2022R1C1C2003274 to K.-T.L., and RS-2025-00555365, RS-2025-02215093, and RS-2025-18362970 to Y.-S.B.); by a grant from the Korea Health Industry Development Institute (KHIDI), funded by the Ministry of Health & Welfare, Republic of Korea (RS-2022-KH131220 and RS-2026-25518347 to Y.-S.B.); and Korea Institute for Advancement of Technology (KIAT) funded by the Ministry of Trade, Industry and Energy in 2026 (RS-2024-00418203 to Y.-S.B.). The work was supported in part by Brain Korea 21(BK21) FOUR program, by AmtixBio, Co., Ltd. (to Y.-S.B.), and by NIH/NIAID R01 grants AI039115-28 and AI050113-20 (to J.H.). L.E.C. is supported by a Canadian Institutes of Health Research Foundation grant (FDN-154288) and is a Canada Research Chair (Tier 1) in Microbial Genomics & Infectious Disease (CRC-2023-00184). L.E.C. and J.H. are co-directors and fellows of CIFAR program Fungal Kingdom: Threats & Opportunities program. We also extend our gratitude to Dr. Aaron P. Mitchell and Dr. Eunsoo Do from the University of Georgia for providing invaluable guidance with the protocols for our ChIP-seq experiments, and to Dr. Blake Billmyre from the University of Georgia for insightful discussions on the interpretation of his Tn-seq data.

## Data availability

Source data and analysis scripts are available on figshare (https://doi.org/10.6084/m9.figshare.32099587.v2). RNA-seq and ChIP-seq data have been deposited in the Gene Expression Omnibus (GEO) under accession numbers GSE328125 and GSE328126, respectively. WGS data have been deposited in NCBI BioProject under accession number PRJNA1459063 (https://dataview.ncbi.nlm.nih.gov/object/PRJNA1459063). All deposited datasets will be publicly available upon publication.

## Author contributions

Y.-S.B. conceived the project. S.-H.L., S.S., Y.-B.J., S.-R.Y., J.-T.C., Y.C., and E.-S.K. performed experiments and analysed the data. J.-S.L., J.H., K.-T.L., and Y.-S.B. supervised the experimental analysis. S.-H.L., S.S., Y.-B.J., S.-R.Y., J.-T.C., Y.C., E.-S.K., J.-S.L., L.E.C., J.H., K.-T.L., and Y.-S.B. wrote the manuscript. All authors reviewed and approved this manuscript.

