## Supplementary Information for "Systematic Profiling of Essential Fungal Transcriptional Regulators Uncovers Ezt1 as a Central Pathobiological and Morphogenic Regulator in *Cryptococcus neoformans*"

This PDF file includes:

**Supplementary Figures 1 – 16**

**Supplementary figure 1 (Lee et al.)**

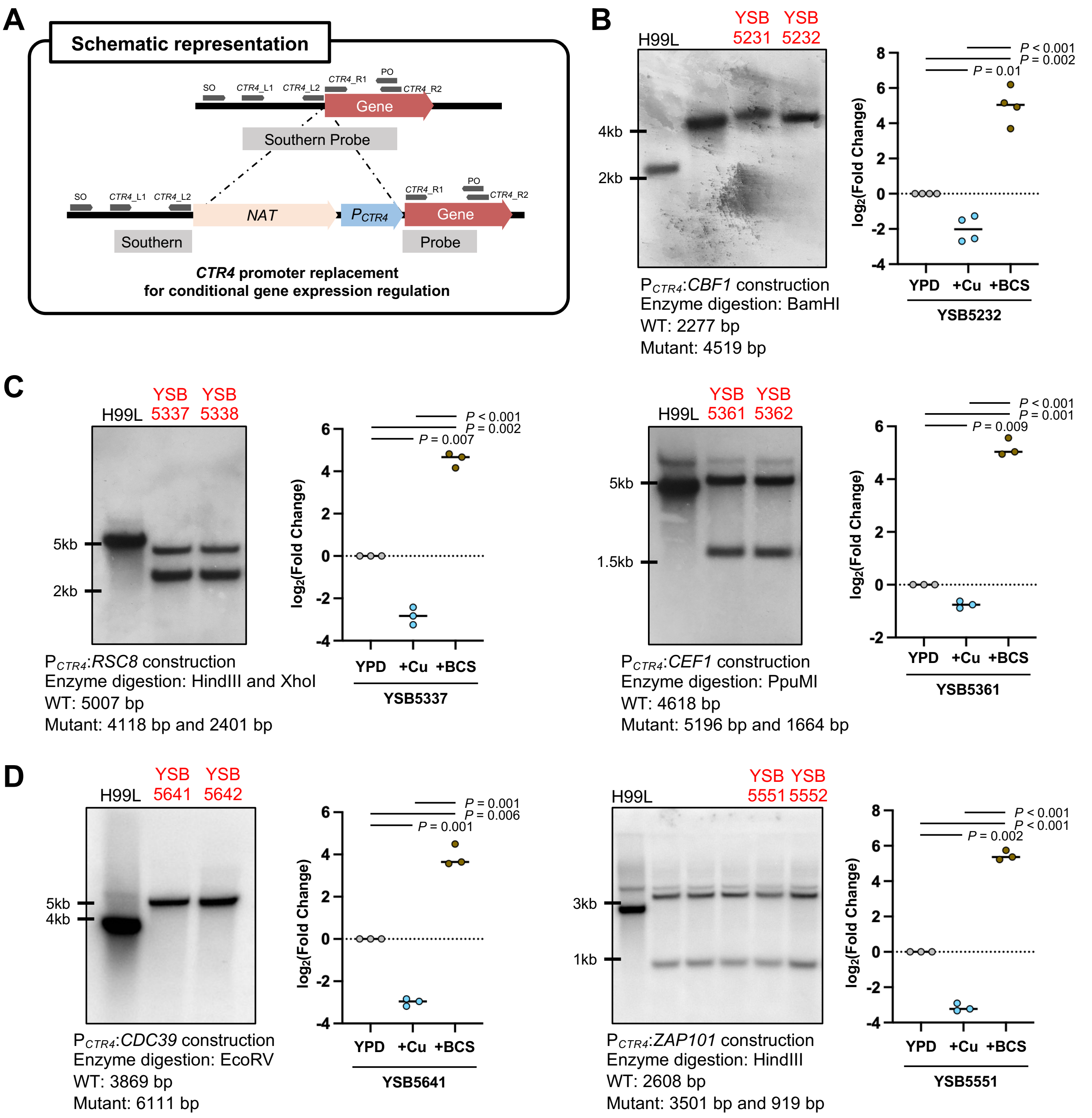


***Continued***

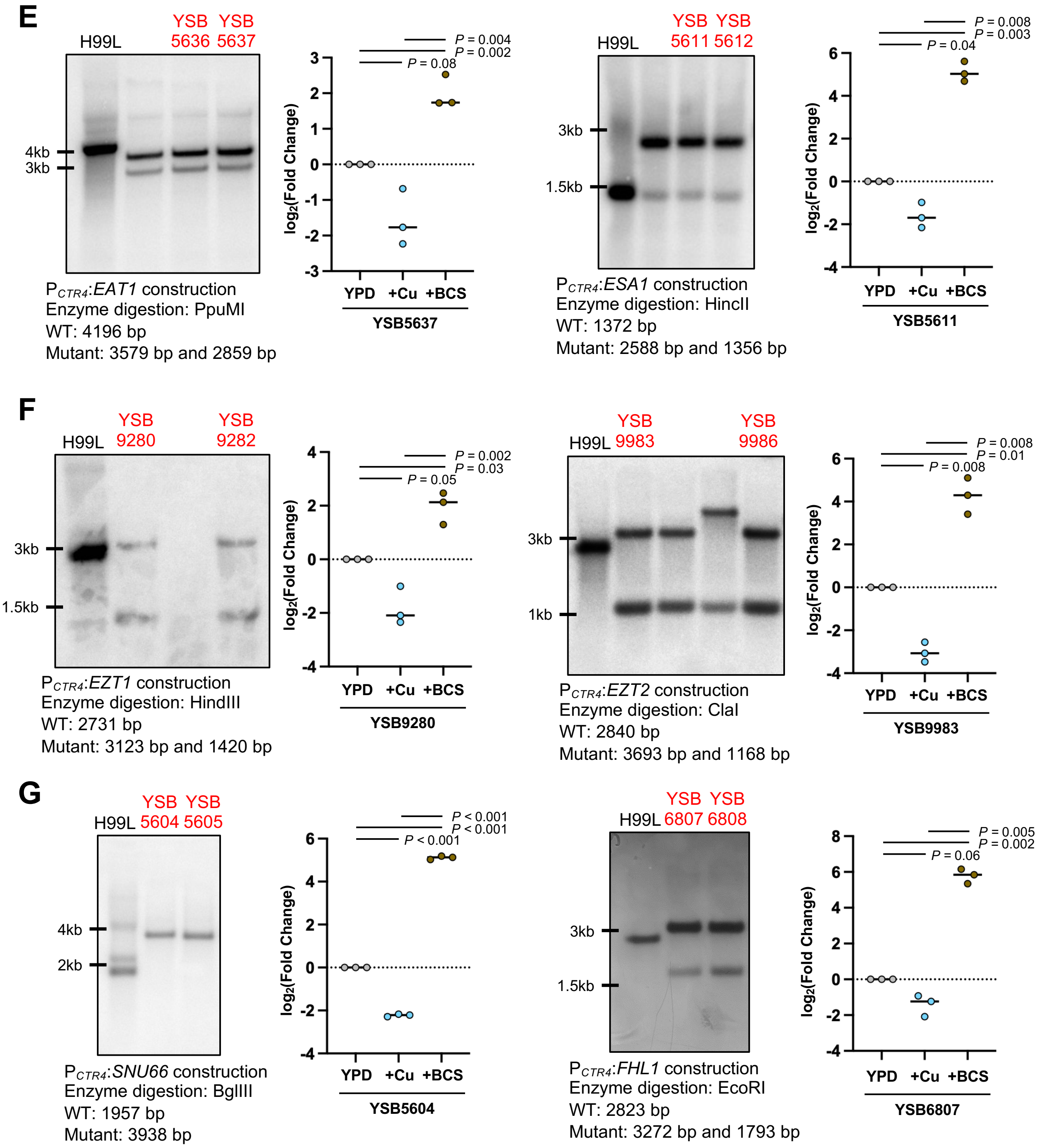
E
H99L
YSB 5636
YSB 5637
4kb
3kb
PCTR4:EAT1 construction
Enzyme digestion: PpuMI
WT: 4196 bp
Mutant: 3579 bp and 2859 bp
P = 0.004
P = 0.002
P = 0.08
log2(Fold Change)
YPD
+Cu
+BCS
YSB5637
H99L
YSB 5611
YSB 5612
3kb
1.5kb
PCTR4:ESA1 construction
Enzyme digestion: HincII
WT: 1372 bp
Mutant: 2588 bp and 1356 bp
P = 0.008
P = 0.003
P = 0.04
YSB5611
F
H99L
YSB 9280
YSB 9282
3kb
1.5kb
PCTR4:EZT1 construction
Enzyme digestion: HindIII
WT: 2731 bp
Mutant: 3123 bp and 1420 bp
P = 0.002
P = 0.03
P = 0.05
YSB9280
H99L
YSB 9983
YSB 9986
3kb
1kb
PCTR4:EZT2 construction
Enzyme digestion: ClaI
WT: 2840 bp
Mutant: 3693 bp and 1168 bp
P = 0.008
P = 0.01
P = 0.008
YSB9983
G
H99L
YSB 5604
YSB 5605
4kb
2kb
PCTR4:SNU66 construction
Enzyme digestion: BglIII
WT: 1957 bp
Mutant: 3938 bp
P < 0.001
P < 0.001
P < 0.001
YSB5604
H99L
YSB 6807
YSB 6808
3kb
1.5kb
PCTR4:FHL1 construction
Enzyme digestion: EcoRI
WT: 2823 bp
Mutant: 3272 bp and 1793 bp
P = 0.005
P = 0.002
P = 0.06
YSB6807

***Continued***

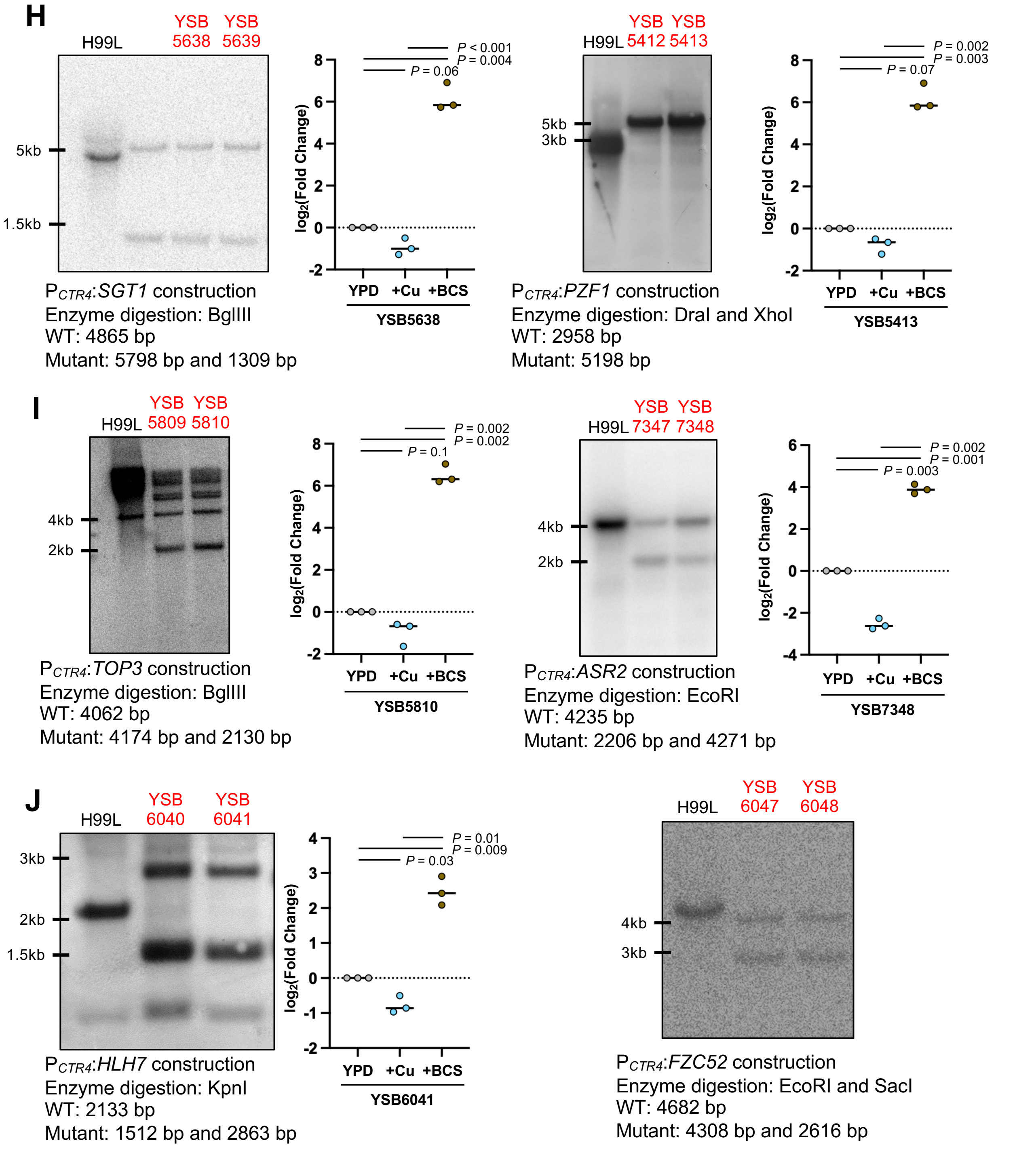


**Supplementary figure 1. Construction of the *CTR4* promoter replacement strains.** (A) Schematic representation for the construction of the conditionally regulatable strains. (B - J) Correct genotypes of the strains were confirmed through Southern blot analysis with the designated restriction enzyme digestion of genomic DNA and gene-specific probes. The regulation of the targeted genes was confirmed with quantitative reverse transcription-PCR (qRT-PCR). The strains were grown overnight at 30°C in 2 ml of liquid YPD medium ($OD_{600}$=0.2), subcultured into 10 ml of fresh YPD medium, YPD + 40 μM $CuSO_4$, and 200 μM BCS (bathocuproinedisulphonic acid) respectively. Cell cultures were further incubated at 30°C in a shaking incubator for eight hours, harvested, frozen in liquid nitrogen, and lyophilized. Total RNAs were isolated using easy-BLUE™ total RNA extraction kit (iNtRON biotechnology, Republic of Korea), and cDNAs were synthesized using Maxima H minus reverse transcriptase followed by the manufacturer's protocol (Thermo Fisher Scientific, USA). qRT-PCRs were performed using gene-specific primer pairs, and the expression levels of the genes were normalized with *ACT1* expression. Individual data are shown, with horizontal lines indicating the mean. Statistical significance of difference of repression and induction was determined by one-sample *t*-test with normalized values of YPD conditions (log2 fold change=0) using Prism 11.0 (*P* values are indicated above or below the bar graph), and differences between CuSO4 and BCS conditions were assessed using two-tailed paired *t*-test (*P* values indicated above).

**Supplementary figure 2 (Lee et al.)**

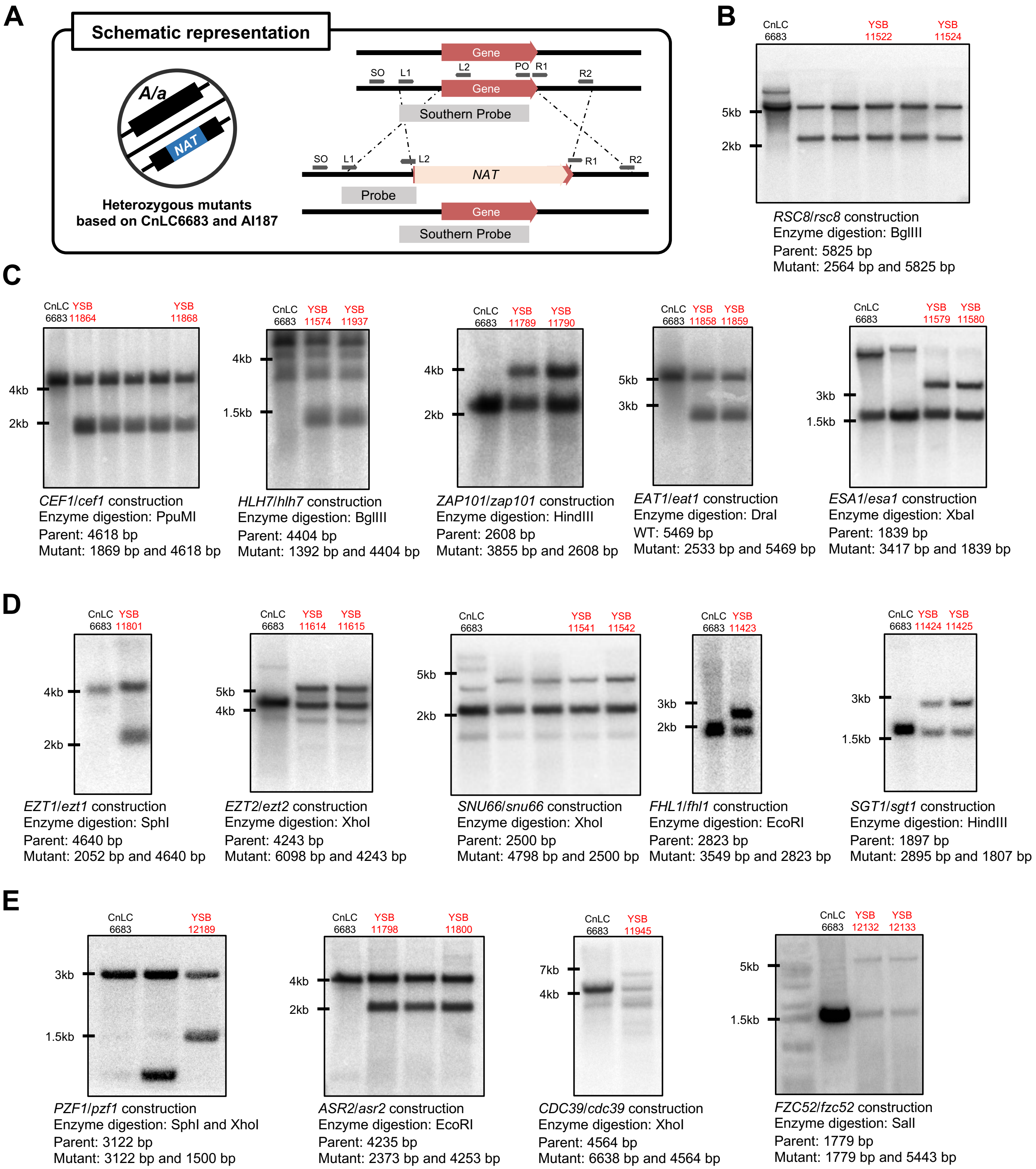


**Supplementary figure 2. Construction of the heterozygous mutants based on diploid CnLC6683.** (A) Schematic representation for the construction of the heterozygous mutant strains in the genetically engineered diploid CnLC6683 strain. (B-E) Correct genotypes of the heterozygous mutant strains were confirmed through Southern blot analysis with the designated restriction enzyme digestion of genomic DNA and gene-specific probes.

### Supplementary figure 3 (Lee et al.)

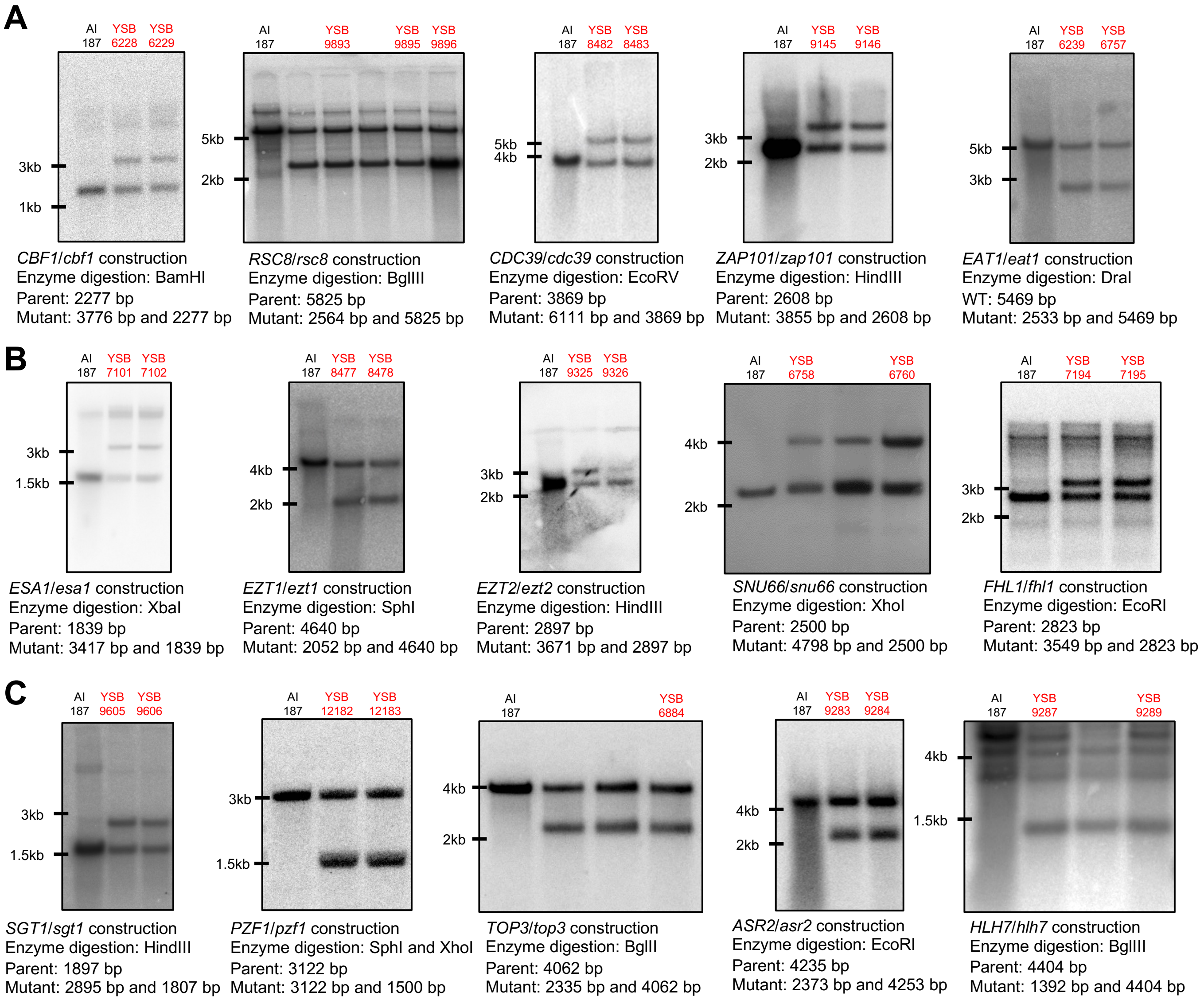


**Supplementary figure 3. Construction of the heterozygous mutants based on diploid AI187.** Schematic representation for the construction of the heterozygous mutant strains in the diploid AI187 strain is shown in the supplementary figure 2A. (A-C) Correct genotypes of the heterozygous mutant strains were confirmed through Southern blot analysis with the designated restriction enzyme digestion of genomic DNA and gene-specific probes.

Supplementary figure 4 (Lee et al.)

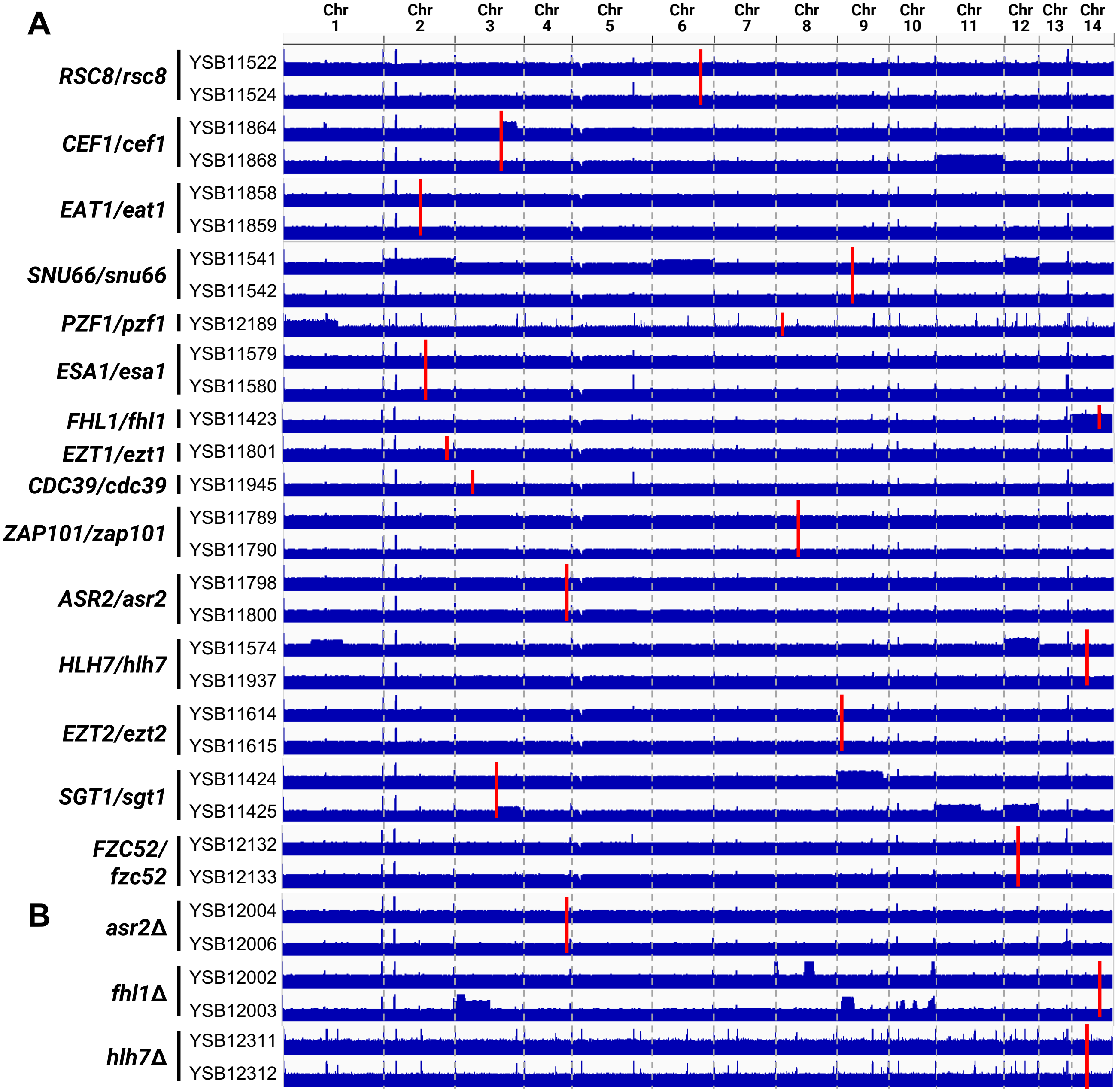


*Continued*

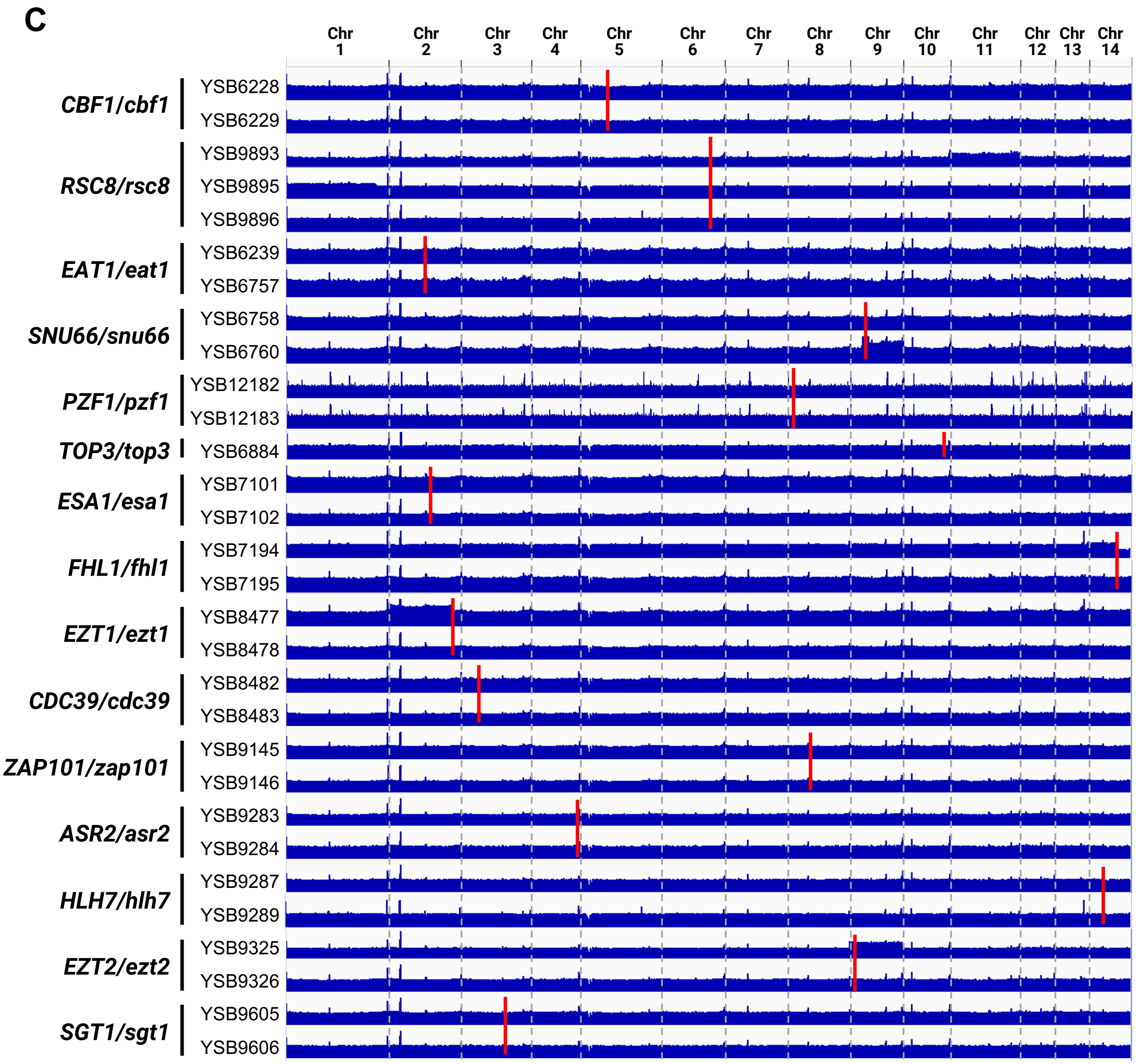


**Supplementary figure 4. Whole-genome sequencing (WGS) coverage plots of (A) heterozygous mutants generated in the CnLC6683 background, (B) knockout progenies derived from the CnLC6683-based heterozygous mutants, and (C) heterozygous mutants constructed in the AI187 background.** Genomic DNA extracted from each strain was subjected to Illumina sequencing, followed by read alignment to the *Cryptococcus neoformans* H99 reference genome using BWA. Alignment files were processed and converted using SAMtools, and genome-wide coverage depth was calculated using a binning size of 100 base pairs. Resulting coverage profiles were visualised using IGV to assess genome integrity, chromosomal CNV patterns, and potential structural variations across strains. In each coverage plot, the chromosomal positions of genes targeted are marked by red bars.

### Supplementary figure 5 (Lee et al.)

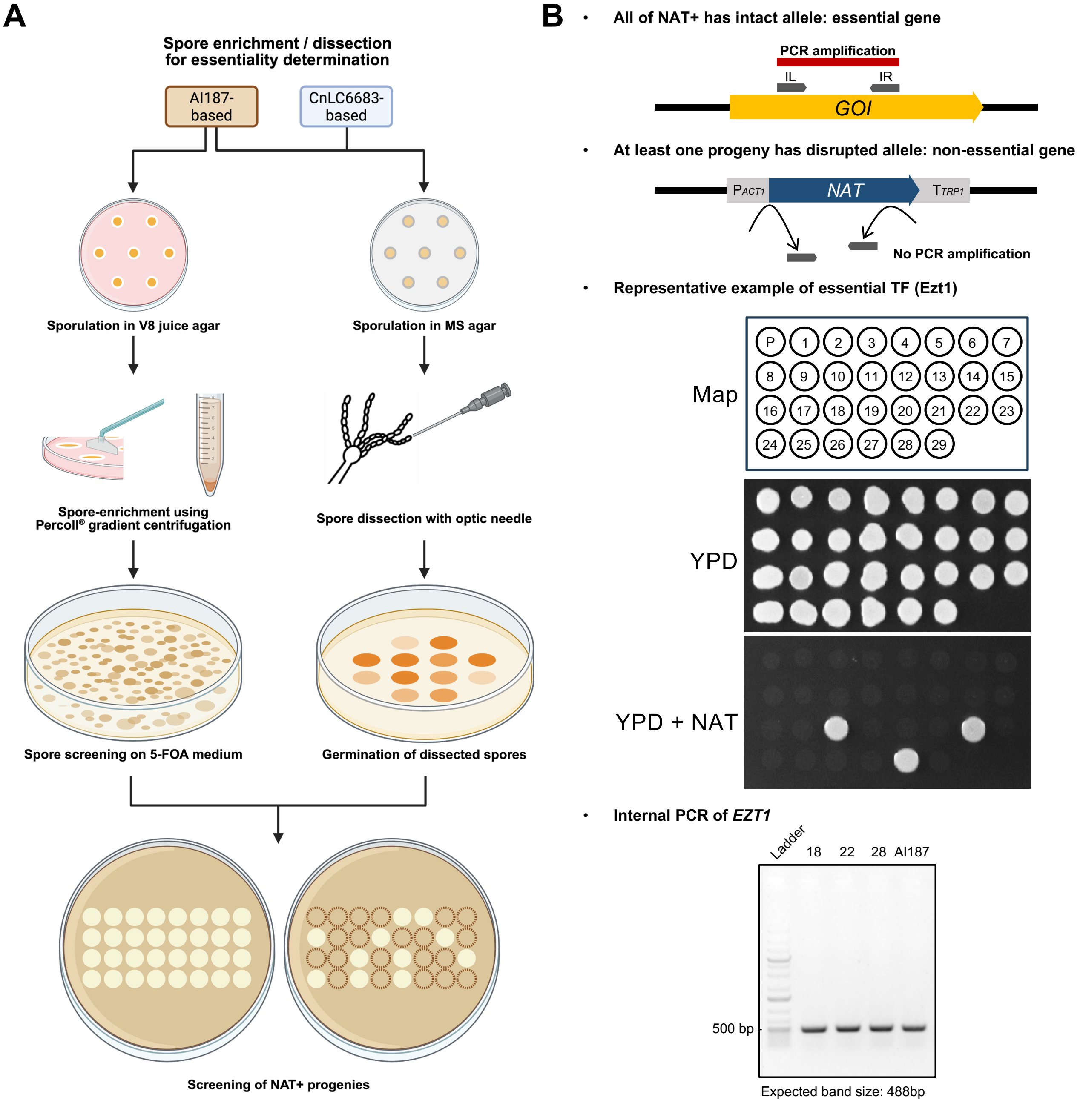


**Supplementary figure 5. Schematic overview and representative example of meiotic spore analysis following spore separation and dissection.** (A) Schematic overview of the spore analysis workflow. Heterozygous mutants generated in the diploid strain AI187 were subjected to spore separation by gradient centrifugation and spore dissection using a micromanipulation needle. Heterozygous mutants generated in the diploid strain CnLC6683 were subjected only to spore dissection because the parental strain lacks auxotrophic markers. After progeny identification by *MAT* locus PCR, progenies were further validated for the presence of the drug resistance marker by spotting on YPD plate containing nourseothricin. The illustration was created with Biorender. (B) Representative example of essentiality validation for *EZT1*. Internal PCR was performed on nourseothricin resistant progeny using the primer pari B6117 and B14784, which anneal within the deletion target region. All essential TFs were validated using the same method shown here.

**Supplementary figure 6 (Lee et al.)**

**A**

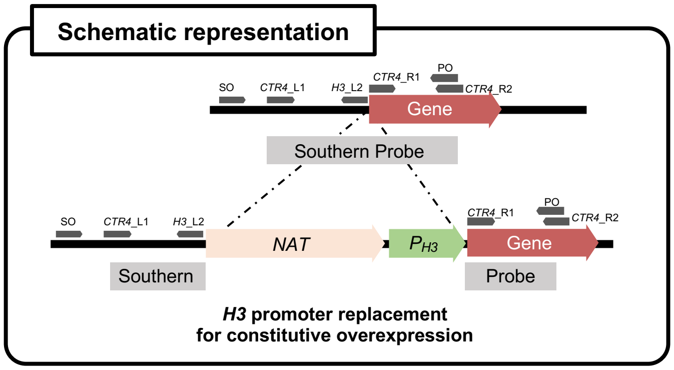


**B**

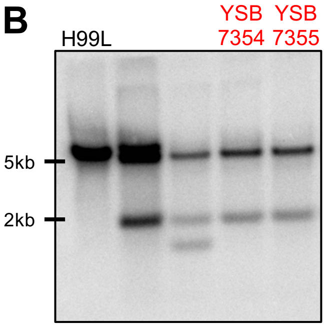


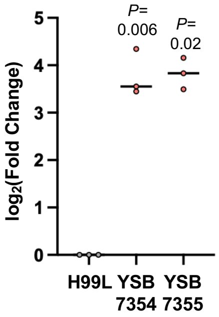


$P_{H3}$:*CBF1* construction
Enzyme digestion: EcoRI
WT: 5857 bp
Mutant: 5613 bp and 2168 bp

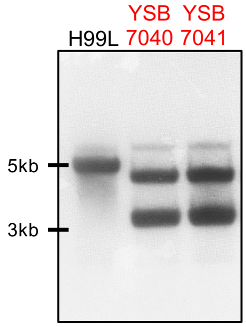


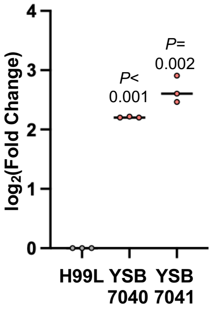


$P_{H3}$:*RSC8* construction
Enzyme digestion: HindIII and XhoI
WT: 5007 bp
Mutant: 4587 bp and 3565 bp

**C**

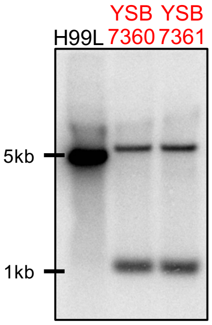


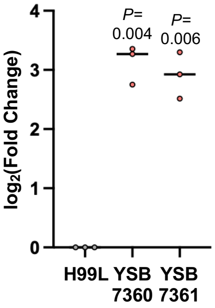


$P_{H3}$:*CEF1* construction
Enzyme digestion: XhoI
WT: 4880 bp
Mutant: 5939 bp and 1086 bp

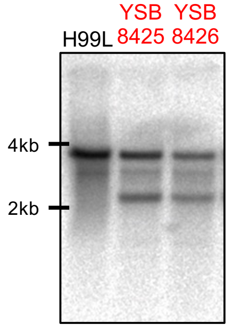


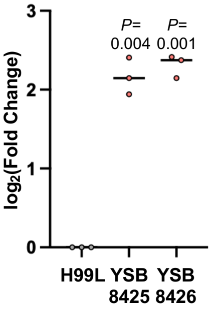


$P_{H3}$:*CDC39* construction
Enzyme digestion: EcoRV
WT: 3869 bp
Mutant: 3846 bp and 2168 bp

***Continued***

**D**

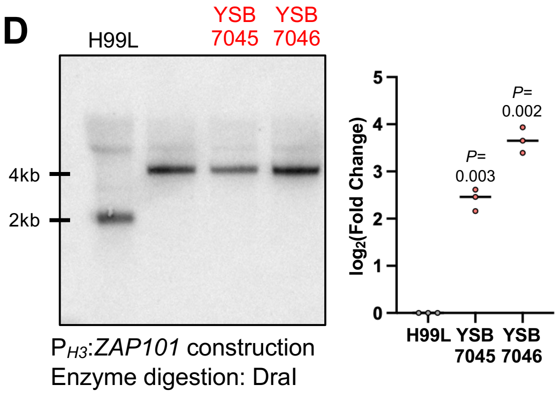


P$_{H3}$:*ZAP101* construction
Enzyme digestion: DraI
WT: 2137 bp
Mutant: 4282 bp

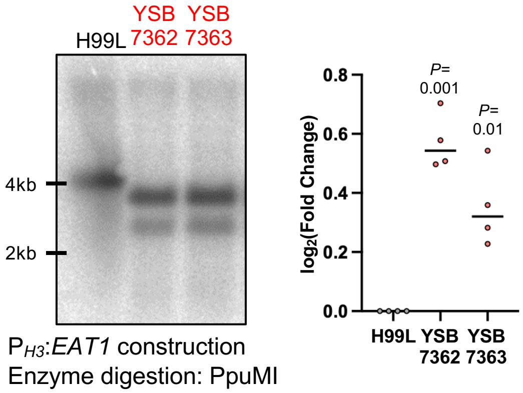


P$_{H3}$:*EAT1* construction
Enzyme digestion: PpuMI
WT: 4196 bp
Mutant: 3581 bp and 2760 bp

**E**

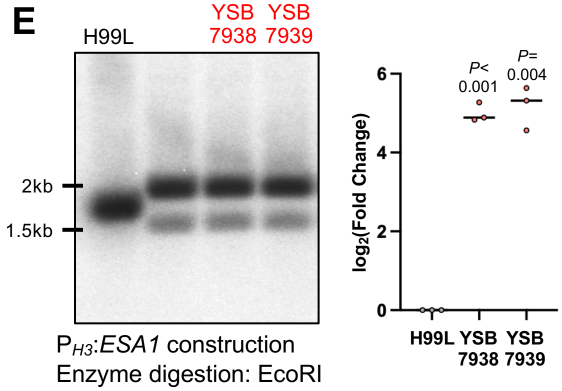


P$_{H3}$:*ESA1* construction
Enzyme digestion: EcoRI
WT: 1840 bp
Mutant: 2186 bp and 1578 bp

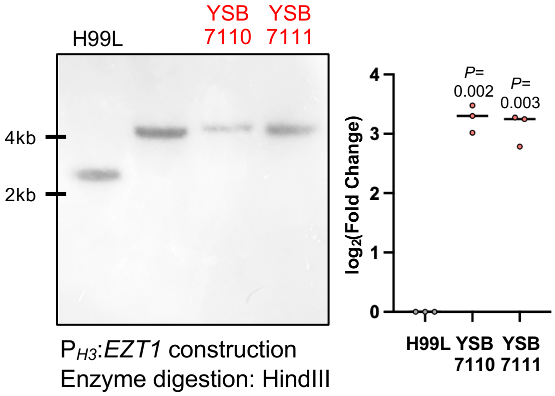


P$_{H3}$:*EZT1* construction
Enzyme digestion: HindIII
WT: 2731 bp
Mutant: 4876 bp

**F**

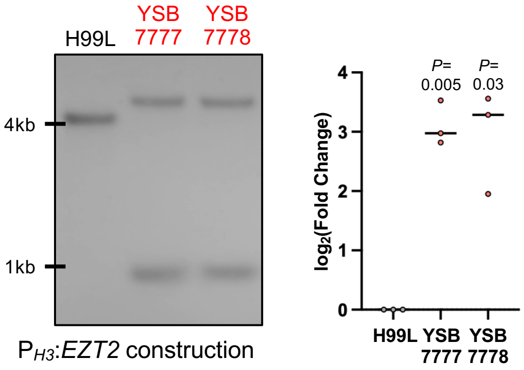


P$_{H3}$:*EZT2* construction
Enzyme digestion: XhoI
WT: 4243 bp
Mutant: 5415 bp and 973 bp

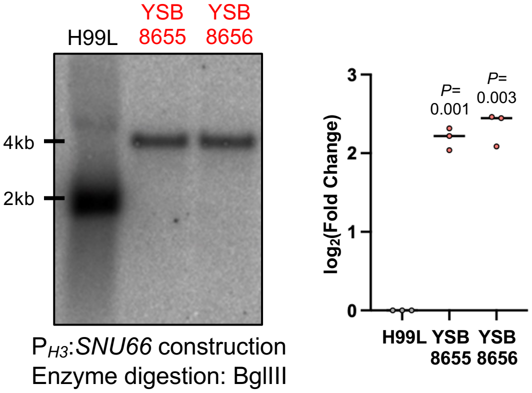


P$_{H3}$:*SNU66* construction
Enzyme digestion: BglIII
WT: 1957 bp
Mutant: 4102 bp

***Continued***

G

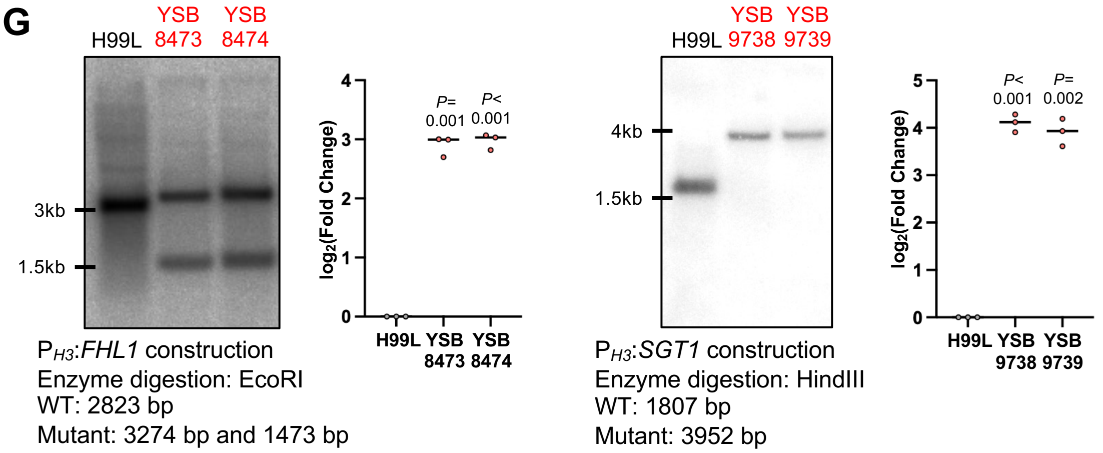


H

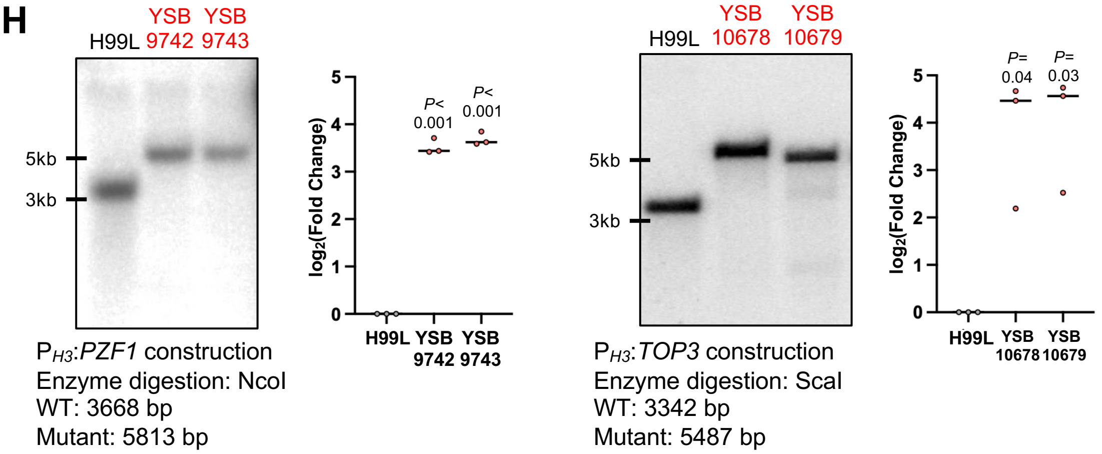


I

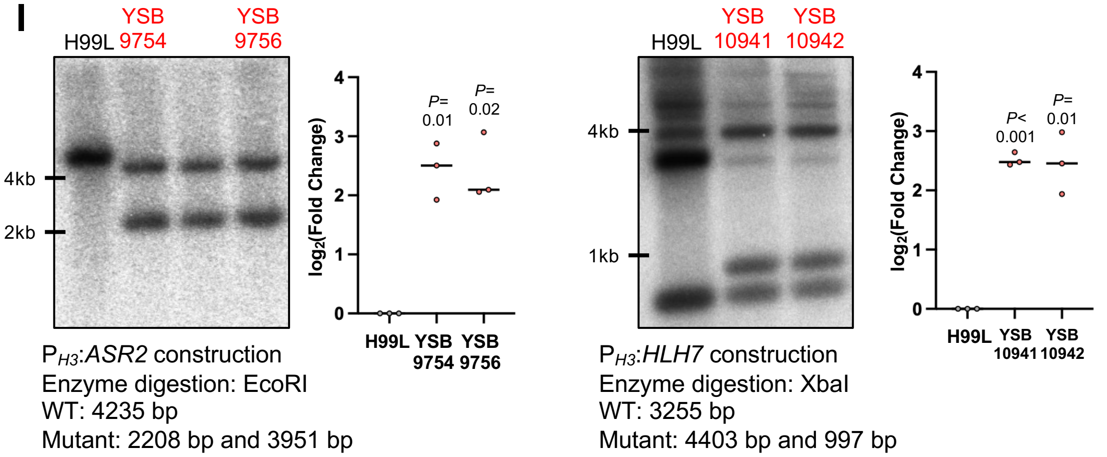


**Supplementary figure 6. Construction of the constitutive overexpression strains.** (A) Schematic representation for the construction of the histone 3 (H3) promoter-derived constitutive overexpression strains. (B-I) Correct genotypes of each overexpression strain were confirmed through Southern blot analysis with the designated restriction enzyme digestion of genomic DNA and gene-specific probes. The overexpression of the targeted genes was confirmed with qRT-PCR. The strains were grown overnight at 30°C in 50 ml of liquid YPD medium, subcultured into 50 ml of fresh YPD medium ($OD_{600nm}$=0.2). Cell cultures were further incubated at 30°C in a shaking incubator until OD600nm reaches between 0.6 and 0.8, harvested, frozen in liquid nitrogen, and lyophilized. Total RNAs were isolated, and cDNAs were synthesized. qRT-PCRs were performed using gene-specific primer pairs and the expression levels of the genes were normalized with *ACT1* expression. Individual data are shown, with horizontal lines indicating the mean. Statistical significance was assessed using a one-sample *t*-test comparing the normalized values of the overexpression strain with the wild-type reference value (=1) in Prism 11.0. *P* values are indicated above the plots.

### Supplementary figure 7 (Lee et al.)

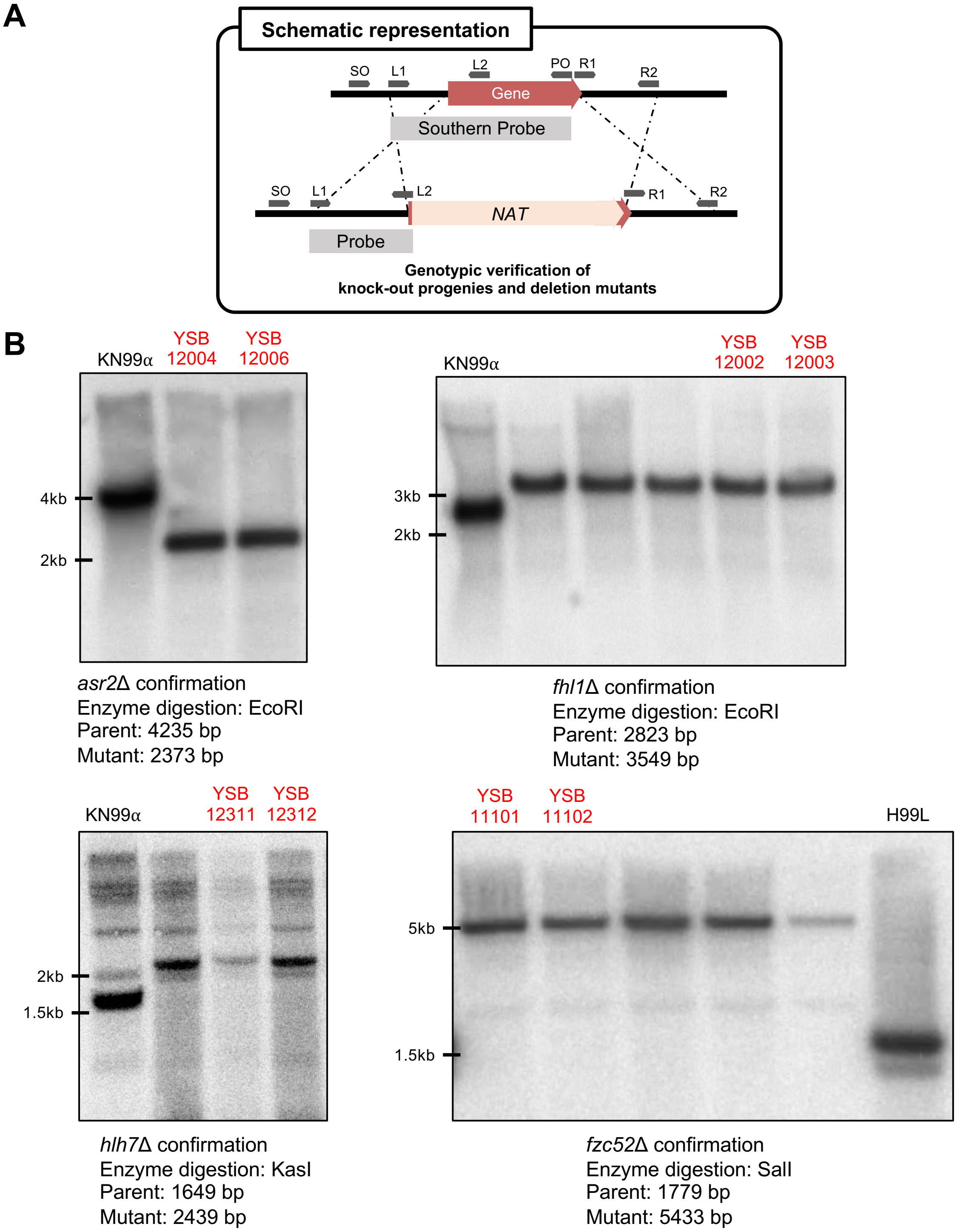


**Supplementary figure 7. Confirmation of the deletion mutants of non-essential TFs (*ASR2*, *FHL1*, *HLH7*, and *FZC52*).** (A) Schematic representation for confirming the haploid knockout strains. (B) The genomic DNA of the wild-types (KN99 for *asr2*Δ spores, *hlh7*Δ spores, and *fhl1*Δ spores, and H99L for *fzc52*Δ), knock-out haploid spores, and mutants were subjected to restriction enzyme digestion and proceeded to Southern blot analysis with the gene-specific probes.

**Supplementary figure 8 (Lee et al.)**

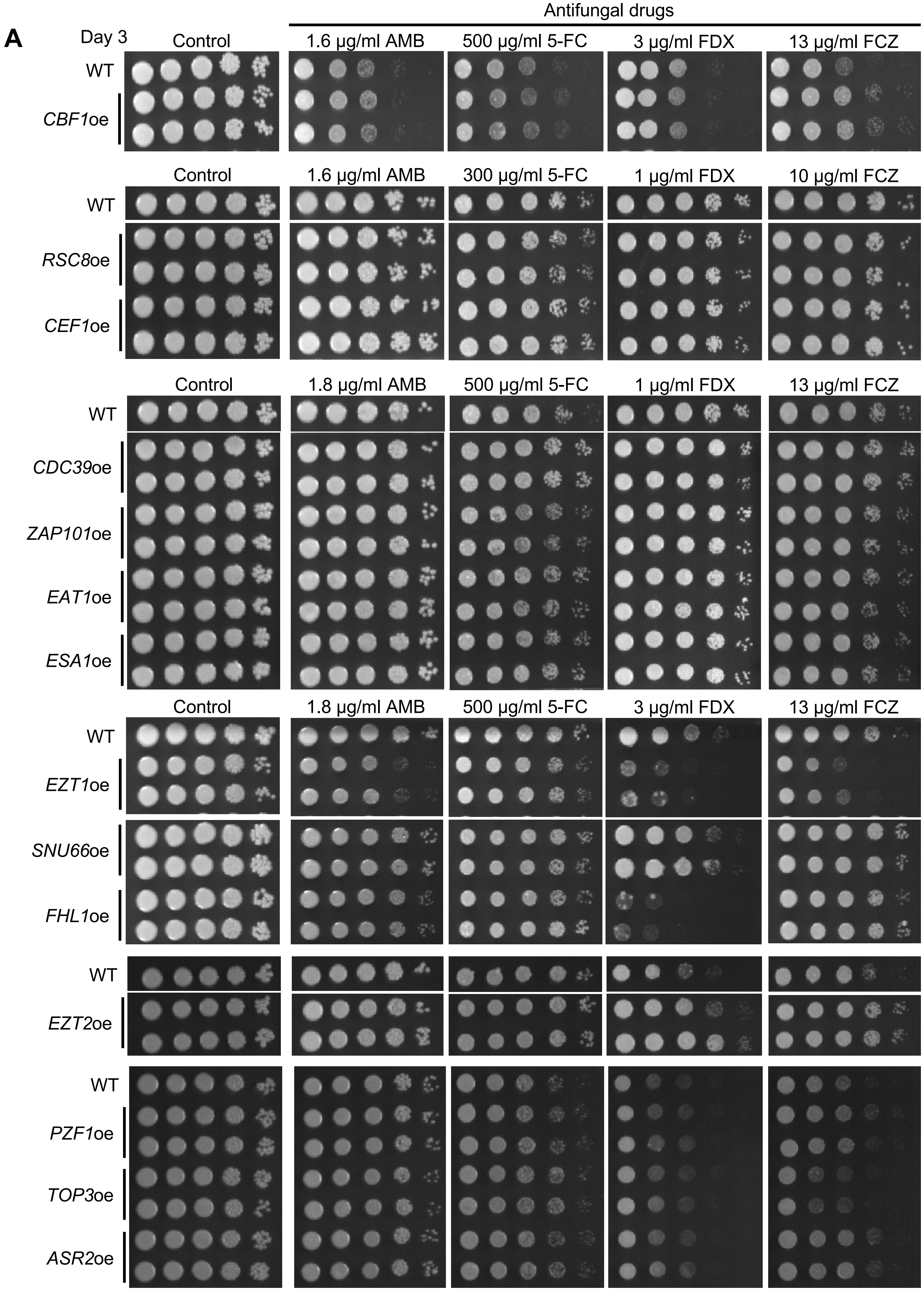


***Continued***

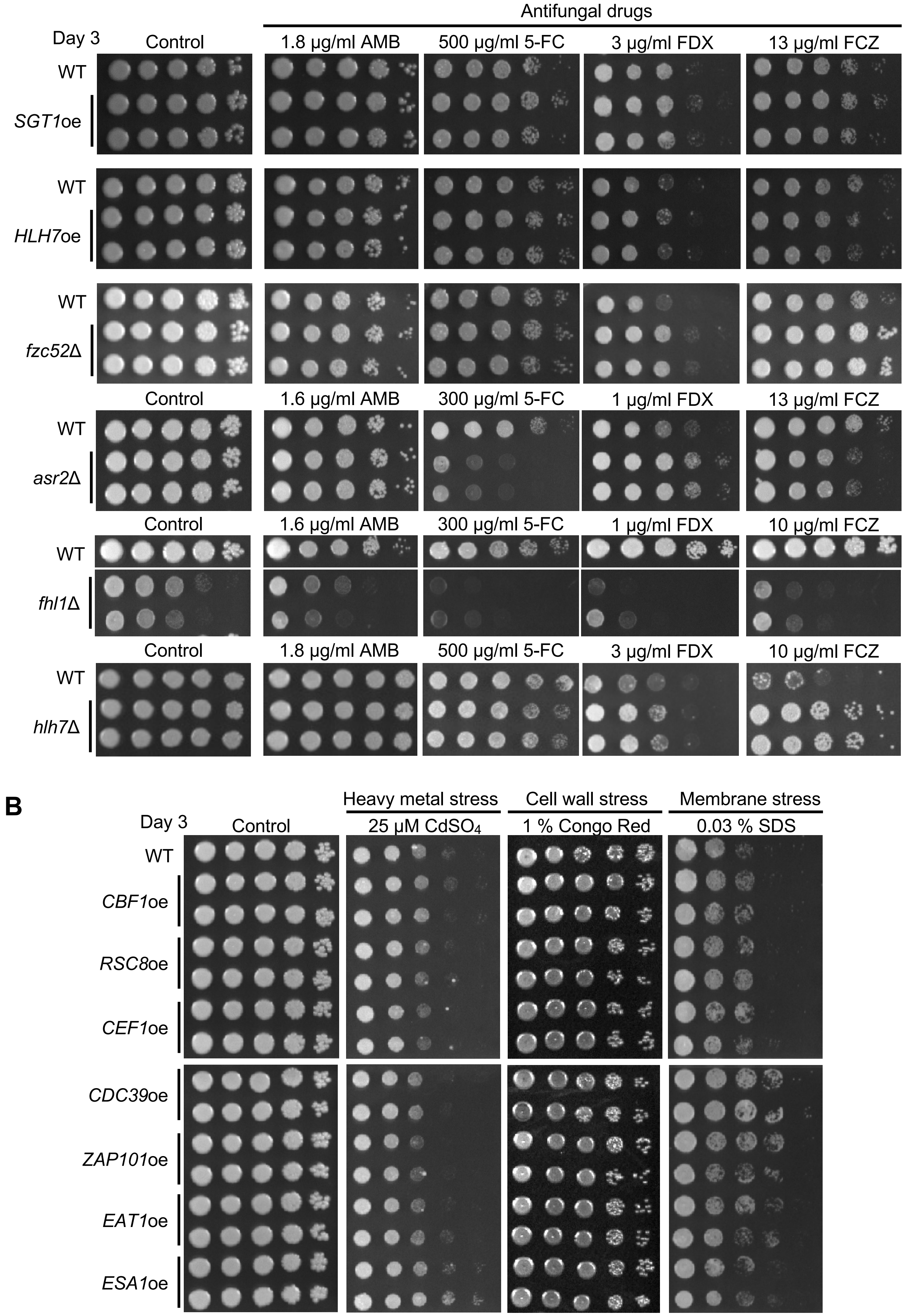
Antifungal drugs
Day 3
Control
1.8 μg/ml AMB
500 μg/ml 5-FC
3 μg/ml FDX
13 μg/ml FCZ
WT
SGT1oe
WT
HLH7oe
WT
fzc52Δ
Control
1.6 μg/ml AMB
300 μg/ml 5-FC
1 μg/ml FDX
13 μg/ml FCZ
WT
asr2Δ
Control
1.6 μg/ml AMB
300 μg/ml 5-FC
1 μg/ml FDX
10 μg/ml FCZ
WT
fhl1Δ
Control
1.8 μg/ml AMB
500 μg/ml 5-FC
3 μg/ml FDX
10 μg/ml FCZ
WT
hlh7Δ
B
Day 3
Control
Heavy metal stress
25 μM CdSO4
Cell wall stress
1 % Congo Red
Membrane stress
0.03 % SDS
WT
CBF1oe
RSC8oe
CEF1oe
CDC39oe
ZAP101oe
EAT1oe
ESA1oe

***Continued***

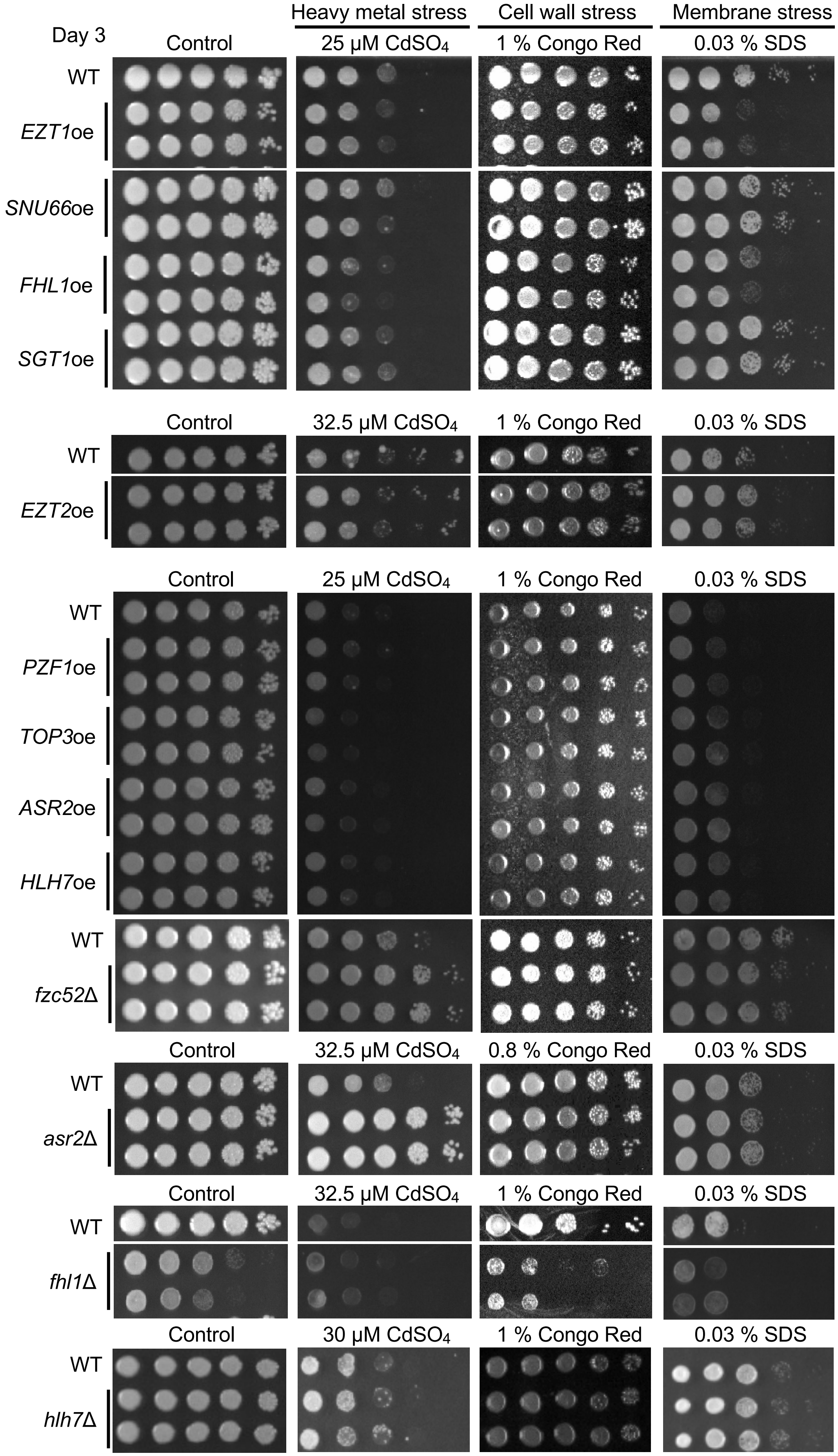
Day 3
Heavy metal stress
Cell wall stress
Membrane stress
Control
25 µM CdSO4
1 % Congo Red
0.03 % SDS
WT
EZT1oe
SNU66oe
FHL1oe
SGT1oe
Control
32.5 µM CdSO4
1 % Congo Red
0.03 % SDS
WT
EZT2oe
Control
25 µM CdSO4
1 % Congo Red
0.03 % SDS
WT
PZF1oe
TOP3oe
ASR2oe
HLH7oe
WT
fzc52Δ
Control
32.5 µM CdSO4
0.8 % Congo Red
0.03 % SDS
WT
asr2Δ
Control
32.5 µM CdSO4
1 % Congo Red
0.03 % SDS
WT
fhl1Δ
Control
30 µM CdSO4
1 % Congo Red
0.03 % SDS
WT
hlh7Δ

***Continued***

C

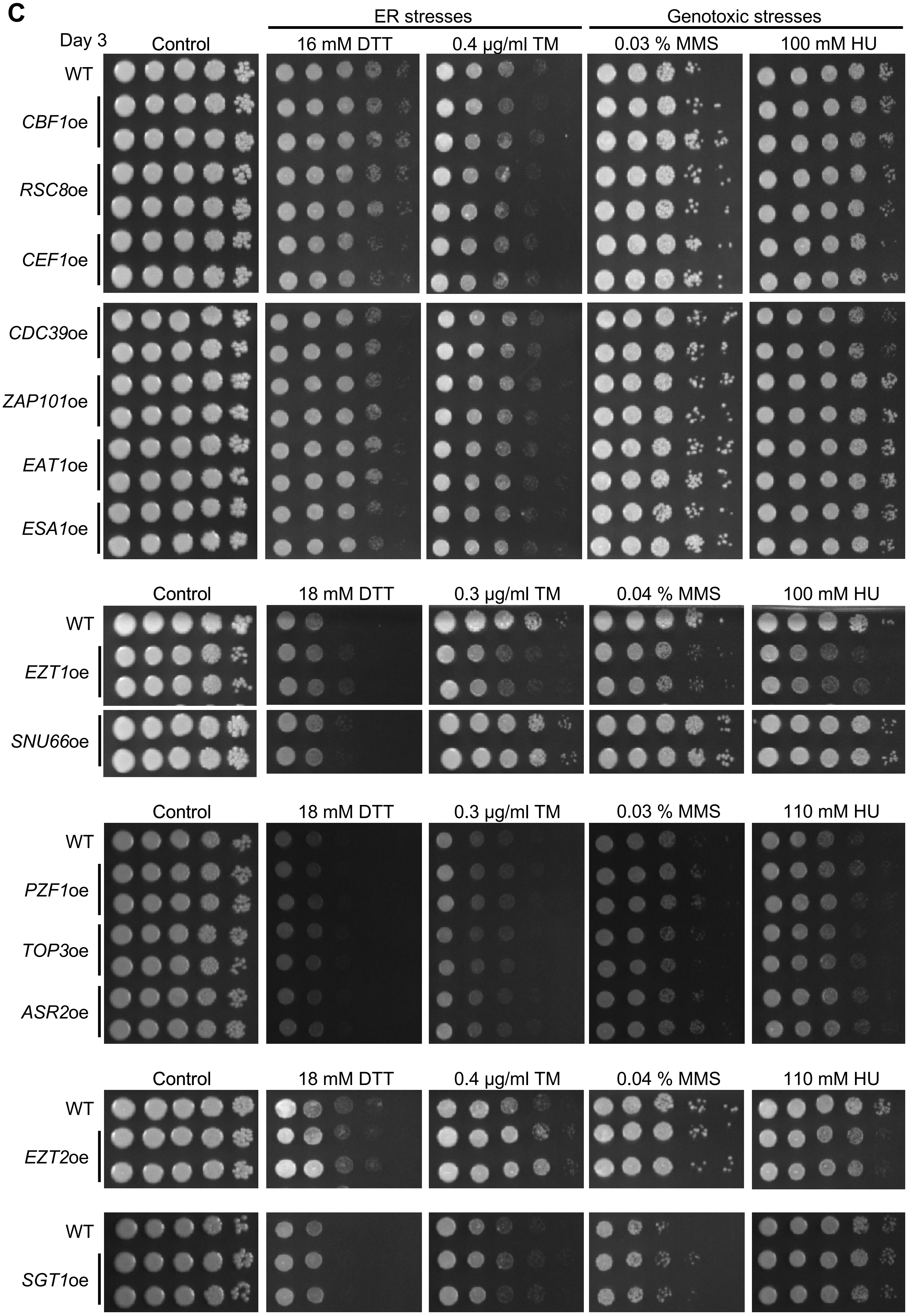
ER stresses
Genotoxic stresses
Day 3
Control
16 mM DTT
0.4 μg/ml TM
0.03 % MMS
100 mM HU
WT
CBF1oe
RSC8oe
CEF1oe
CDC39oe
ZAP101oe
EAT1oe
ESA1oe
Control
18 mM DTT
0.3 μg/ml TM
0.04 % MMS
100 mM HU
WT
EZT1oe
SNU66oe
Control
18 mM DTT
0.3 μg/ml TM
0.03 % MMS
110 mM HU
WT
PZF1oe
TOP3oe
ASR2oe
Control
18 mM DTT
0.4 μg/ml TM
0.04 % MMS
110 mM HU
WT
EZT2oe
WT
SGT1oe

***Continued***

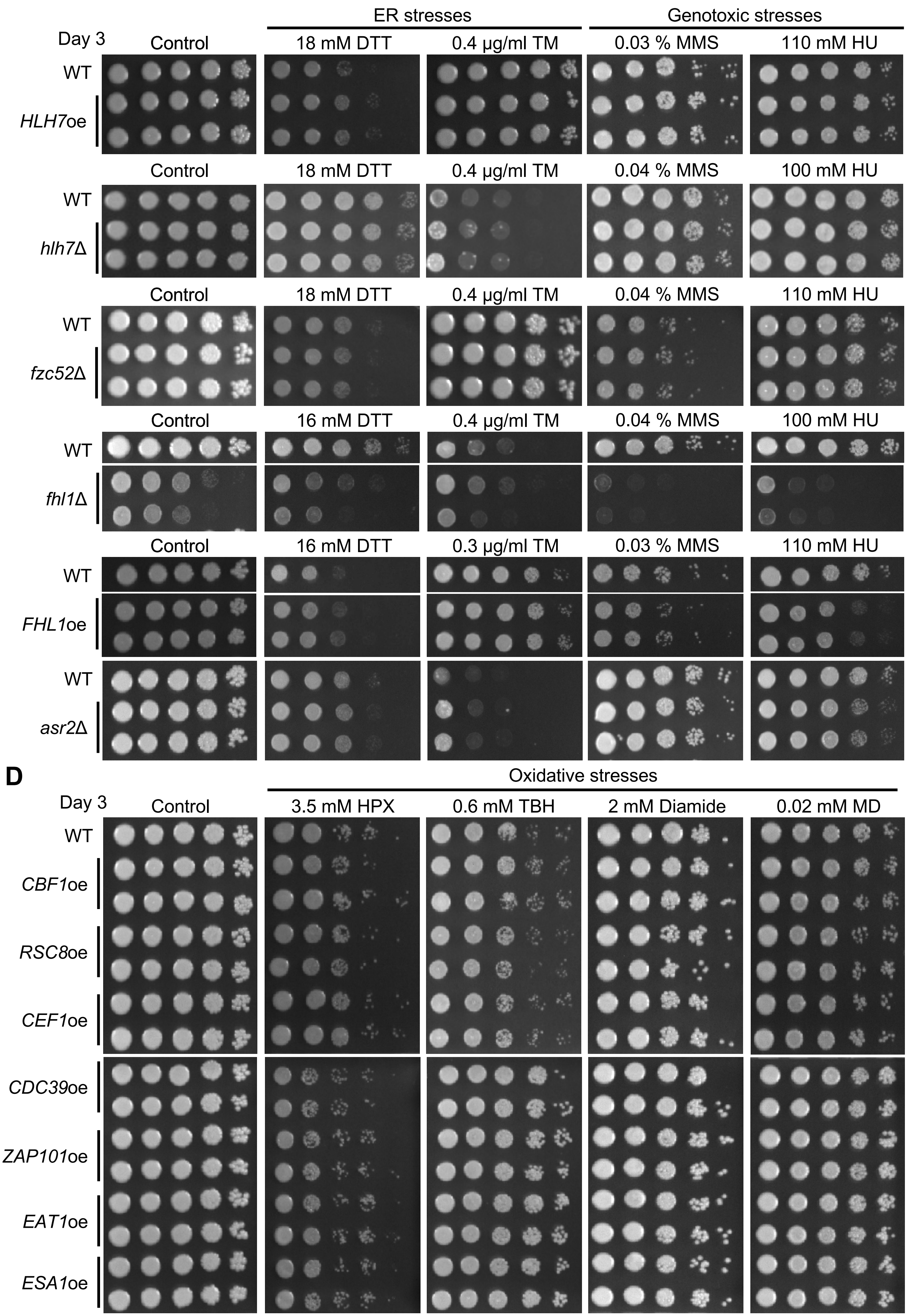
ER stresses
Genotoxic stresses
Day 3
Control
18 mM DTT
0.4 µg/ml TM
0.03 % MMS
110 mM HU
WT
HLH7oe
0.04 % MMS
100 mM HU
hlh7Δ
fzc52Δ
16 mM DTT
fhl1Δ
0.3 µg/ml TM
FHL1oe
asr2Δ
D
Oxidative stresses
3.5 mM HPX
0.6 mM TBH
2 mM Diamide
0.02 mM MD
CBF1oe
RSC8oe
CEF1oe
CDC39oe
ZAP101oe
EAT1oe
ESA1oe

***Continued***

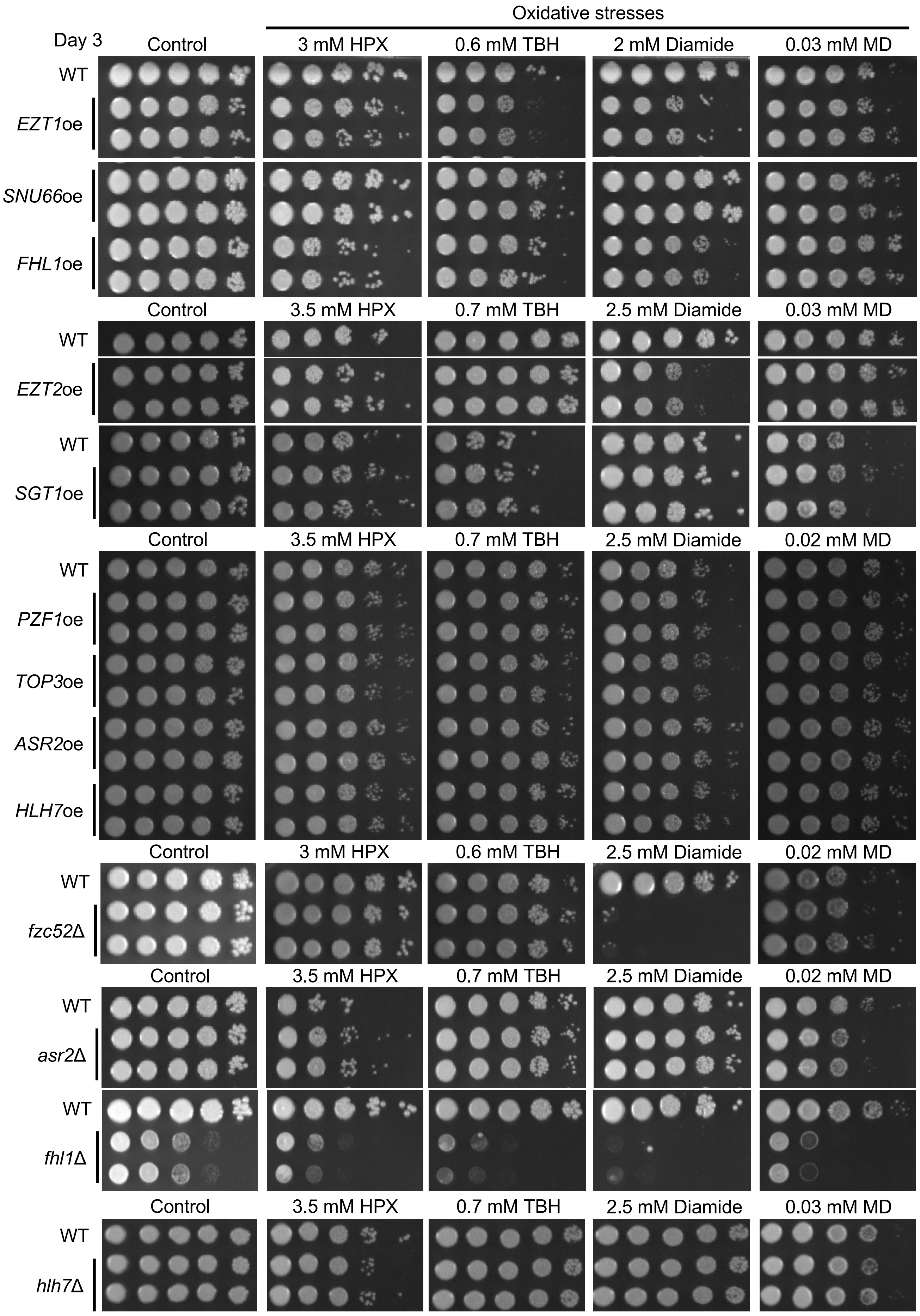
Oxidative stresses
Day 3
Control
3 mM HPX
0.6 mM TBH
2 mM Diamide
0.03 mM MD
WT
EZT1oe
SNU66oe
FHL1oe
Control
3.5 mM HPX
0.7 mM TBH
2.5 mM Diamide
0.03 mM MD
WT
EZT2oe
WT
SGT1oe
Control
3.5 mM HPX
0.7 mM TBH
2.5 mM Diamide
0.02 mM MD
WT
PZF1oe
TOP3oe
ASR2oe
HLH7oe
Control
3 mM HPX
0.6 mM TBH
2.5 mM Diamide
0.02 mM MD
WT
fzc52Δ
Control
3.5 mM HPX
0.7 mM TBH
2.5 mM Diamide
0.02 mM MD
WT
asr2Δ
WT
fhl1Δ
Control
3.5 mM HPX
0.7 mM TBH
2.5 mM Diamide
0.03 mM MD
WT
hlh7Δ

***Continued***

E

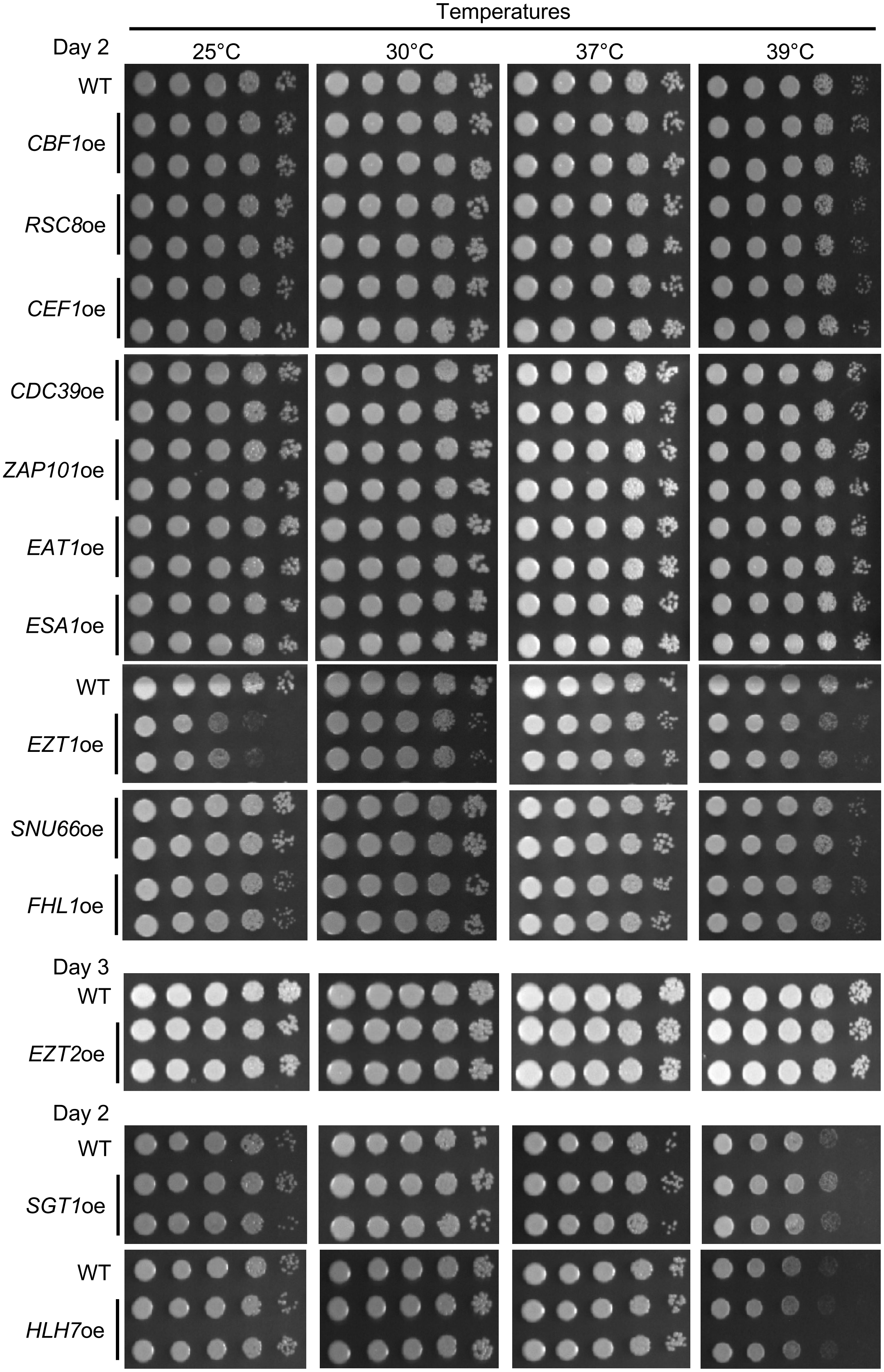

Temperatures
Day 2
25°C
30°C
37°C
39°C
WT
CBF1oe
RSC8oe
CEF1oe
CDC39oe
ZAP101oe
EAT1oe
ESA1oe
WT
EZT1oe
SNU66oe
FHL1oe
Day 3
WT
EZT2oe
Day 2
WT
SGT1oe
WT
HLH7oe


Continued

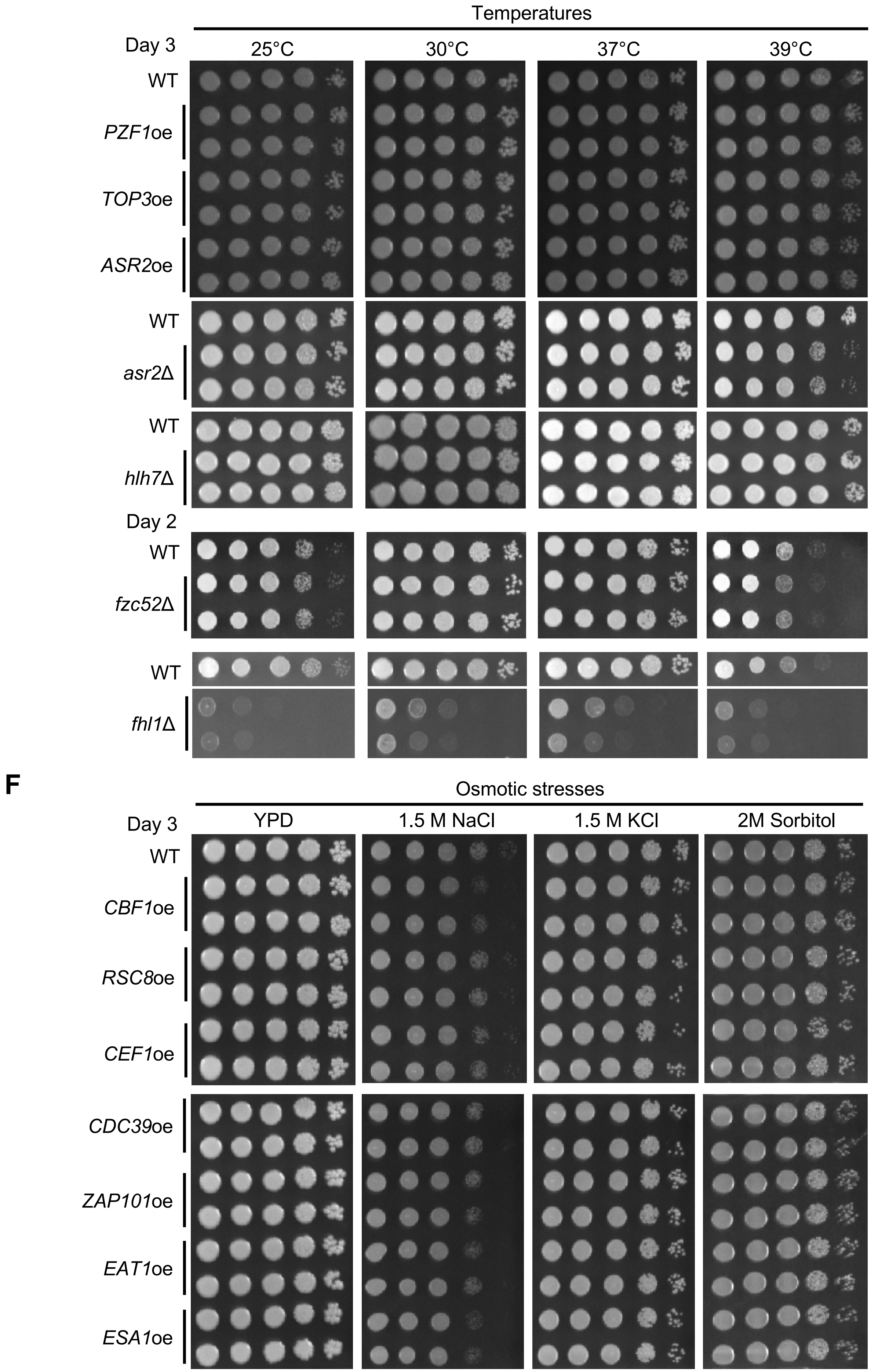
Temperatures
Day 3
25°C
30°C
37°C
39°C
WT
PZF1oe
TOP3oe
ASR2oe
WT
asr2Δ
WT
hlh7Δ
Day 2
WT
fzc52Δ
WT
fhl1Δ
F
Osmotic stresses
Day 3
YPD
1.5 M NaCl
1.5 M KCl
2M Sorbitol
WT
CBF1oe
RSC8oe
CEF1oe
CDC39oe
ZAP101oe
EAT1oe
ESA1oe

***Continued***

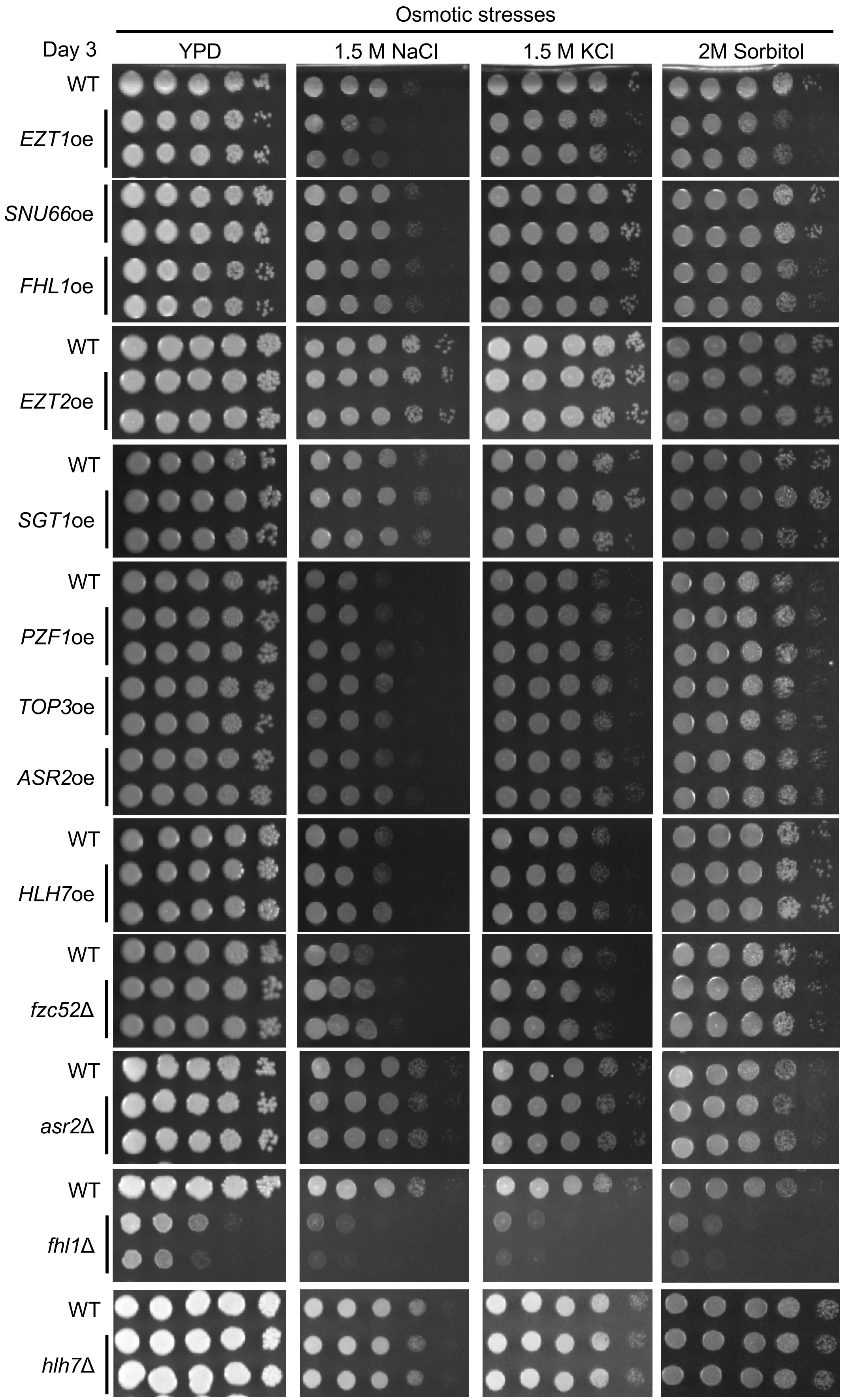
Osmotic stresses
Day 3
YPD
1.5 M NaCl
1.5 M KCl
2M Sorbitol
WT
EZT1oe
SNU66oe
FHL1oe
WT
EZT2oe
WT
SGT1oe
WT
PZF1oe
TOP3oe
ASR2oe
WT
HLH7oe
WT
fzc52Δ
WT
asr2Δ
WT
fhl1Δ
WT
hlh7Δ

***Continued***

G

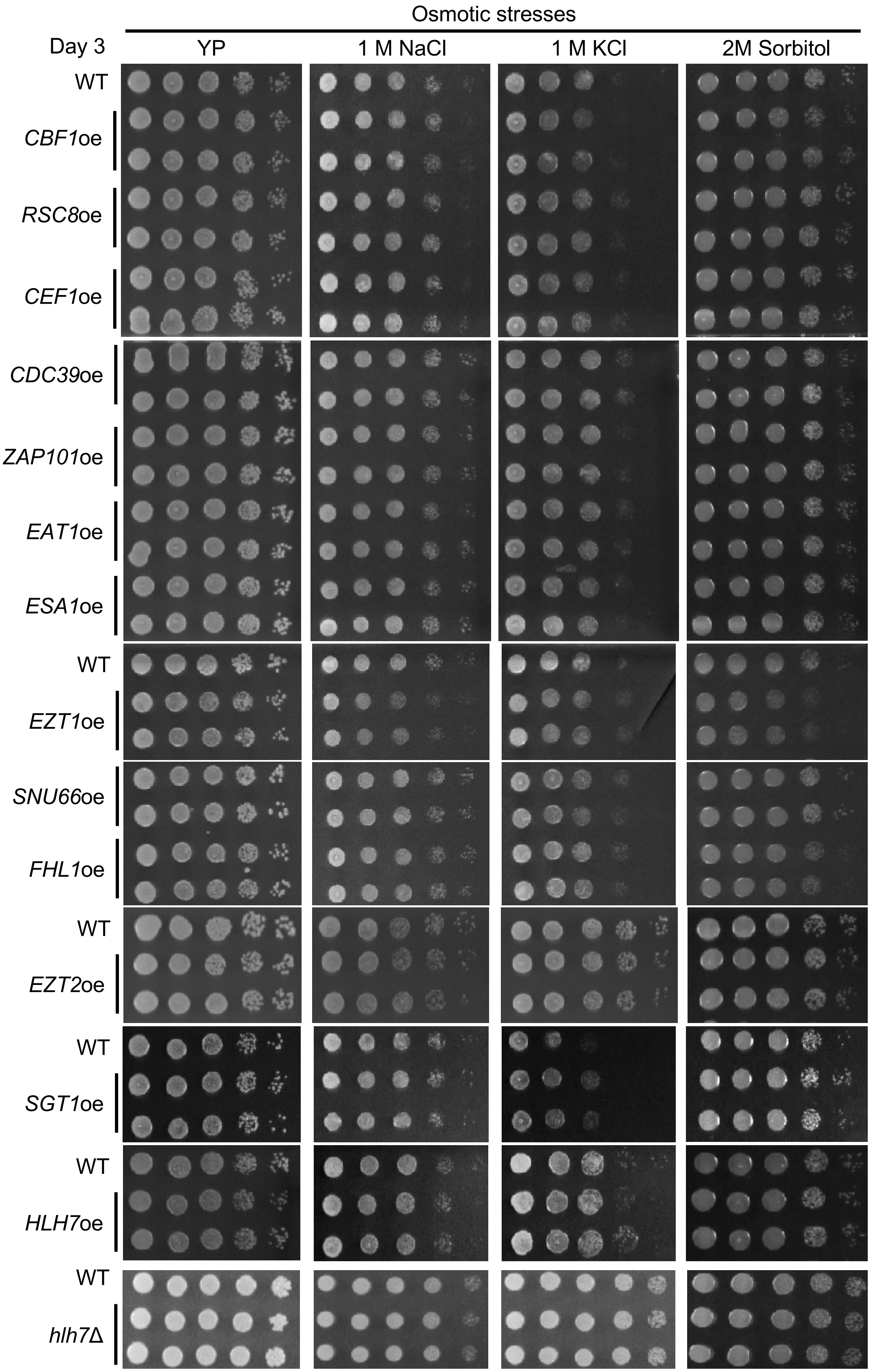
Osmotic stresses
Day 3
YP
1 M NaCl
1 M KCl
2M Sorbitol
WT
CBF1oe
RSC8oe
CEF1oe
CDC39oe
ZAP101oe
EAT1oe
ESA1oe
WT
EZT1oe
SNU66oe
FHL1oe
WT
EZT2oe
WT
SGT1oe
WT
HLH7oe
WT
hlh7Δ

***Continued***

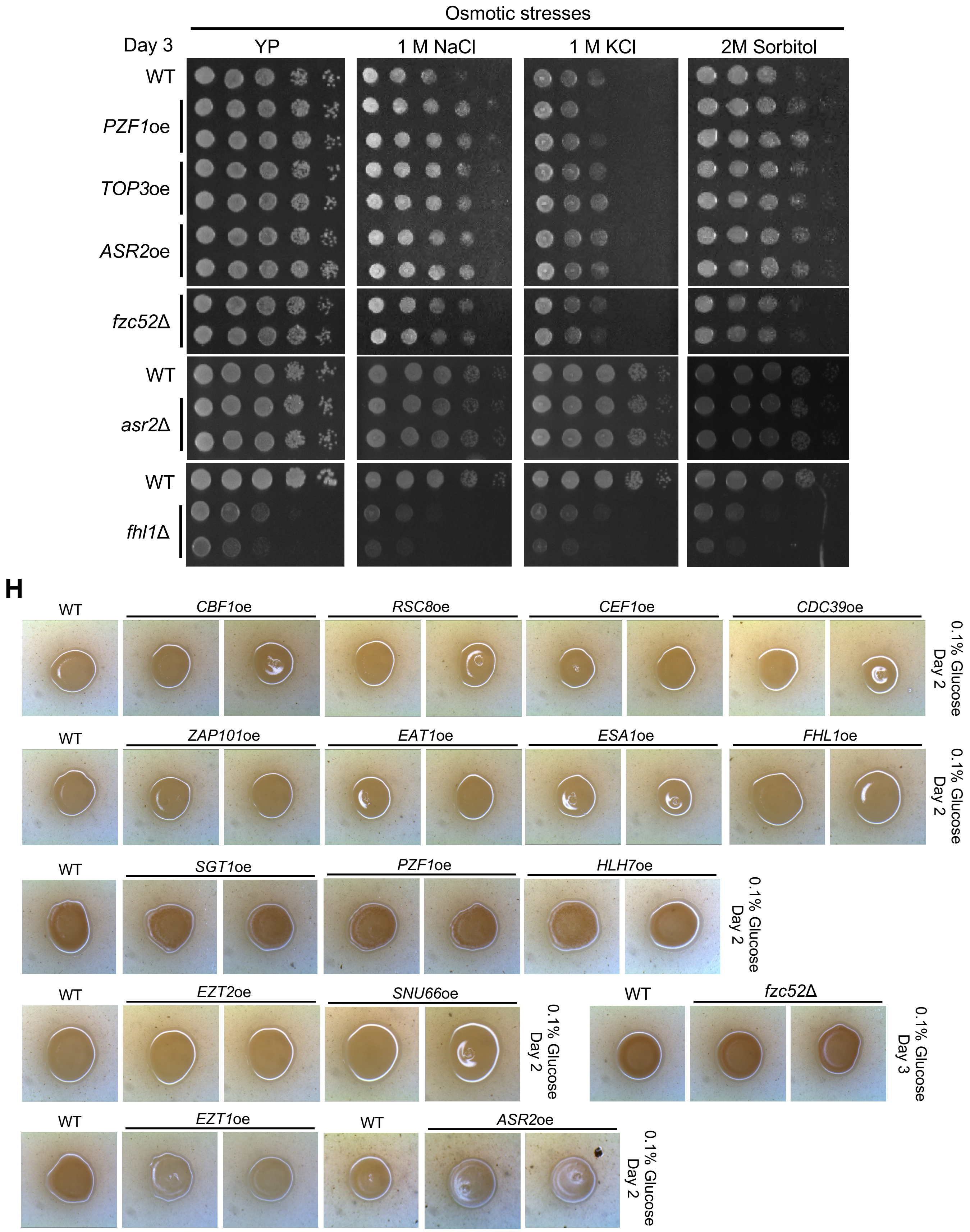
Osmotic stresses
Day 3
YP
1 M NaCl
1 M KCl
2M Sorbitol
WT
PZF1oe
TOP3oe
ASR2oe
fzc52Δ
WT
asr2Δ
WT
fhl1Δ
H
WT
CBF1oe
RSC8oe
CEF1oe
CDC39oe
0.1% Glucose Day 2
WT
ZAP101oe
EAT1oe
ESA1oe
FHL1oe
0.1% Glucose Day 2
WT
SGT1oe
PZF1oe
HLH7oe
0.1% Glucose Day 2
WT
EZT2oe
SNU66oe
0.1% Glucose Day 2
WT
fzc52Δ
0.1% Glucose Day 3
WT
EZT1oe
WT
ASR2oe
0.1% Glucose Day 2

**Continued**

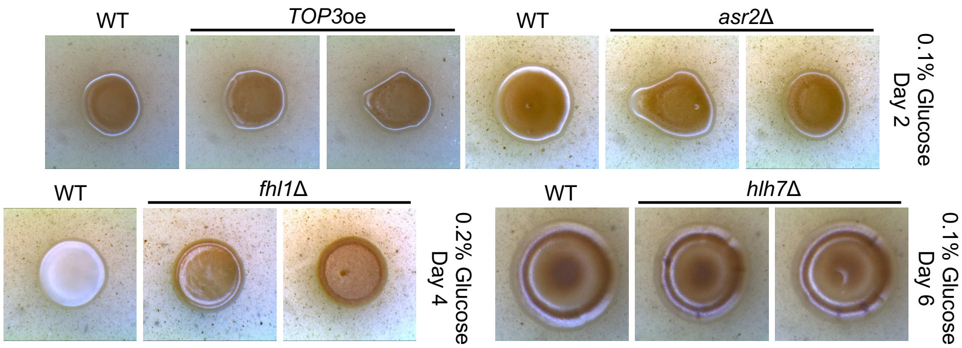


I

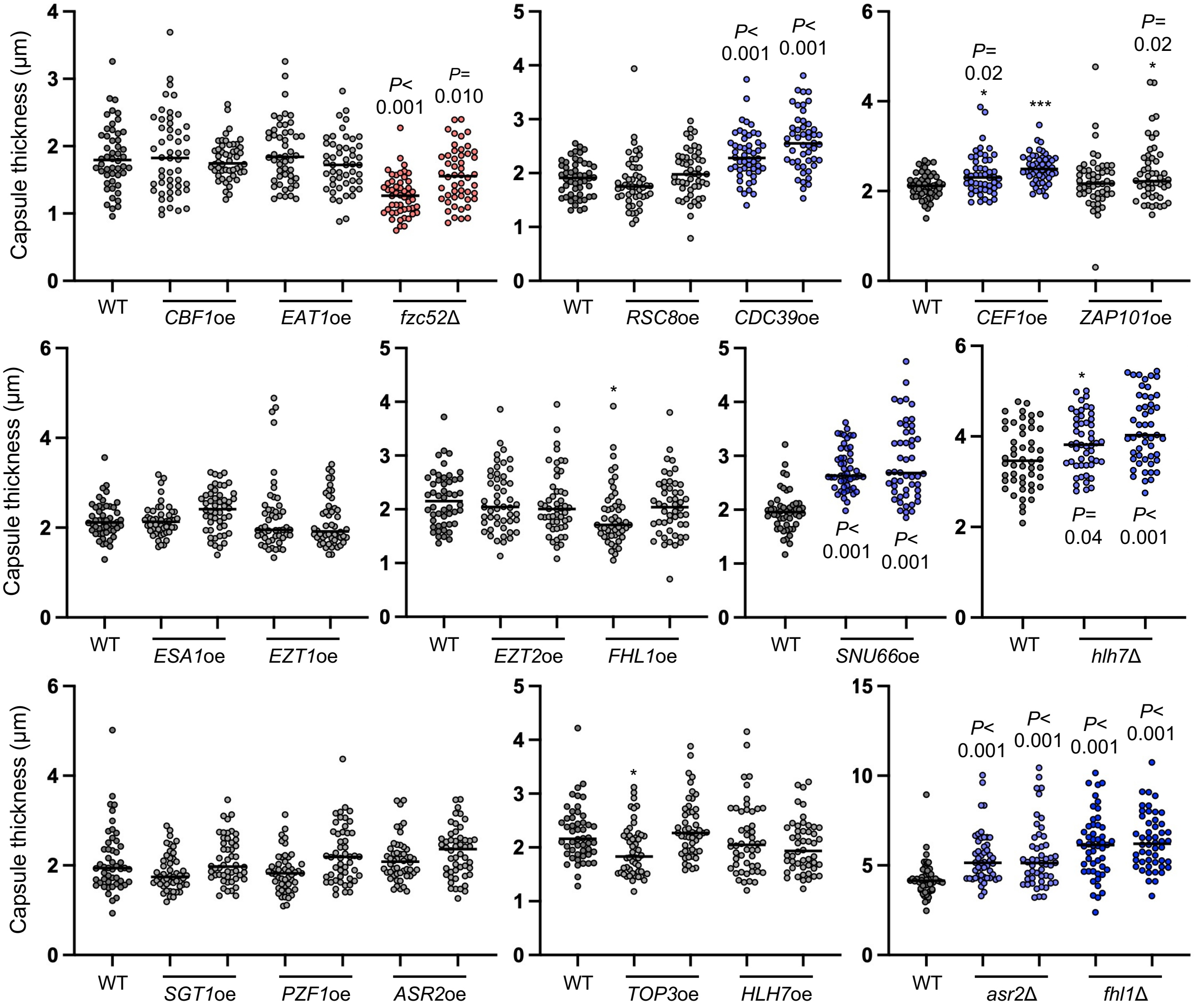


J

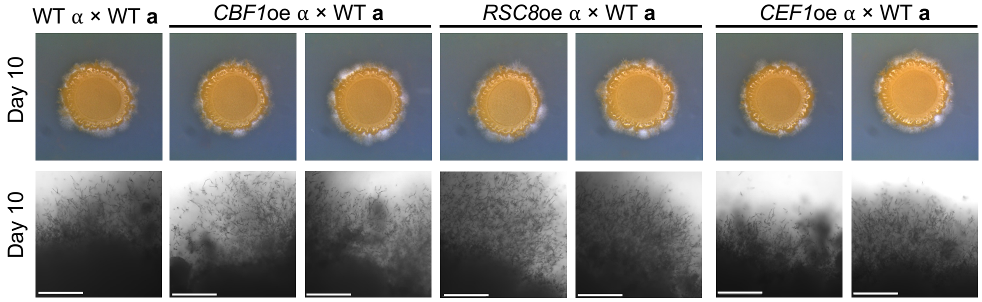


**Continued**

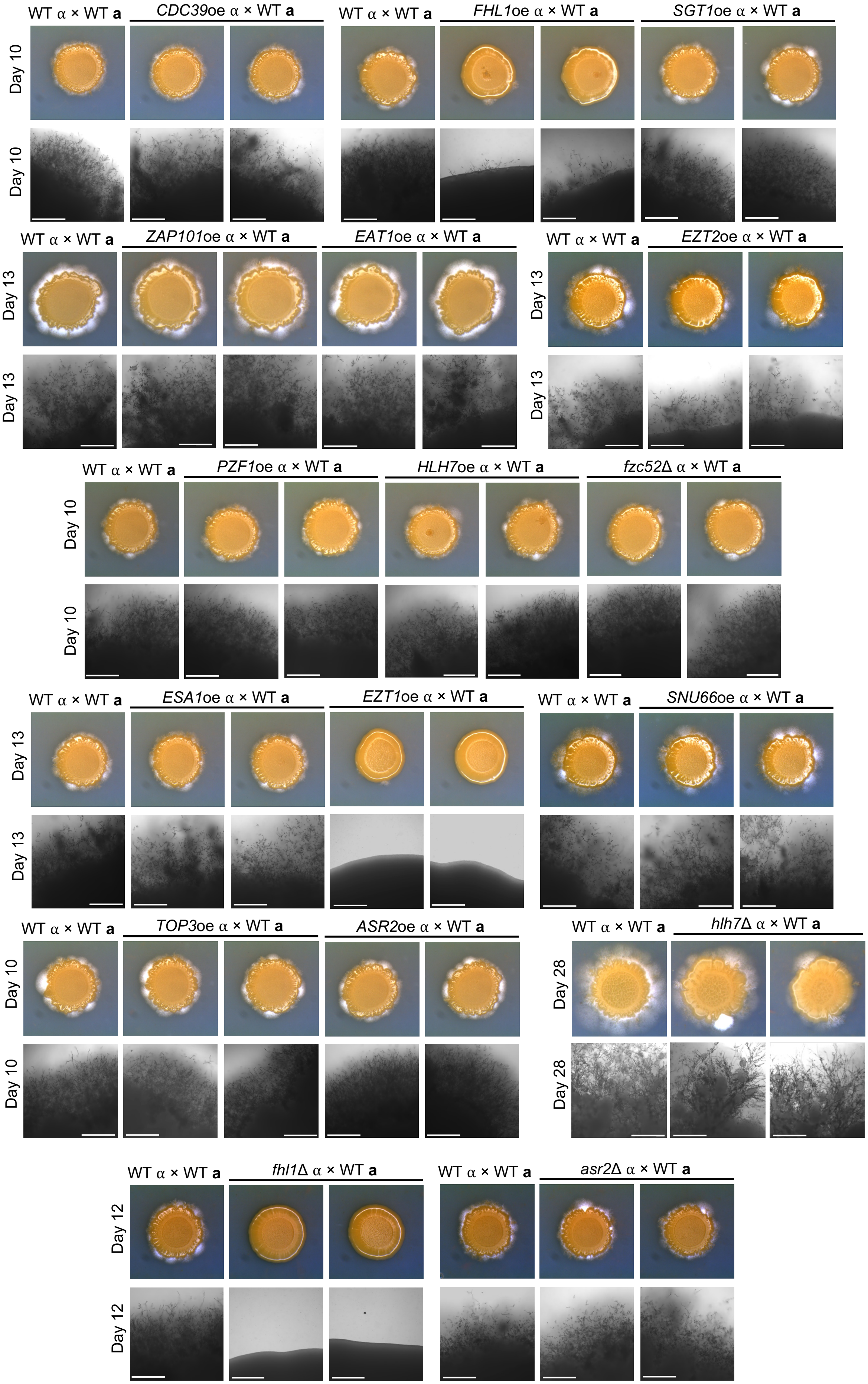


**Figure legend is described on the next page.**

**Supplementary figure 8. Phenotypic analysis of the overexpression strains was conducted through various stress and antifungal drug treatments, virulence factor induction assays, and mating experiments.** (A-G) The wild-type and the overexpression strains were cultured overnight at 30°C in liquid YPD medium, serially diluted (1 to $10^4$), and spotted onto the YPD plates (3 µl) containing the indicated amount of stress or antifungal agents. Plates were incubated at 30°C (or designated temperature) and photographed after indicated days. (H and I) For virulence factor production assays, strains were grown in liquid YPD, washed twice with PBS, and 3 µl aliquots were spotted on 35 g/l Niger seed media with designated concentration of glucose (melanin induction assay) and DME media (capsule induction assays). Melanin induction was observed at 37°C and photographed at the indicated days with BX41 microscope (Olympus, Japan) with a SPOT digital camera (Diagnostic Instruments, Inc. USA). DME media plate was then incubated at 37°C for 2 days, and India ink staining was performed to visualise the capsules under a microscope. Capsule sizes were quantified by measuring both cell and capsule diameters using Nikon NIS software. Individual data points are shown, with the median indicated by a line. Statistical significances of differences between wild-type and the overexpression strain were determined by one-way ANOVA with Dunnett's multiple comparison test using Prism 11.0; *P* values are indicated above. (J) Mating filaments were observed after mating of the overexpression strains (*MAT*α) with wild-type YL99 *MAT* **a** and photographed at the indicated days using a differential interference contrast microscope ECLIPSE Ni (Nikon, Japan) equipped with DS-QI2 camera (scale bar: 400 µm) and BX51 microscope with a SPOT digital camera software. Each assay was repeated two or three times independently, and a representative image or dataset is shown here. Duplicated control images and data indicated that the same batch of wild-type and mutant strains were used in the indicated experiments and are presented here for comparison purposes.

### Supplementary figure 9 (Lee et al.)

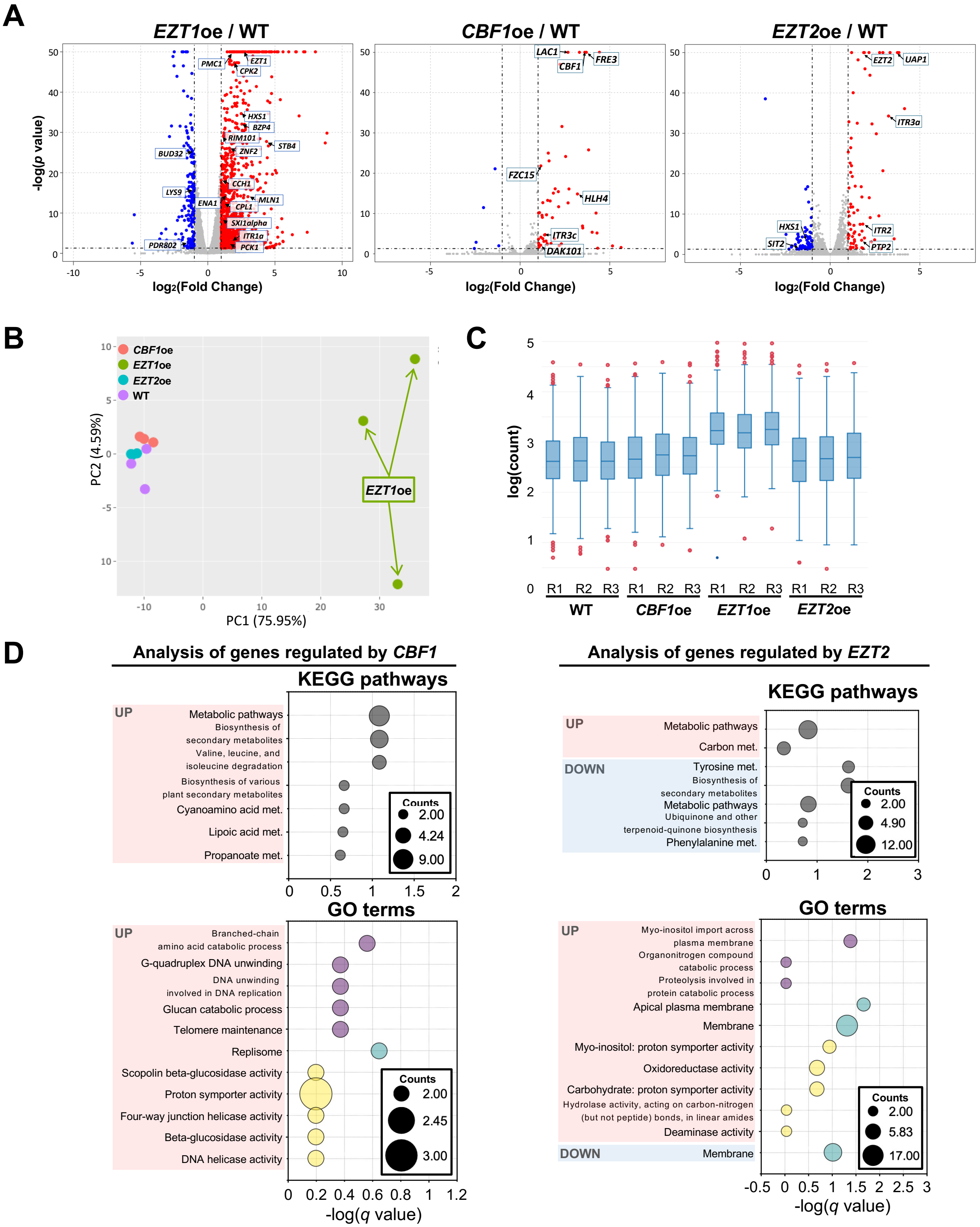


**Supplementary figure 9. Transcriptome analysis of *EZT1*oe, *CBF1*oe, and *EZT2*oe.** (A) Volcano plots showing differentially expressed genes (DEGs) in *EZT1*oe versus wild-type strains (left panel), *CBF1*oe versus wild-type strains (middle panel), and *EZT2*oe versus wild-type strains (right panel). The x-axis represents the $\log_2$ fold change (fc), and the y-axis represents the -$\log_{10}$(p-value). Genes with significant differential expression ($|\log_2 fc| > 1$ and adjusted $p$ value $< 0.05$) are highlighted (red dots indicate upregulated genes, blue dots indicate downregulated genes, and gray dots indicate none-significant genes). (B) The principal component analysis (PCA) plot illustrating the extensive transcriptomic divergence caused by *EZT1*oe compared to *CBF1*oe and *EZT2*oe. (C) Interquartile range (IQR) analysis highlighting more pronounced transcriptomic changes in *EZT1*oe compared to the limited effects in *CBF1*oe and *EZT2*oe (R=biological replicate). (D) KEGG (Kyoto Encyclopedia of Genes and Genomes) analysis and GO (Gene Ontology) term analysis of the significantly regulated genes by the overexpression of *CBF1* (left panel) and *EZT2* (right panel) identified significantly overrepresented biological processes and regulated metabolic and signalling pathways.

Supplementary figure 10 (Lee et al.)

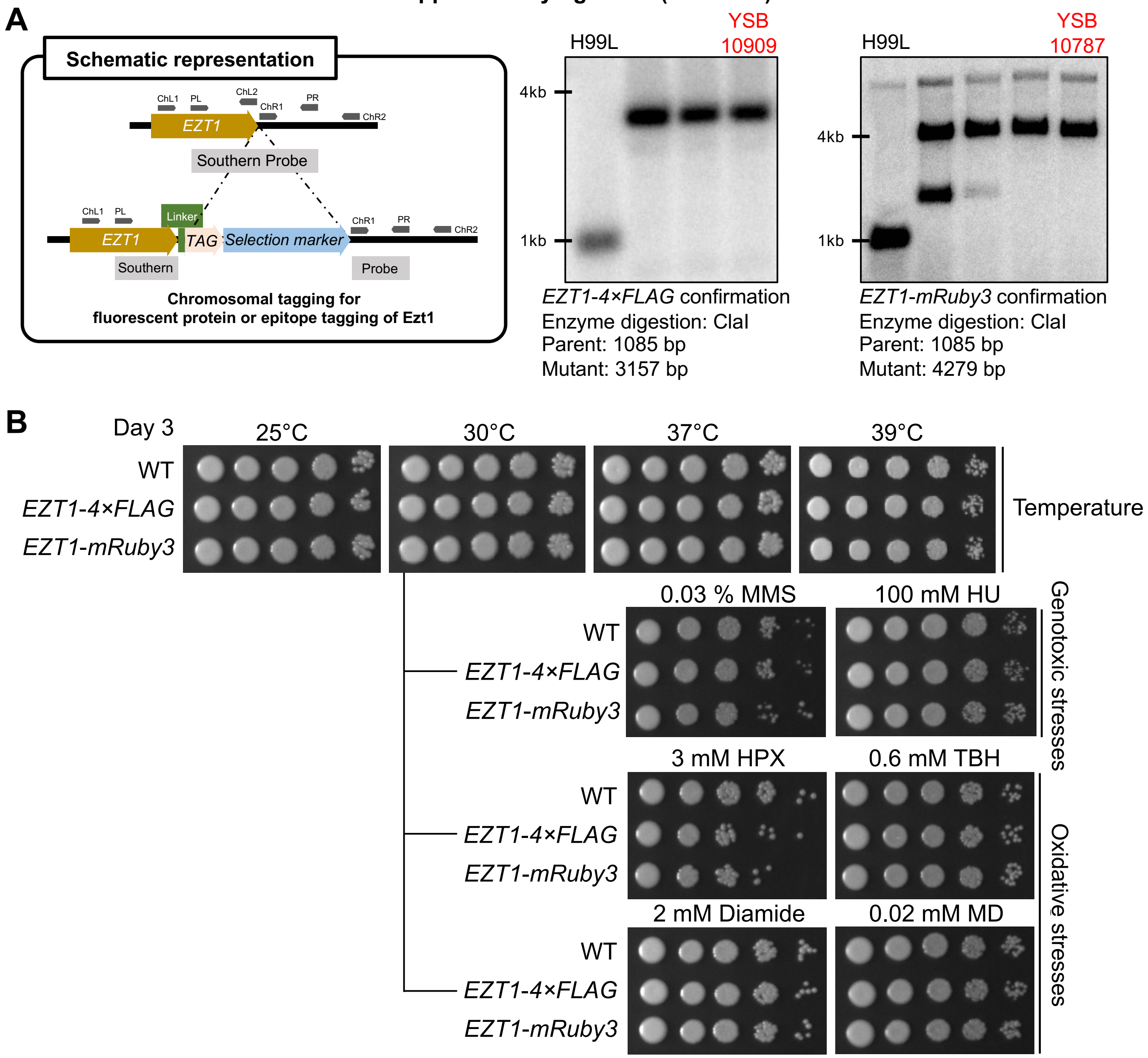


*Continued*

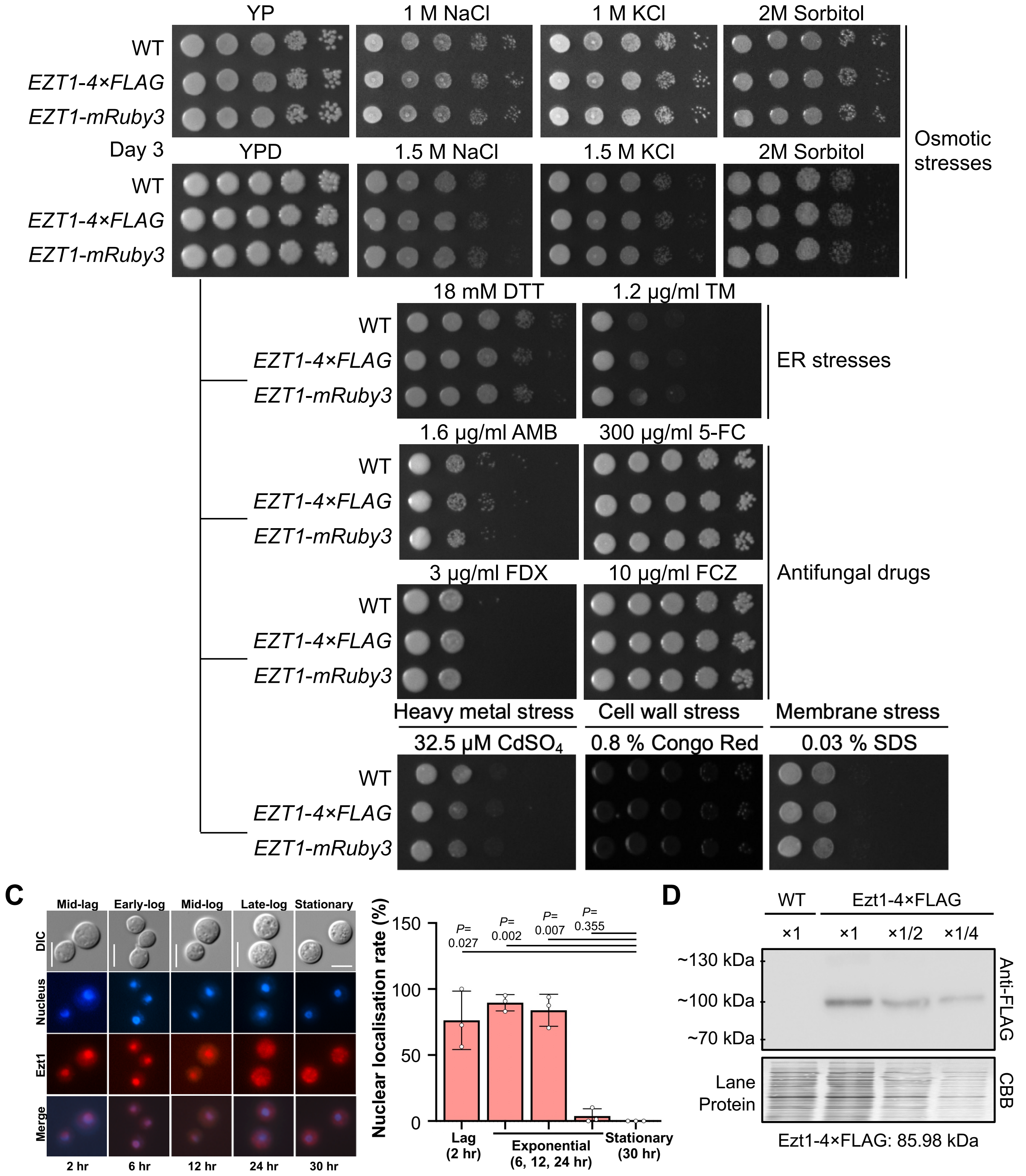


**Supplementary figure 10. Chromosomal tagging of Ezt1.** (A) Schematic representation for the construction of Ezt1-4×FLAG and Ezt1-mRuby3 strains is shown in box. Correct genotypes of the strains were confirmed through Southern blot analysis with the restriction enzyme ClaI and *EZT1*-specific probe. (B) The functional validation of Ezt1 proteins tagged with FLAG and mRuby3 was conducted through phenotypic analysis. The wild-type and the chromosomally tagged strains were cultured overnight at 30°C in liquid YPD medium, serially diluted (1 to $10^4$), and spotted onto the YPD plates (3 µl) containing the indicated amount of stress or antifungal agents. Plates were incubated at 30°C (or designated temperature) and photographed after 3 days. (C) Cellular localisation of Ezt1. Fluorescence microscopy showed nuclear localisation of Ezt1-mRuby3 during the exponential growth phase, with cytoplasmic translocation occurring from the late-log to stationary phases (scale bar: 5 µm). The 6 h and 30h time points are also presented in Fig. 4d as representative of Ezt1 nuclear localisation during exponential phase and stationary phase. Data are represented as mean ± SD. Statistical significance of difference was determined by one-sample *t*-test using Prism 11.0 (*P* values are stated above). (D) Detection of Ezt1-4×FLAG by Western blotting. Wild-type and Ezt1-4×FLAG strains were cultured in YPD at 30°C, diluted to $OD_{600}$ = 0.2, and grown to 0.8. Cells were washed once with water, frozen in liquid nitrogen, and lysed in 650 µL of cracking buffer (0.42 M NaOH and 1.9% β-mercaptoethanol) for 10 min on ice. Proteins were precipitated with 150 µL of 50% trichloroacetic acid, centrifuged at 14,000×g for 10 min at 4°C, washed with 500 µL ice-cold acetone, and resuspended in 150 µL loading buffer containing 40 mM Tris-HCl (pH 6.8), 5% SDS, 100 mM NaEDTA, and 8.3 M urea. Samples were incubated at 42°C for 10 min and centrifuged at 14,000×g for 2 min. Whole-cell lysates were quantified using in-house 10% stain-free gels containing 0.5% (vol/vol) 2,2,2-trichloroethanol and Image Lab software (Bio-rad). Comparable protein amounts were separated on 10% Novex Tris-glycine gel, transferred to PVDF membrane, and blocked with 10% skim milk in TBS-T containing 10 mM Tris-HCl (pH 7.5), 150 mM NaCl, and 0.05% Tween-20. Ezt1-4×FLAG was detected using anti-FLAG antibody (Sigma-Aldrich, F3165) and Clarity ECL HRP substrate (Bio-Rad). Lane protein was visualised with Coomassie brilliant blue (CBB) R-250 staining of the membrane.

**Supplementary figure 11 (Lee et al.)**

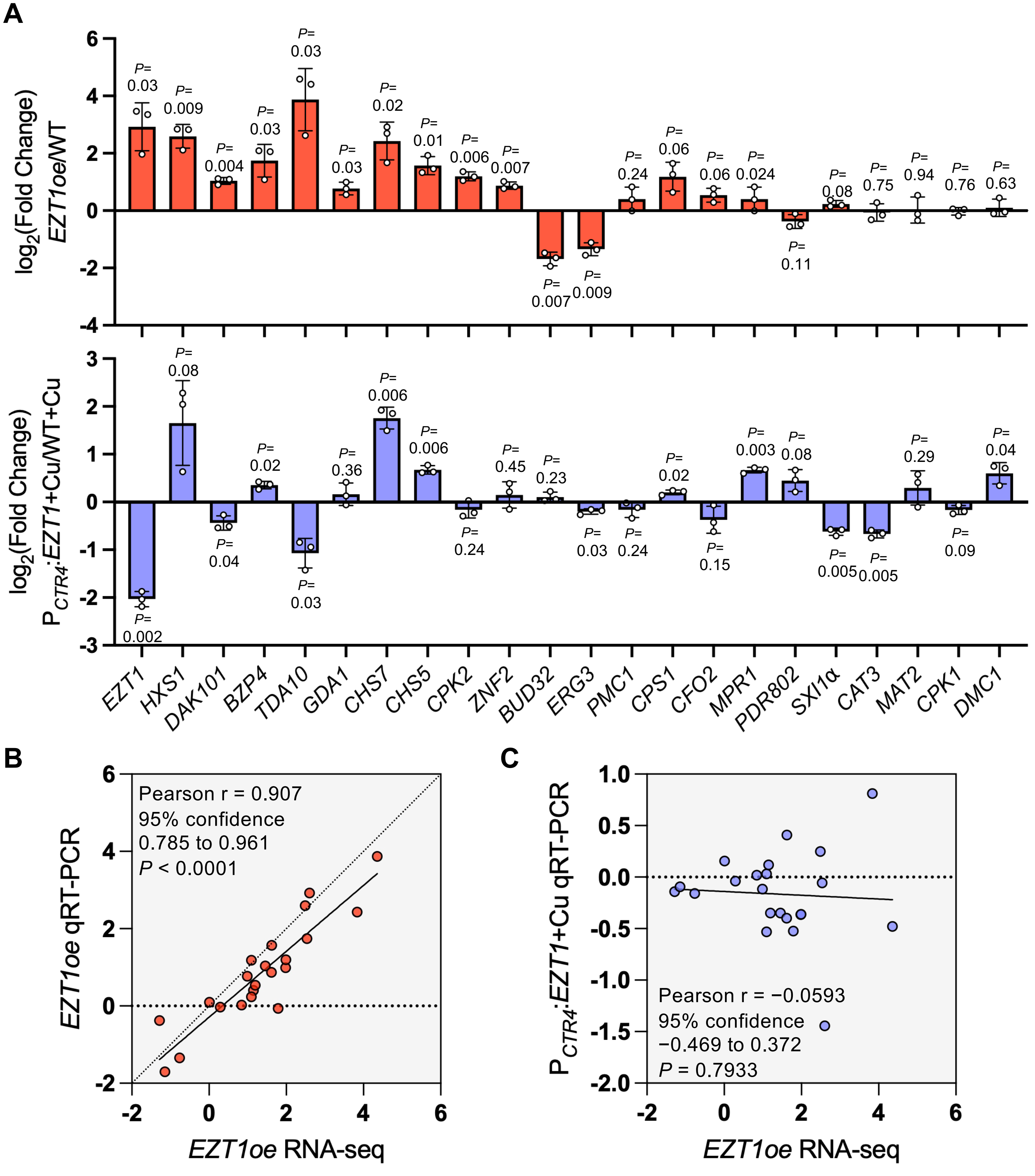


**Supplementary figure 11. Independent qRT-PCR assessment of *EZT1*-responsive genes following *EZT1* overexpression and Cu-mediated partial repression**. (A) qRT-PCR analysis of 22 transcripts in the *EZT1oe* strain relative to the wild-type (upper panel) and qRT-PCR analysis of the same 22 transcripts in $P_{CTR4}$:*EZT1* relative to the wild-type in the presence of $CuSO_4$ (bottom panel). The panel comprised 15 differentially expressed genes with promoter-proximal Ezt1 binding, four differentially expressed genes without detectable promoter binding, *EZT1* as an expression control, and two genes (*CPK1* and *DMC1*) meeting neither criterion in *EZT1oe* RNA-seq results. Values represent log2 fold changes in *EZT1oe* relative to wild type (upper panel), and in $P_{CTR4}$:*EZT1* + Cu relative to WT + Cu (bottom panel). Individual points represent independent biological replicates, and horizontal lines represent the mean ± SD (n=3). Statistical significance was assessed using two-tailed one-sample *t*-test against a theoretical log2 fold change of 0, and *P* values are shown. (B) Correlation between RNA-seq- and qRT-PCR-derived log2 fold changes for the 22 transcripts in *EZT1oe* relative to WT. (C) Correlation between RNA-seq-derived log2 fold changes in *EZT1oe* relative to WT and the qRT-PCR-derived log2 fold changes in $P_{CTR4}$:*EZT1* + Cu relative to WT + Cu. For (C), each point represents one transcript, solid lines indicate least-square linear regression. Pearson's r correlation coefficients, two-sided *P* values and 95% confidence intervals are shown. Statistical analyses were performed using Prism 11.0.

**Supplementary figure 12 (Lee et al.)**

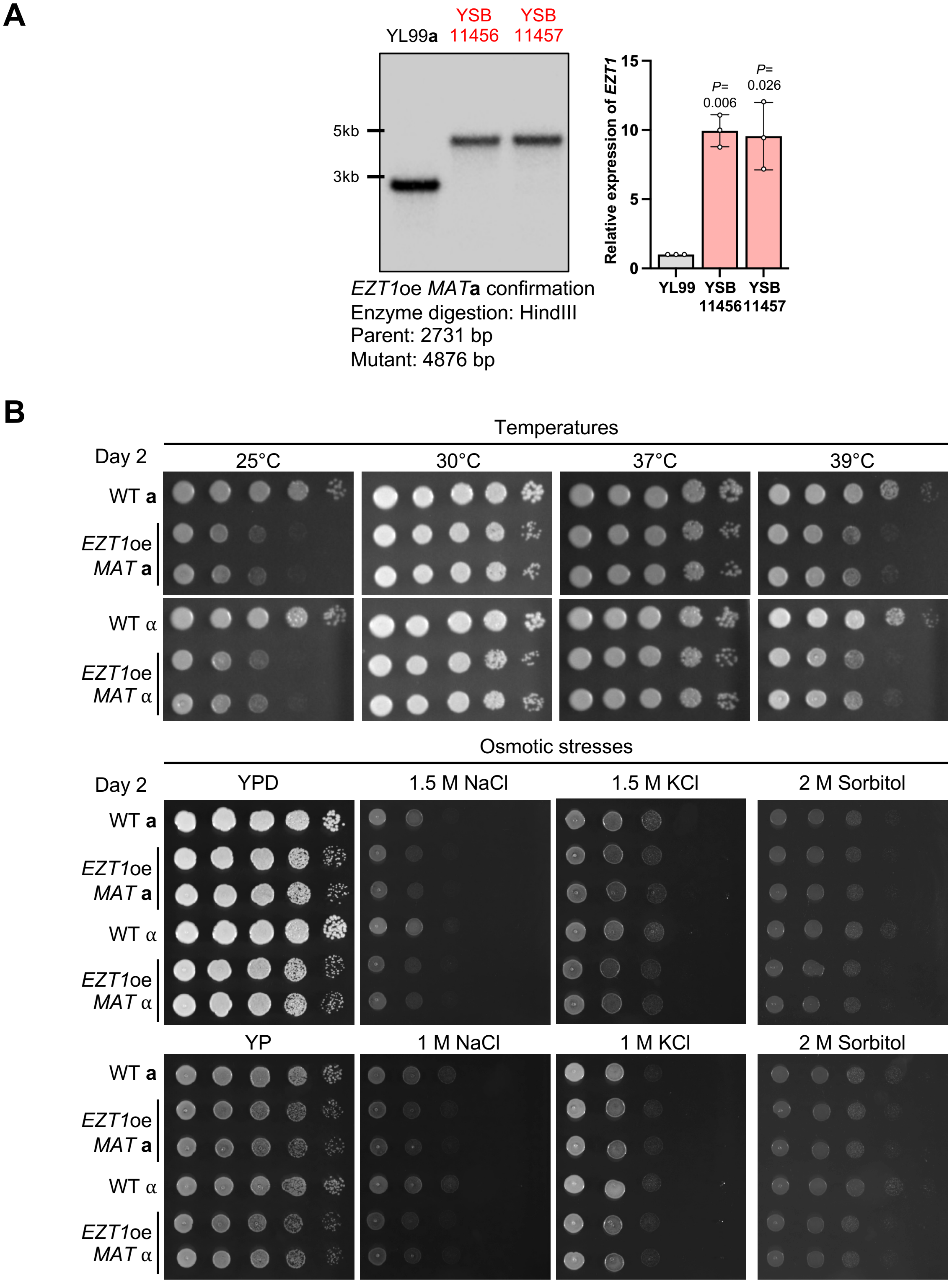


***Continued***

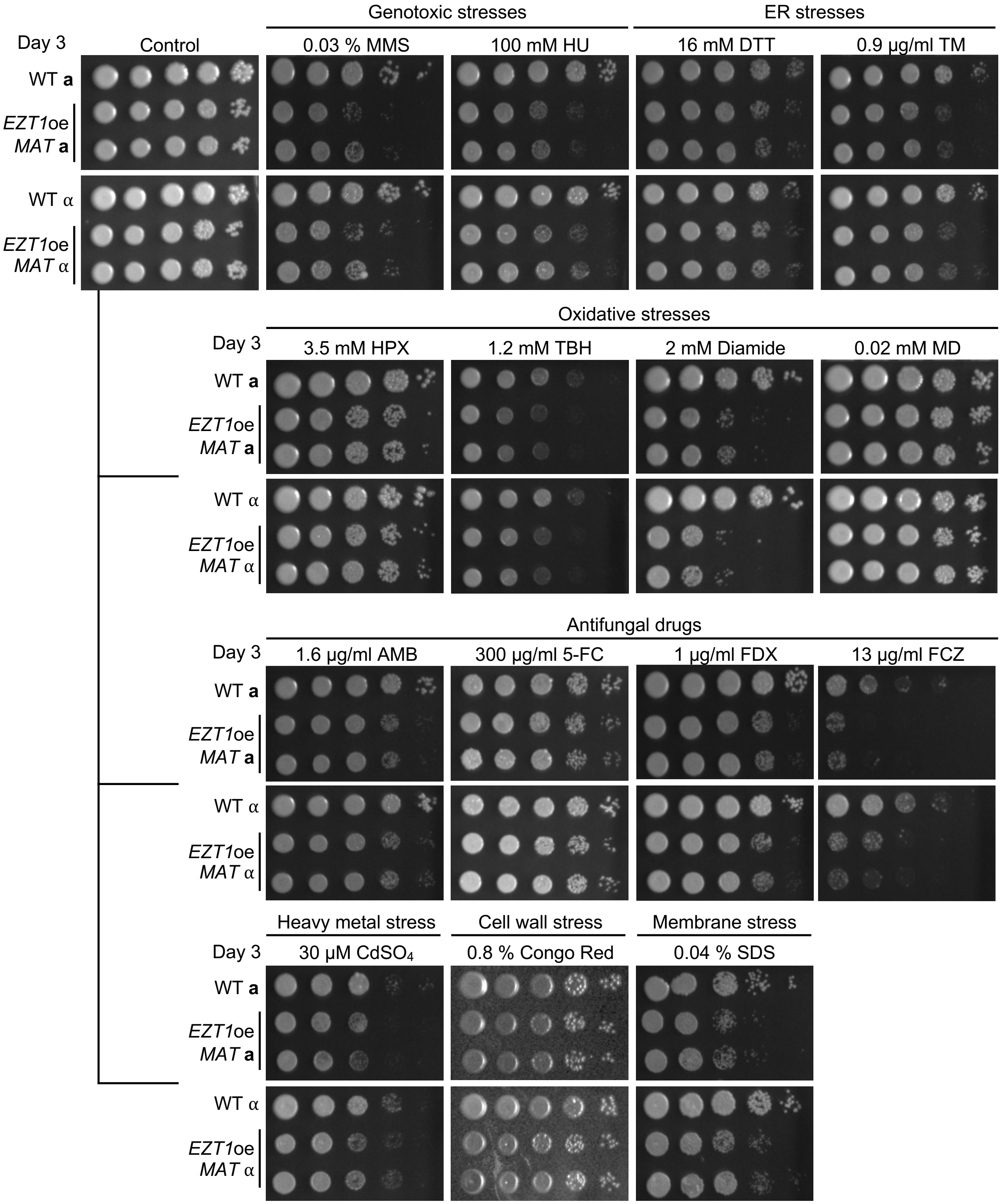


**Supplementary figure 12. The *MAT*a *EZT1*oe strain phenocopied the *MAT*α *EZT1*oe strain**. (A) The correct genotype of the *MAT*a YL99 *EZT1*oe strains was confirmed through Southern blot analysis with the restriction enzyme HindIII and *EZT1*-specific probe. The overexpression of the targeted genes was confirmed with qRT-PCR using the identical procedure as for the expression validation of the essential TF overexpression strains. Data are represented as mean ± SD. Statistical significances of differences between wild-type and the overexpression strain were determined by one-sample *t*-test using Prism 11.0 (*P* values are stated above). (B) The phenotypic analysis confirmed that *EZT1*oe *MAT* **a** (YSB11456 and YSB11457) in WT **a** (YL99 *MAT* **a**) and *EZT1*oe *MAT* α (YSB7110 and YSB7111) in H99L *MAT* α (WT α) exhibit the same phenotype. The wild-types and the overexpression strains from both mating types were cultured overnight at 30°C in liquid YPD medium, serially diluted (1 to $10^4$), and spotted onto the YPD plates (3 μl) containing the indicated amount of stress or antifungal agents. Plates were incubated at 30°C (or designated temperature) and photographed after designated days.

### Supplementary figure 13 (Lee et al.)

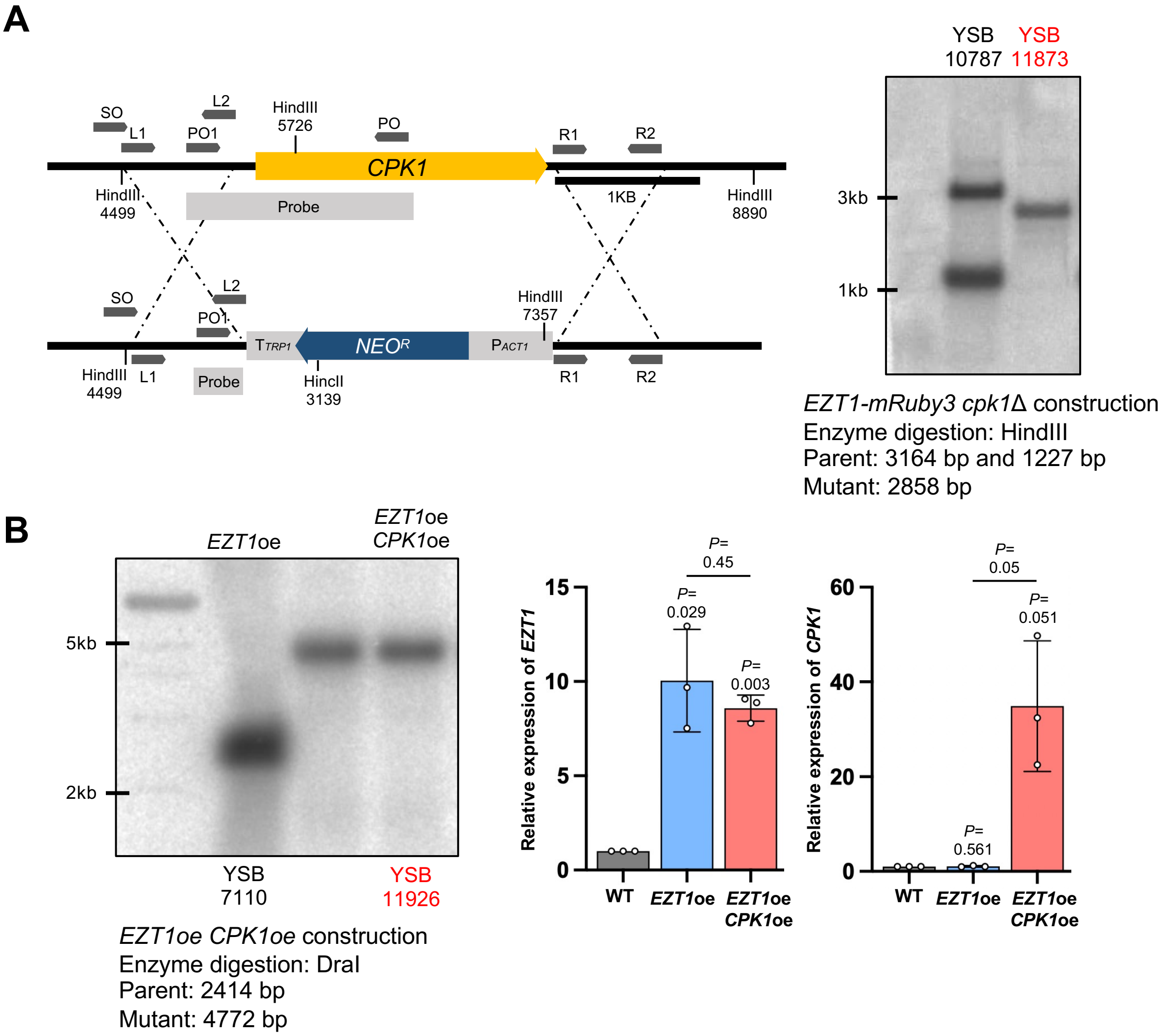


**Supplementary figure 13. Construction of strains to investigate the relationship between *EZT1* and *CPK1*.** (A) Strategy and confirmation of the constructed *EZT1*-mRuby3 *cpk1*Δ strain. Southern blot analysis confirmed the genotype of the constructed mutant with the restriction enzyme HindIII digestion. (B) Correct genotype of the *EZT1* and *CPK1* double overexpression strain was confirmed through Southern blot analysis with the restriction enzyme DraI and *CPK1*-specific probe. The overexpression of the targeted genes was confirmed with qRT-PCR using the identical procedure as for the expression validation of the essential TF overexpression strains. Data are represented as mean ± SD. Statistical significances of differences were determined by one-sample *t*-test for the overexpression and unpaired *t*-test between *EZT1*oe (YSB7110) and *EZT1*oe *CPK1*oe (YSB11926) using Prism 11.0 (*P* values are indicated above).

### Supplementary figure 14 (Lee et al.)

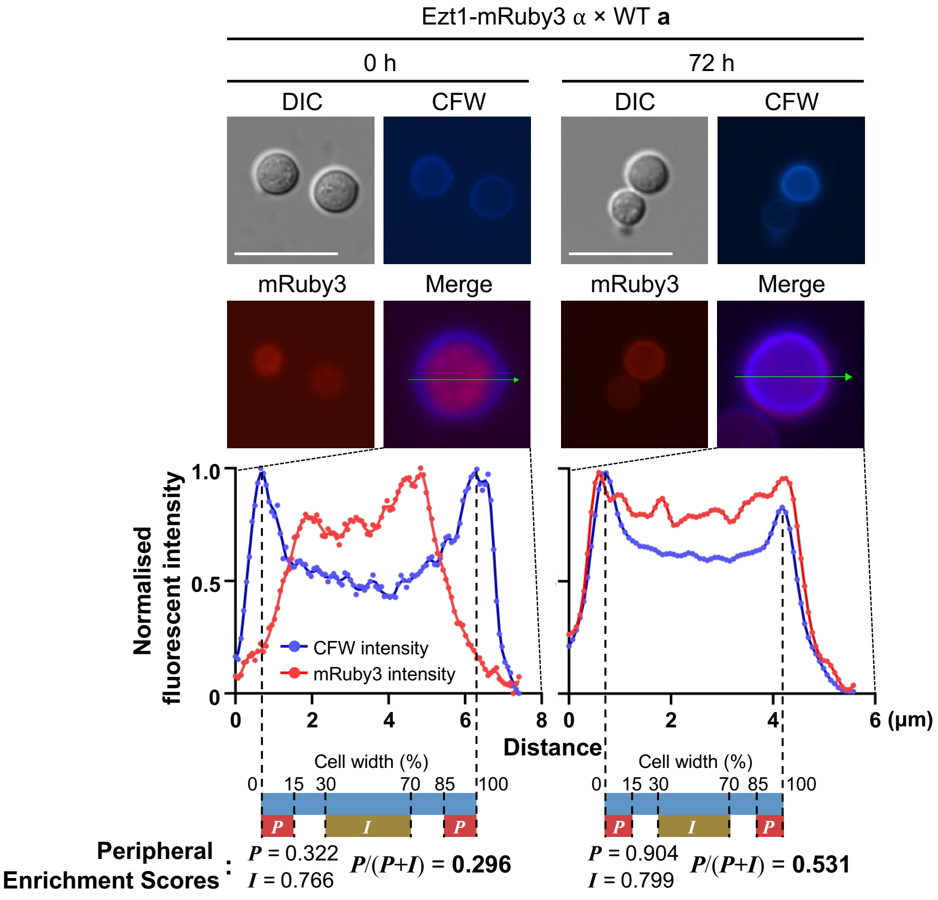


**Supplementary figure 14. Example for the quantification of Ezt1-mRuby3 peripheral enrichment during mating.** Representative images shown here are those presented in Fig. 5C. Ezt1-mRuby3 and Ezt1-mRuby3 *cpk1Δ* cells were stained with calcofluor white (CFW) to define the cell-wall boundaries (scale bar: 10 µm). For each cell, a line was drawn through the cell centre and across the two opposing CFW-stained cell walls, and fluorescence intensity profiles of CFW and mRuby3 were obtained along the line with Fiji/ImageJ. Cell-specific background intensity was subtracted separately from the mRuby3 and CFW fluorescence profiles. The two highest CFW peaks flanking the cell centre were defined as the cell-wall boundaries, and the region between the boundaries was normalised to 0-100% of the cell width. Peripheral intensity (*P*) was defined as the mean normalised mRuby3 fluorescence intensity within the 0-15% and 85-100% regions (red boxes), whereas interior intensity (*I*) was defined as the mean normalised mRuby3 fluorescence intensity within the 30–70% region (brown boxes). Peripheral enrichment score was calculated as *P*/(*P*+*I*).

### Supplementary figure 15 (Lee et al.)

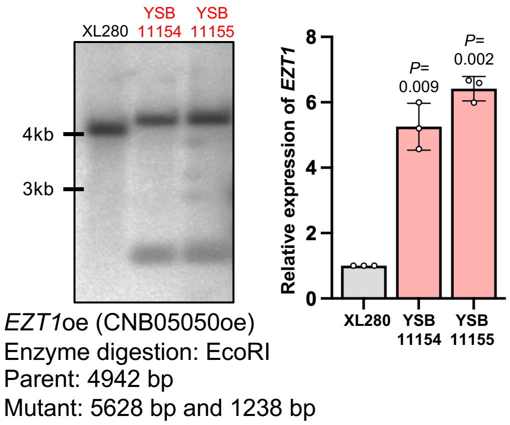


**Supplementary figure 15. Construction of *EZT1* overexpression strains in XL280.** (A) Correct genotype of the *Cryptococcus deneoformans EZT1* (CNB05050) overexpression strains was confirmed through Southern blot analysis with the restriction enzyme EcoRI and *EZT1*-specific probe. The overexpression of the targeted genes was confirmed with qRT-PCR using the identical procedure as for the expression validation of the essential TF overexpression strains. Data are represented as mean ± SD. Statistical significances of differences between wild-type and the overexpression strains were determined by one-sample *t*-test compared to the wild-type for the overexpression strains using Prism 10.0 (*P* values are indicated above)

### Supplementary figure 16 (Lee et al.)

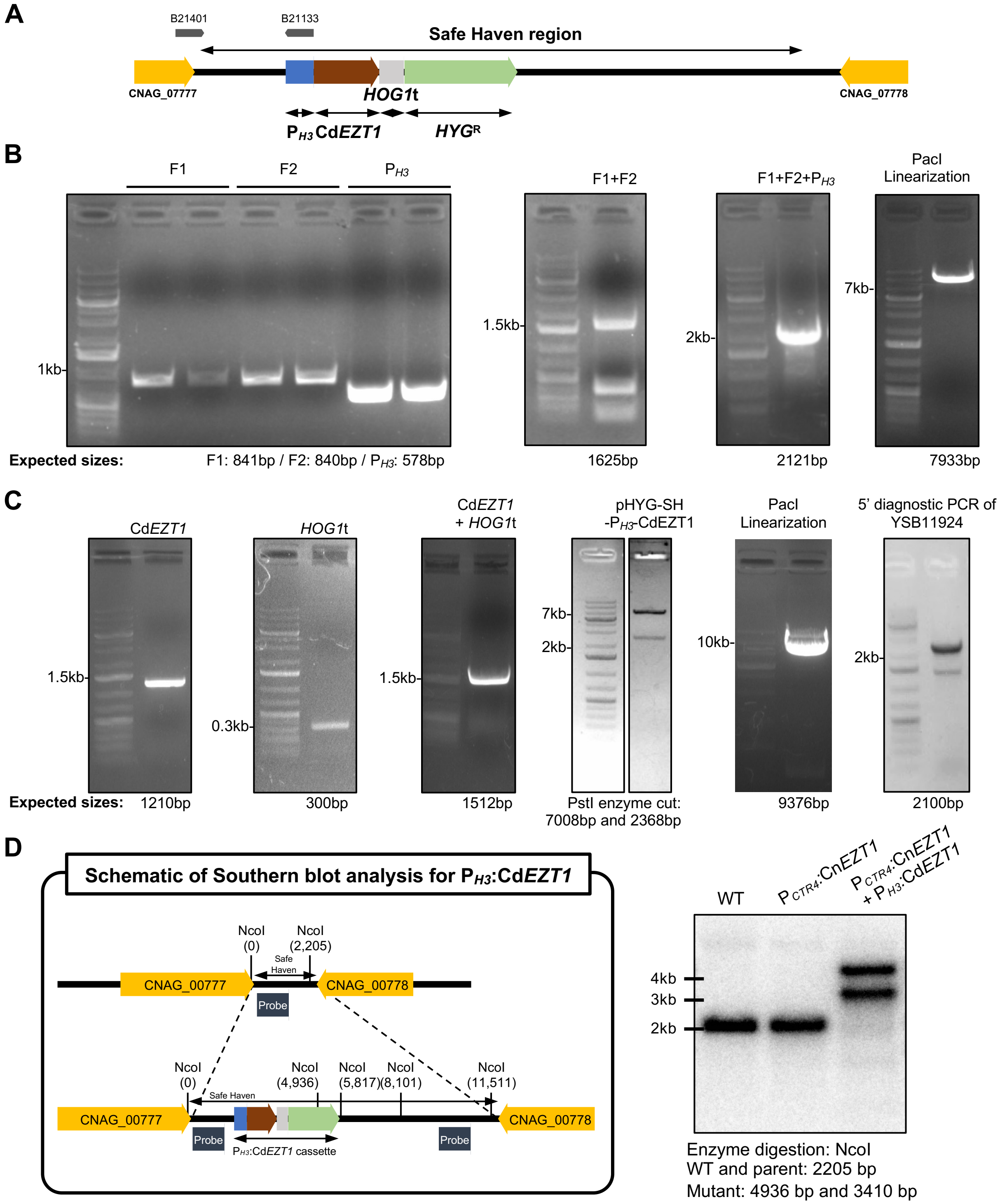


**Supplementary figure 16. Heterologous expression of *Cryptococcus depauperatus EZT1*.** (A) Schematic strategy for constructing a strain heterologously overexpressing *Cryptococcus depauperatus EZT1* (Cd*EZT1*). (B) To construct the pHYG-SH(Safe Haven)-$P_{H3}$-Cd*EZT1* plasmid, the intergenic region between CNAG_00777 and CNAG_00778 was divided into two fragments (F1 and F2) and amplified by PCR. These fragments were then fused through overlap PCR to introduce AscI, BaeI, and PacI restriction sites in the middle of the intergenic region. Subsequently, the *H3* promoter ($P_{H3}$) was amplified and integrated with the intergenic region via overlap PCR. This fused fragment was ligated with pHYG using Gibson assembly (New England Biolabs, USA), resulting in the pHYG-SH-$P_{H3}$ plasmid (confirmed with PacI linearization). (C) To introduce Cd*EZT1*, the L203_03018 gene from *C. depauperatus* CBS7841 was PCR-amplified and assembled into pHYG-SH-$P_{H3}$ using Gibson assembly, yielding the final pHYG-SH-$P_{H3}$ -Cd*EZT1* plasmid (confirmed with PstI enzyme digestion). The plasmid was linearized by PacI digestion, and the purified cassette was subsequently introduced into the $P_{CTR4}$:*EZT1* strain. Transformant (YSB11924) is confirmed with 5' junctional diagnostic PCR. (D) Schematic of the Southern blotting strategy for the *CdEZT1* heterologous expression strain (left) and the corresponding Southern blot result obtained using NcoI-digested genomic DNA (right). The genetic elements in the schematic are color-coded as in panel (A), with matching colors indicating identical DNA fragments. Correct integration of the $P_{H3}$:*CdEZT1* construct into the conditional *EZT1* expression strain ($P_{CTR4}$:Cn*EZT1*) was confirmed by comparing the WT strain (H99L), the parental strain $P_{CTR4}$:*EZT1* (YSB9280), and the $P_{CTR4}$:*EZT1* + $P_{H3}$:Cd*EZT1* strain (YSB11924).
