## Supplementary material for "Systematic Profiling of Essential Fungal Transcriptional Regulators Uncovers Ezt1 as a Central Pathobiological and Morphogenic Regulator in *Cryptococcus neoformans*": Description of Addtional Supplementary Informations

**Description of Additional Supplementary Files**

File names:

1. Supplementary Data 1. Essentiality of the 17 putative essential transcriptional regulators
2. Supplementary Data 2. BLAST analysis and matrix of the 17 putative essential transcriptional regulators.
3. Supplementary Data 3. RNA sequencing data for transcriptome analysis of *EZT1*, *EZT2*, or *CBF1* overexpression strains compared with the wild-type strain.
4. Supplementary Data 4. Chromatin-immunoprecipitation sequencing (ChIP-seq) results for the Ezt1-4×FLAG strain.
5. Supplementary Data 5. Primers used in this study.
6. Supplementary Data 6. Strains used in this study.
7. Supplementary Data 7. *P* values for expression of mating-related genes in bilateral *EZT1* overexpression and wild-type mating.

File description:

1. Essentiality analysis of 17 putative essential transcriptional regulators, including functional classification, essentiality in other fungal species, conditional expression phenotypes, and meiotic spore analysis results.
2. Comparative BLAST analysis of 17 putative essential transcriptional regulators, including BLASTp and reciprocal BLAST results and a comparative fungal BLAST matrix.
3. RNA-seq analysis of *EZT1*, *EZT2*, and *CBF1* overexpression strains compared with the wild-type strain, including complete expression datasets and lists of significantly upregulated and downregulated genes.
4. ChIP-seq analysis of Ezt1-4×FLAG, including complete and high-confidence promoter-binding peaks and genes directly regulated by Ezt1 based on integration with transcriptomic data.
5. Sequences and descriptions of all primers used in this study.
6. Genotypes, parental strain information, references, and corresponding figures for all fungal strains used in this study.
7. *P* values from statistical comparisons of mating-related gene expression between bilateral *EZT1* overexpression and wild-type mating across the indicated time points, corresponding to the data presented in Main Figure 6E.
